# Proteomizer: Leveraging the Transcriptome-Proteome Mismatch to Infer Novel Gene Regulatory Relations

**DOI:** 10.1101/2025.06.22.660946

**Authors:** Giulio Deangeli, Maria Grazia Spillantini, Pietro Liò

**Affiliations:** University of Cambridge, Department of Clinical Neurosciences, Clifford Allbutt Building, Hills Road, CB2 0HA Cambridge, UK; University of Cambridge, Department of Computer Science and Technology, William Gates Building, 15 J. J. Thomson Ave, CB3 0FD Cambridge, UK

## Abstract

The correlation between transcriptomic (Tx) and proteomic (Px) profiles remains modest, typically around *r* = 0.5 across genes and *r* = 0.3 across samples, limiting the utility of transcriptomic data as a proxy for protein abundance. To address this, we introduce Proteomizer, a deep learning platform designed to infer a sample’s Px landscape from its Tx and miRNomic (Mx) profiles. Trained on 8,613 matched Tx-Mx-Px samples from TCGA and CPTAC, Proteomizer achieved a Tx-Px correlation of *r* = 0.68, representing the highest performance reported to date for this task. We further developed a Monte Carlo simulation framework to evaluate the impact of proteomization on differential expression analysis. Proteomizer substantially improved the accuracy of differential gene expression detection, with p-value precision increasing by up to 62-fold, and by as much as six orders of magnitude for a subset of genes enriched in mitochondrial and ribosomal functions. However, performance gains did not generalize to unseen tissue types or datasets generated using different protocols. Finally, we applied explainable AI (XAI) techniques to identify regulatory relations contributing to Tx-Px discrepancies. Our predictions from 100 highly annotated genes were cross-compared against by a literature-based biological knowledge graph of 322 million annotations: our explainers achieved a ROC-AUC of 0.74 in predicting miRNA-gene downregulation interactions. To our knowledge, this is the first study to systematically evaluate the biological relevance, limitations, and interpretability of proteomization models, establishing Proteomizer as a state-of-the-art tool for multiomic integration and hypothesis generation.

**Graphical Abstract:** 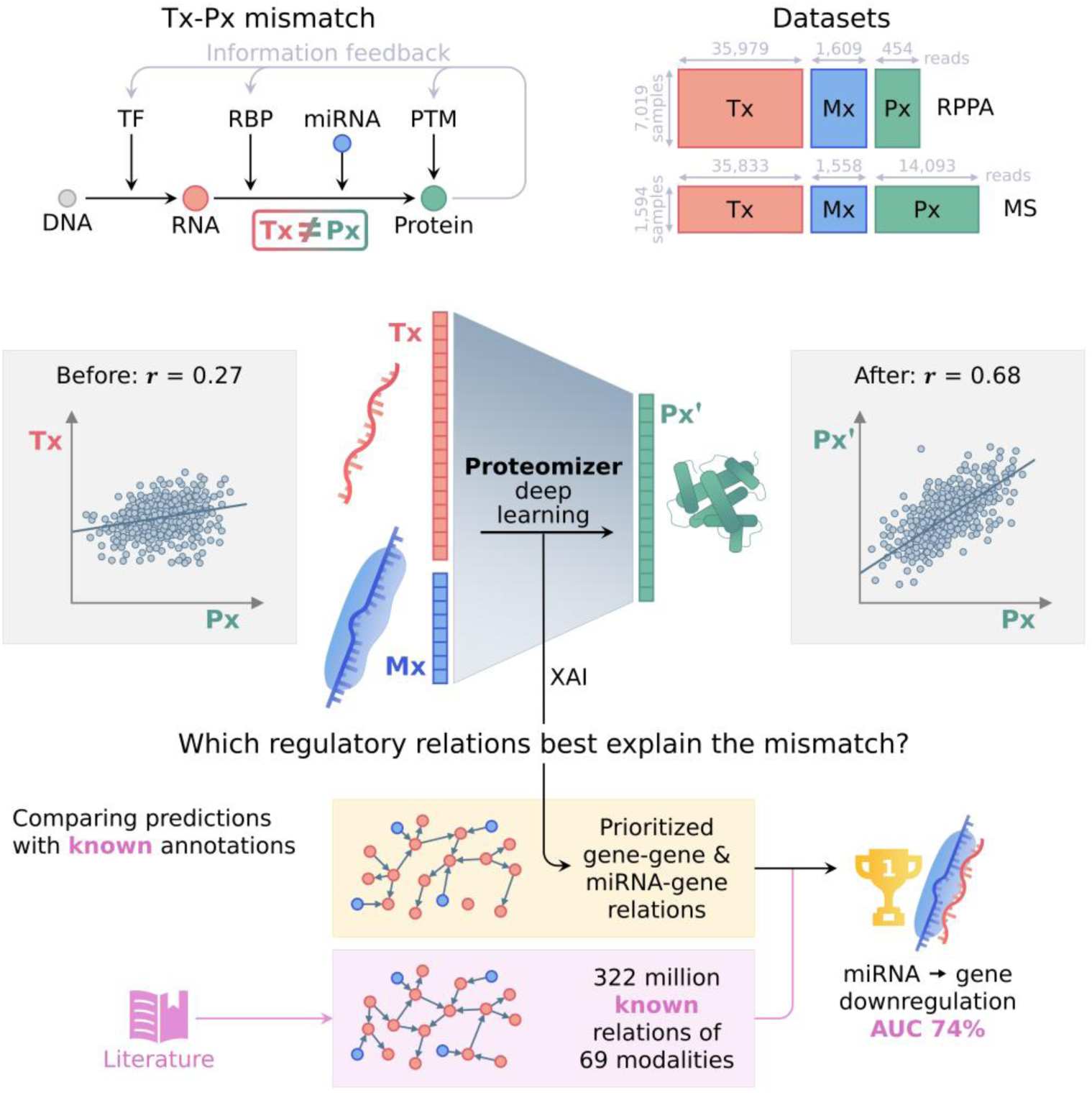

miRNA: microRNA; Mx: miRNome; MS: mass spectrometry; PTM: post-translational modifications; Px: proteome; Px’: predicted proteome; *r*: Pearson correlation coefficient; RBP: RNA binding proteins; ROC-AUC: area under the receiver operating characteristic curve; RPPA: reverse-phase protein array; TF: transcription factors; Tx: transcriptome; XAI: explainable artificial intelligence.

## Introduction

Through the early 2000s, “omic” techniques have become universally widespread in molecular biology, allowing investigators to collect big data from biological models. When studying a condition (say, diseased versus healthy, perturbed versus unperturbed, etc.) using omics, it is common practice to measure gene expression differences in terms of messenger RNA (mRNA), that is, by transcriptomics. However, the investigators’ interest usually lies in the proteins (not the RNAs), for it is the proteins that carry out most of the biological functions.^1,2^ Proteins act as drug targets, carry out pathogenetic mechanisms, work as biomarkers, and have the strongest association with disease severity.^3,4^ Still, RNA levels are habitually employed as a proxy for protein concentration for better convenience: transcriptomics (Tx) is significantly cheaper than proteomics (Px),^5,6^ more adaptable (e.g., can be performed at single-cell resolution^7^), and it is operationally less complex, resulting in greater availability.

By using Tx as a surrogate of Px, researchers are implicitly assuming that genes’ RNA levels approximately mirror those of their cognate proteins. This assumption has been scrutinized since the early 2000’s, leading to the worrying observation that, in most cases, mRNA levels are mismatched from those of their cognate proteins, posing a threat to the interpretability of Tx.^8^ The Tx-Px mismatch is usually quantified by the Pearson *r* and the Spearman *ρ* correlation coefficients. Because Tx and Px data are usually Poisson-like distributed, Spearman *ρ* is theoretically preferrable, as it does not assume normal distribution.^9,10^ However, in practical terms, their values closely match.^11^ Tx-Px correlation can be measured in two distinct fashions. Correlation “across the genes” (i.e., within samples) measures how the Tx expression of different genes is matched by their respective Px values; it is usually applied when only *n* = 1 sample is available, as it used to be the norm until the mid-2010’s. (ii) Conversely, correlation “across the samples” (i.e., within genes) measures how the Tx expression of the same gene in many samples is matched by that gene’s Px values, on average, for all genes in the datasets; this method is by far the most biologically relevant, as it better captures the endpoint of Tx, which is the pair-wise comparison between biologically different sample groups.^11^

Since as early as 1999,^12^ experimentalists have quantified Tx-Px mismatch across a wide array of techniques, species and stimuli (Table 1). The reported correlations have been consistently very modest, with *r* values around 0.4-0.7 across genes, and 0.1-0.5 across samples. For example, in a large dataset of glioblastoma samples, approximately one third of genes showed negative Spearman correlation across samples.^13^ Even in optimal experimental conditions, such as the realization of a mouse proteomic atlas, cross-gene correlation was measured at a mere *r* = 0.43.^14^

**Table 1.**
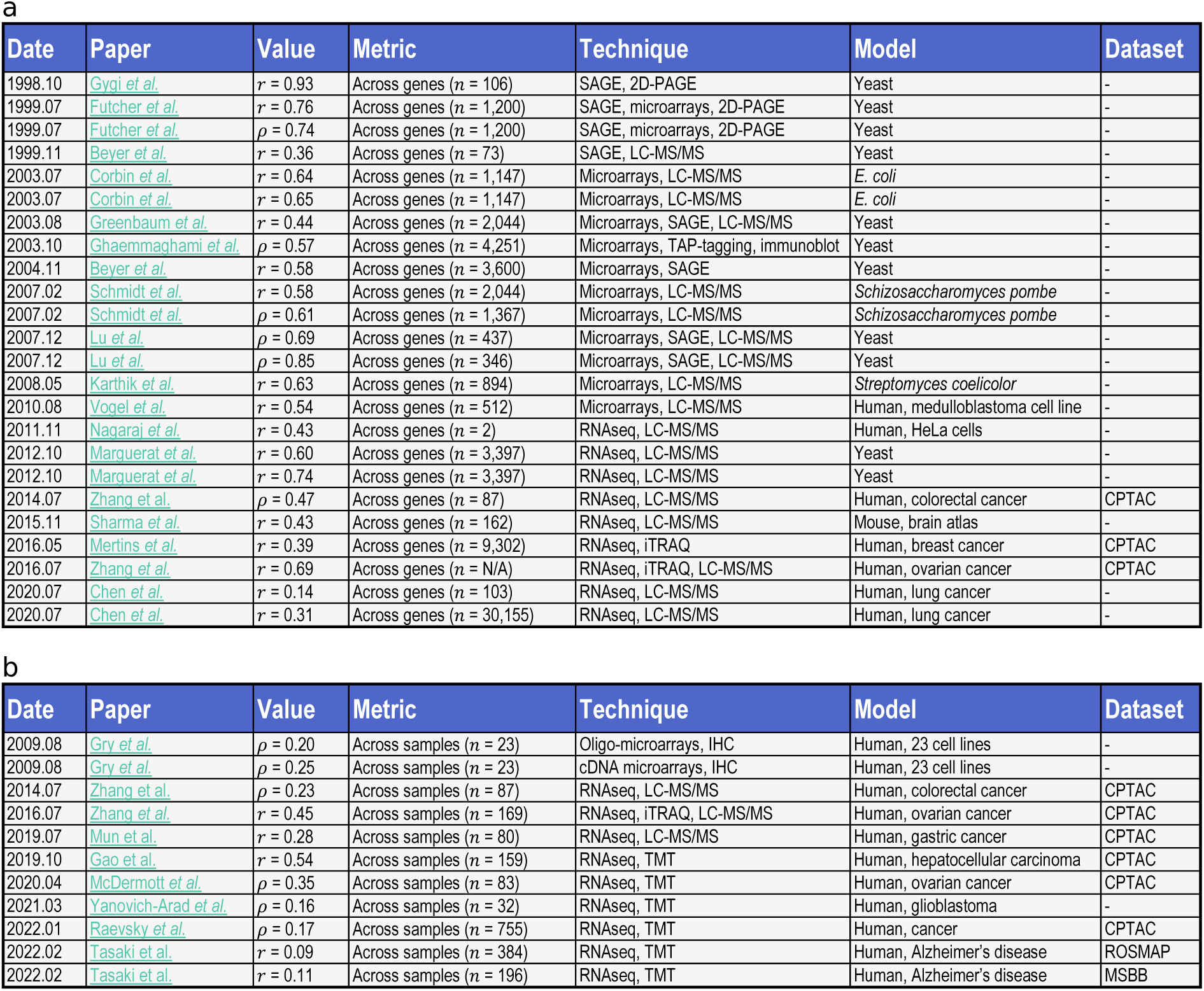
Literature overview of transcriptomic-proteomic mismatch, quantified by correlation metrics across genes and across samples. The table offers a birds-eye overview of the last 25 years of literature on the mismatch between transcriptomics (Tx) and proteomics (Px). It reports some of the most notable examples of papers that measured the mismatch, either by the Pearson correlation coefficient *r*, or by the Spearman correlation coefficient *ρ*. While it does not have any ambition of completeness— as it would be far beyond the scope of the present research—it makes it easy to spot general patterns in researchers’ preference along the years, in terms of the reported values, measurement strategy and biochemical technique. Two approaches are possible to quantify the mismatch: (a) across the genes, i.e., within the samples; (b) across the samples, i.e., within the genes. The first approach used to be popular before 2010, as it can be performed with *n* = 1 sample size, which was common before the popularization of proteomics in the 2010’s. Conversely, the latter approach is more “biological”, as it compares the same gene across multiple samples in an analogous way to differential expression analysis (which is usually the biological endpoint of Tx), and then, it averages out the results for all genes. Of note, it cannot be applied using tandem mass tag (TMT) mass spectrometry, as TMT quantifies proteins as a ratio to a provided common reference sample; as a result, each protein is read in a different scale from that of the other proteins. Papers are sorted by the date in which they were published. The “Model” column highlights what model organism what used, and in what condition. The “Dataset” column reports whether data were sourced from any of the established repositories, the most notable being: the Clinical Proteomic Tumor Analysis Consortium (CPTAC), the Religious Orders Study/Memory and Aging Project (ROSMAP) and the Mount Sinai Brain Bank study (MSBB). 2D-PAGE: 2-dimensional polyacrylamide gel electrophoresis; cDNA: complementary DNA; iTRAQ: isobaric tags for relative and absolute quantitation mass spectrometry; LC-MS/MS: liquid chromatography-tandem mass spectrometry; RNAseq: RNA sequencing; SAGE: serial analysis of gene expression; TAP-tagging: tandem affinity purification tagging; TMT: tandem mass tags mass spectrometry.

Indeed, the mismatch can be justified on three different levels. (1) From an evolutionary standpoint, stringent regulation of gene product concentrations is energetically costly. As result, evolutionary pressures have favored a hierarchical prioritization: tighter control is exerted over proteins than RNAs,^15,16^ and among proteins, those with essential biological functions are more stringently regulated than those with lesser roles.^17,18^ Specifically, post-translational regulation is inherently wasteful (as it often involves the inactivation or degradation of already synthesized proteins), while it offers the advantage of rapid response times, making it a prerogative of critical cellular processes.^19–21^ (2) From a biological standpoint, several mechanisms contribute to the Tx-Px decoupling. RNA interference, mediated by microRNAs (miRNAs) and other non-coding RNAs, can variably suppress translation, promote mRNA degradation,^22^ or in some contexts even enhance protein expression^23^; these effects are irregularly mirrored at Tx level, leading to complex regulatory patterns.^24^ Additionally, translational efficiency can be proactively modulated by a variety of factors, such as the optional use of internal ribosome entry sites (IRES)^25^ or upstream open reading frames (uORFs),^26^ ribosome density,^27,28^ codon bias^29^ and amino acid bias^30^: all of these can be subject to condition-dependent regulation. Protein turnover is also strictly regulated, by means of the ubiquitin-proteasome and autophagy pathways.^31,32^ Furthermore, in the case of proteins with very long half-lives, their mRNA levels should conceptually mirror the production rate of their cognate protein species, rather than the protein levels themselves. (3) Finally, technical justifications used to play a major role before 2010, when Tx and Px were commonly measured by DNA microarrays and reverse-phase protein arrays (RPPA), respectively. However, recent advances in RNA sequencing and tandem mass tag (TMT)-based mass spectrometry have significantly improved the accuracy and resolution of both measurements, rendering technical artifacts a comparatively minor contributor to the Tx-Px mismatch.^2,33^

The increasing accessibility of multiomic data over the past two decades has enabled investigators to apply machine learning (ML) to impute the Px landscape of samples, based on their Tx landscapes, sometimes in combination with other omic data. To indicate this kind of ML task, we coined the neologism of “proteomizing” the transcriptome—a “proteomization” task. The implicit assumption is that, because (most) proteins are kept at controlled amounts in a cell, their concentration information must be backpropagated into gene expression, and will therefore be mirrored somewhere in the Tx. Or equivalently, we postulate the existence of a cell (or tissue) “state”, which is reflected in both its Px vector and its Tx vector.^3^ Thereby, ML plays the role of an isomorphism from the Tx state to the Px state—analogously to how language models may map the embedding space of one language into that of another language.^34^ ML-based proteomization would allow to correct for post-transcriptional regulation, effectively transforming the affordable and widely available Tx, into the biologically accurate yet costly Px.

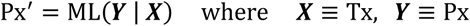

Proteomization literature goes back to as early as 2007, when Tuller *et al*. combined all the yeast datasets available at the time (two Tx and four Px ones) with 32 static gene features, and deployed linear models and support vector machines, improving cross-gene correlation from *ρ* = 0.55 to 0.76.^35^

In 2005, The Cancer Genome Atlas (TCGA) was established as a collaborative effort, led by led by the National Cancer Institute (NCI) and the National Human Genome Research Institute (NHGRI).^36^ The project aimed at collecting full multiomic profiling for thousands of primary cancers and matched normal samples. Characterization included genomic panel of mutations, Tx profiling (first by microarrays, later on by RNA sequencing), Px (by RPPA), miRNomics, but also metabolomics, epigenomics and more.^37^ In 2011, a sister project was announced called Clinical Proteomic Tumor Analysis Consortium (CPTAC), which applied mass spectrometry (MS) for a deeper Px characterization.^38,39^ In 2017, the NCI-CPTAC Dialogue on Reverse Engineering Assessment and Methods (DREAM) Proteogenomics Challenge was launched. Subchallenge 2 was aimed at predicting the Px landscape of 77 breast and 105 ovarian cancer samples from TCGA/CPTAC, based on their respective genomic mutations and Tx landscape.^40,41^ The winner model by Li *et al.* employed a random forest architecture and improved cross-sample agreement from the baseline model’s *r* = 0.44 to 0.53.^11^

In 2020, Magnusson *et al.* isolated naïve CD4^+^ T-helper cells from human donors, and induced them to differentiate into Th1 cells, while measuring RNA sequencing-based Tx and MS-based Px at 7 timepoints along the differentiation protocol (0, 1, 2, 6, 24 hours and 5 days). For each individual gene, they applied a linear model to predict the Px time series based on the Tx time series, with splicing variant resolution. They showed that correlation across samples improved from *r* = 0.21 to 0.86.^4^

In 2022, Srivastava *et al.* published a follow-up paper to the DREAM challenge, where they tested 5 increasingly larger subsets of the dataset, in combination with 3 ML models. They showed that the best-performing models improved with increasing input sizes, with the greatest benefit coming from the introduction of STRING^42^ partners, boosting cross-sample *ρ* from ∼0.3 to 0.599. They applied explainability techniques within the known binding partners of each gene. However, no validation was provided for their results, except for a few anecdotal example genes, showing they were consistent with previous literature.^2^

In the meantime, a handful of clinical multiomics studies had been released, focusing on Alzheimer’s disease (AD) and other dementias. The most notable examples, both published in 2018 and expanded over time, were the Religious Orders Study/Memory and Aging Project (ROSMAP)^43^ and the Mount Sinai Brain Bank (MSBB).^44^ In 2022, Tasaki *et al.*, trained a variational autoencoder-based model^45^ to infer the Px of 384 dorsolateral prefrontal cortex samples, based on their model Tx. The model improved cross-gene correlation from *r* = 0.09 to 0.18. Next, they applied the predictor to the MSBB dataset, where it improved from *r* = 0.09 to 0.14. The authors showed that the proteomized output, compared to raw Tx, performed better at both differential expression analysis and pseudotime analysis, as it matched more closely the Px result and the clinical cognitive decline, respectively. They also applied explainability, although again, without providing biological validation.^3^

Finally in 2024, Cranney and Meyer employed deep learning on a proteomization task, benefitting from the 1,256 samples available in CPTAC by then. However, their paper shares little detail on their model’s performance. While it mentions anecdotal examples of genes being improved, and a correlation distribution before-versus-after proteomization, correlation values are not provided. Once again explainability was applied, but none of their stated results underwent further biological or bioinformatic validation.^46^

Building upon this background, we hereby introduce PROTEOMIZER, a ML platform that applies state-of-the art deep learning to address two fundamental questions. (i) Is proteomization effective at ameliorating differential expression analysis? (ii) Conversely, treating the Tx-Px as a source of precious information rather than a problem, is it possible to leverage it to extract novel biological insight on protein regulation?

To train Proteomizer, we assembled two datasets from TCGA and CPTAC, encompassing three omics, namely transcriptomics (Tx), miRNomics (Mx) and proteomics (Px); the two datasets differed by the Px technique choice: RPPA in one case (*n* = 7,019, 30+ tissues), tandem mass tag MS in the other (*n* = 1,594, 9 tissues). On both datasets, we applied deep learning to predict the Px landscape, based on the Tx and Mx one, in a multi-variable regression task. Proteomizer improved the Tx-Px correlation from *r* = 0.30 to 0.70 in the RPPA version, and from *r* = 0.27 to 0.68 in the MS version, in the test dataset prediction. To this day, this represents the best published performance in the proteomization literature. Next, we applied a Monte Carlo method to simulate thousands of real-world differential expression analysis experiments. The identification of differentially expressed genes (DEGs) became more accurate after proteomization, boosting the average p-value accuracy by up to 2.3× for small sample sizes, and up to 62× for large sample sizes; for a select group of genes (5%), the p-value accuracy boost reached as high as 6 orders of magnitude in the test dataset. Conversely, no benefit could be observed when testing was conducted on an organ that had been excluded from the training set, or when translating the predictions to an independent dataset: this suggests that proteomization models need to “see” the tissue of interest in training, and to receive data that were collected using a consistent protocol. Finally, we applied explainable artificial intelligence (XAI) to identify which genes or miRNAs are most effective at explaining the Tx-Px mismatch, for a given gene of interest (GOI). We applied our analysis to 100 highly annotated GOIs, and cross-compared predictions against a purposely created “Biology Mega Graph”, encompassing 322 million gene-gene or miRNA-gene relations in 69 modalities. Interestingly, in 82 of the 100 GOIs, miRNA annotations appeared as the best correlated with the explainers’ predictions, reaching a ROC AUC of 0.75. These results suggest that Proteomizer explainer is an effective, literature-unbiased tool to predict novel miRNA-gene relations.

To the best of our knowledge, this constitutes the first work that elaborates the biological relevance and limits of a proteomization architecture; and the first time proteomization explainability is cross-referenced to actual biological data, exploring the scope and extent of its predictive power.

## Results

### Mismatch quantification

TCGA data were downloaded from the National Institute of Health (NIH) Genomic Data Commons (GDC) portal,^47^ and CPTAC data were downloaded from the NIH Proteomic Data Commons (PDC) portal.^48^ We isolated the subset of samples where the Tx+Mx+Px triplet was available, leading to the assembly of two independent datasets: (i) one where the Px channel was made by RPPA (*n* = 7,019), and (ii) one where Px was made by MS (*n* = 1,594). Uniform manifold approximation and projection (UMAP) confirmed that tissue-of-origin clusters were clearly identifiable in each of the three omics, in both the RPPA-based dataset (Supplementary Figure 1) and the MS-based one (Supplementary Figure 2).

Before starting, we quantified the Tx-Px mismatch across the samples in both datasets, either as a whole or within individual tissues, or conditions (healthy or cancerous). Of note, mismatch across genes could not be quantified, because neither RPPA nor TMT measure absolute protein quantities. Results are displayed in Table 2, and are largely consistent with those reported in the previous literature.

**Table 2.**
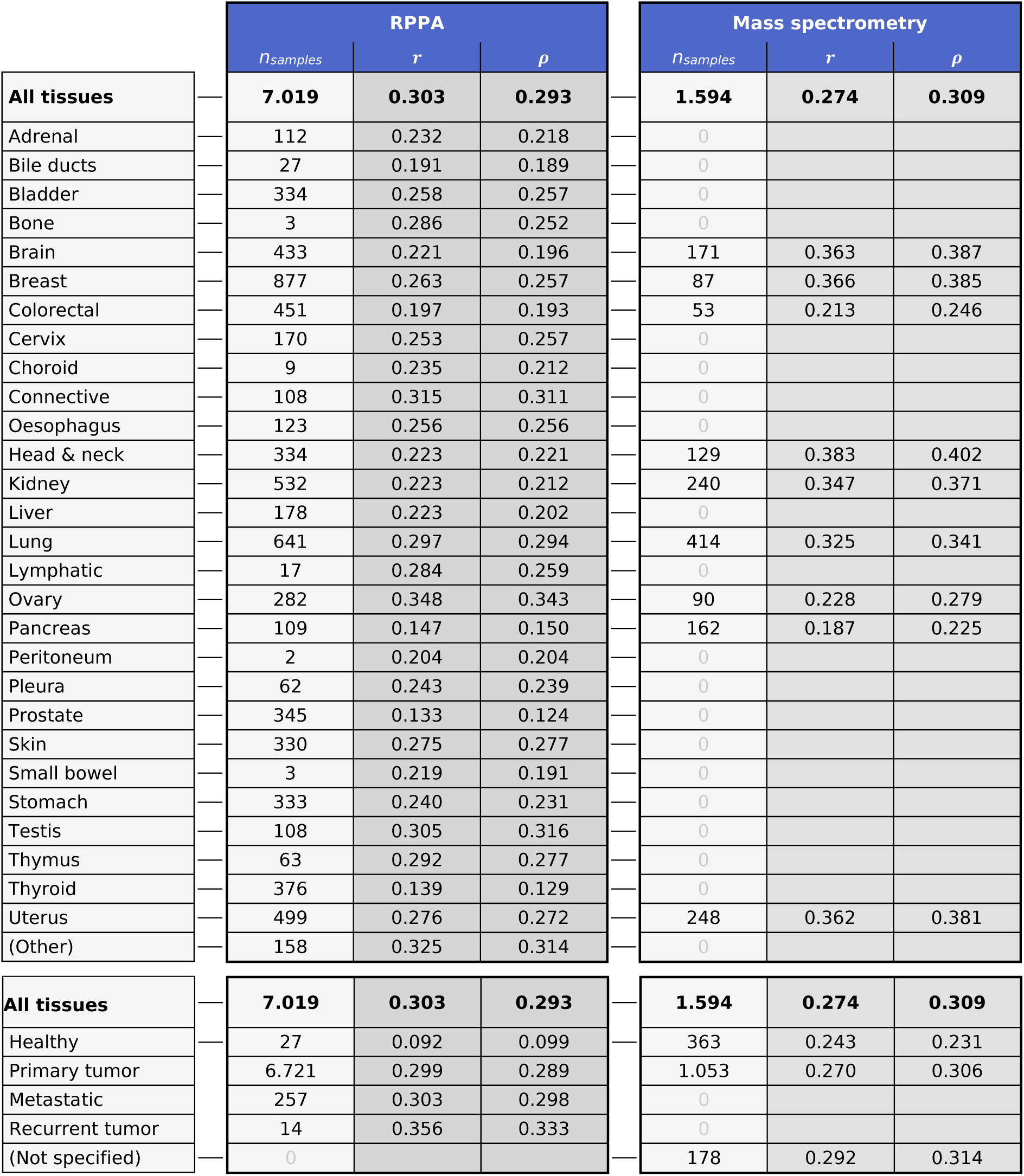
Mismatch in the TCGA and TCGA-CPTAC datasets across tissues of origin and disease states, before proteomization. To train Proteomizer, two independent multiomic datasets were assembled, encompassing a set of healthy and cancerous human biopsies. Both datasets entailed transcriptomic (Tx), miRNomic (Mx) and proteomic (Px) measurements. Version 1 (left table) utilized data from The Cancer Genome Atlas (TCGA), where Tx was assessed by RNA sequencing, Mx by microRNA sequencing, and Px by reverse-phase protein array (RPPA). Version 2 (right table) utilized data from TCGA and Clinical Proteomic Tumor Analysis Consortium (CPTAC), where Tx was assessed by RNA sequencing, Mx by microRNA sequencing, and Px by mass spectrometry, specifically by tandem mass tag (TMT). This table expresses the correlation between Tx and Px reads, measured across samples, in the two versions of the dataset. Values in *n*_samples_ represent the number of samples where the full Tx+Mx+Px triad was available, for each version of the dataset. *r* is the Pearson correlation coefficient, averaged across all genes. Conversely, *ρ* is the signed Spearman correlation coefficient, averaged across all genes. The top part of the table is a breakdown based on the tissues of origin (for metastatic cancer, this corresponds to the primary histological origin of the tumor, e.g., colorectal metastasis in the liver would qualify as colorectal). The bottom part of the table is a breakdown based on the tissue’s nature (healthy or cancerous). For direct comparison with the mismatch after proteomization, please consult Table 3.

### PROTEOMIZER is effective at enhancing Tx-Px correlation metrics

Next, for both versions of the dataset (RPPA and MS), we trained a multi-layer perceptron to predict the Px channel from the Tx and Mx ones. Both experiments were carried out in a 10-fold cross-validation setting, stratified with respect to the tissues of origin. We optimized the hyperparameters by a grid search, using early stop to prevent overfitting. Performance was measured in terms of mean absolute error (MAE), benefitting from the fact that both RPPA and MS data had been normalized by gene-centric Z-scoring, therefore, they were expressed in the form of “number of standard deviations”. The best MAEs recorded were 0.498 for RPPA, and 0.367 for MS, both of which appeared largely insensitive to the hyperparameter choice (Supplementary Figure 5)

Proteomization proved remarkably effective at improving the correlation metrics in the prediction from the test dataset. Cross-sample Pearson *r* improved from 0.30 to 0.71 for RPPA, and from 0.21 to as high as 0.68 for MS. Likewise, cross-sample Spearman *ρ* improved from 0.29 to 0.69 for RPPA, and from 0.31 to 0.54 for MS (Figure 1, Supplementary Figure 6). We also checked whether the performance varied based on the tissue of origin, or the disease state. MAE, Pearson *r* and Spearman *ρ* all appeared remarkably steady, suggesting that the prediction quality is comparable, with respect to tissues or disease state (Table 3).

**Figure 1.**
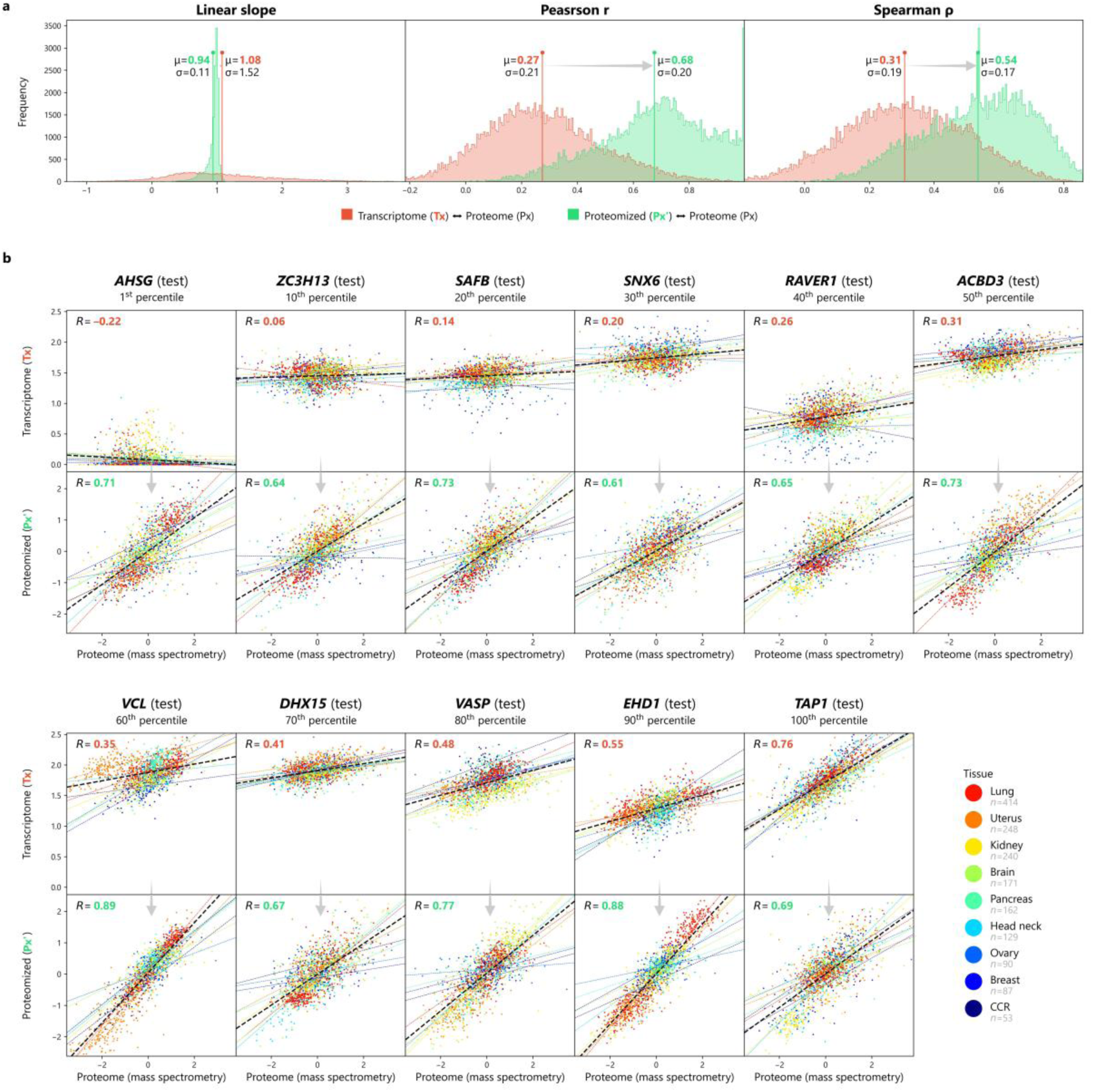
Correlation metrics before and after proteomization, in the mass spectrometry version of the TCGA-CPTAC dataset. Proteomizer was applied to predict the mass spectrometry proteomic (Px) landscape, based on the transcriptomic (Tx) and miRNomic (Mx) ones, on 1,594 samples sourced from The Cancer Genome Atlas (TCGA) and Clinical Proteomic Tumor Analysis Consortium (CPTAC). After training, the predicted Px’ in the test dataset was compared with the original Tx and Px. (a-c) Three correlation metrics—namely the linear slope, the Pearson *r* correlation coefficient and the Spearman *ρ* correlation coefficient—were applied between raw Tx and Px (red), and between predicted Px’ and Px (green), for each of the 14,041 genes with both Tx and Px availability; the distribution profiles show the effect of proteomization, in a before-after fashion. (d) Scatter plots of the 1,594 samples for 11 exemplary genes, depicting the ground truth Px coordinates on the x-axis, and the raw Tx (top) or the predicted Px’ (bottom) on the y-axis. Genes were sampled in a diverse range of Tx-Px mismatch values, from the most mismatched (1^st^ percentile) to the most aligned (100^th^ percentile). Coordinates are identical from one gene to another. Colors indicate the tissue of origin (see Table 2). A linear regression line was computed in each plot, both for the entire gene set (black dashed line), and for individual organs (solid colored lines).

**Table 3.** Mismatch in the TCGA and TCGA-CPTAC datasets across tissues of origin and disease states, after proteomization. In order to train Proteomizer, two independent multiomic datasets were assembled, encompassing a set of healthy and cancerous human biopsies. Both datasets entailed transcriptomic (Tx), miRNomic (Mx) and proteomic (Px) measurements. Version 1 (left table) utilized data from The Cancer Genome Atlas (TCGA), where Tx was assessed by RNA sequencing, Mx by microRNA sequencing, and Px by reverse-phase protein array (RPPA). Version 2 (right table) utilized data from TCGA and Clinical Proteomic Tumor Analysis Consortium (CPTAC), where Tx was assessed by RNA sequencing, Mx by microRNA sequencing, and Px by mass spectrometry, specifically tandem mass tag (TMT). Tx underwent proteomization by the Proteomizer deep learning model, generating a predicted proteome (Px’) from the test dataset prediction. This table expresses the correlation between Px’ and Px reads, measured across samples, in the two versions of the dataset. Values in *n*_samples_represent the number of samples where the full Tx+Mx+Px triad was available, for each version of the dataset. MAE is the mean absolute error, which was utilized in Proteomizer for both the early stop, and hyperparameter selection (both of which were conducted in the validation dataset). *r* is the Pearson correlation coefficient, averaged across all genes. Conversely, *ρ* is the signed Spearman correlation coefficient, averaged across all genes. The top part of the table is a breakdown based on the tissues of origin (for metastatic cancer, this corresponds to the primary histological origin of the tumor, e.g., colorectal metastasis in the liver would qualify as colorectal). The bottom part of the table is a breakdown based on the tissue’s nature (healthy or cancerous). For direct comparison with the mismatch before proteomization, please consult Table 2.

|  | RPPA |  |  |  | Mass spectrometry |  |  |  |
| --- | --- | --- | --- | --- | --- | --- | --- | --- |
| | $n_{\text{samples}}$ | MAE | $r$ | $\rho$ | $n_{\text{samples}}$ | MAE | $r$ | $\rho$ |
| **All tissues** | **7.019** | **0.497** | **0.706** | **0.690** | **1.594** | **0.363** | **0.675** | **0.536** |
| Adrenal | 112 | 0.503 | 0.533 | 0.521 | 0 |  |  |  |
| Bile ducts | 27 | 0.697 | 0.314 | 0.308 | 0 |  |  |  |
| Bladder | 334 | 0.571 | 0.452 | 0.454 | 0 |  |  |  |
| Bone | 3 | 0.586 | 0.633 | 0.575 | 0 |  |  |  |
| Brain | 433 | 0.436 | 0.506 | 0.493 | 171 | 0.359 | 0.402 | 0.385 |
| Breast | 877 | 0.519 | 0.468 | 0.458 | 87 | 0.365 | 0.225 | 0.232 |
| Cervix | 170 | 0.544 | 0.449 | 0.453 | 0 |  |  |  |
| Choroid | 9 | 0.501 | 0.181 | 0.163 | 0 |  |  |  |
| Colorectal | 451 | 0.497 | 0.449 | 0.443 | 53 | 0.483 | 0.154 | 0.182 |
| Connective | 108 | 0.549 | 0.531 | 0.508 | 0 |  |  |  |
| Oesophagus | 123 | 0.584 | 0.358 | 0.361 | 0 |  |  |  |
| Head & neck | 334 | 0.555 | 0.407 | 0.400 | 129 | 0.348 | 0.363 | 0.377 |
| Kidney | 532 | 0.447 | 0.642 | 0.637 | 240 | 0.335 | 0.539 | 0.516 |
| Liver | 178 | 0.452 | 0.496 | 0.485 | 0 |  |  |  |
| Lung | 641 | 0.480 | 0.500 | 0.499 | 414 | 0.324 | 0.433 | 0.442 |
| Lymphatic | 17 | 0.510 | 0.292 | 0.268 | 0 |  |  |  |
| Ovary | 282 | 0.439 | 0.465 | 0.460 | 90 | 0.559 | 0.150 | 0.161 |
| Pancreas | 109 | 0.560 | 0.298 | 0.294 | 162 | 0.381 | 0.333 | 0.281 |
| Peritoneum | 2 | 0.822 | 0.447 | 0.447 | 0 |  |  |  |
| Pleura | 62 | 0.594 | 0.321 | 0.339 | 0 |  |  |  |
| Prostate | 345 | 0.398 | 0.359 | 0.353 | 0 |  |  |  |
| Skin | 330 | 0.548 | 0.436 | 0.440 | 0 |  |  |  |
| Small bowel | 3 | 0.566 | 0.204 | 0.173 | 0 |  |  |  |
| Stomach | 333 | 0.537 | 0.457 | 0.453 | 0 |  |  |  |
| Testis | 108 | 0.428 | 0.561 | 0.561 | 0 |  |  |  |
| Thymus | 63 | 0.426 | 0.528 | 0.494 | 0 |  |  |  |
| Thyroid | 376 | 0.471 | 0.521 | 0.512 | 0 |  |  |  |
| Uterus | 499 | 0.504 | 0.543 | 0.521 | 248 | 248 | 0.357 | 0.353 |
| (Other) | 158 | 0.512 | 0.666 | 0.656 | 0 |  |  |  |
| **All tissues** | **7.019** | **0.497** | **0.706** | **0.690** | **1.594** | **0.363** | **0.675** | **0.536** |
| Healthy | 27 | 0.555 | 0.254 | 0.224 | 363 | 0.291 | 0.650 | 0.515 |
| Primary tumor | 6.721 | 0.495 | 0.704 | 0.690 | 1.053 | 0.389 | 0.660 | 0.511 |
| Metastatic | 257 | 0.536 | 0.491 | 0.469 | 0 |  |  |  |
| Recurrent tumor | 14 | 0.546 | 0.635 | 0.607 | 0 |  |  |  |
| (Not specified) | 0 | 0.497 | 0.706 | 0.690 | 178 | 0.356 | 0.597 | 0.491 |

### Gene set enrichment analysis

Next, we investigated what biological functions were shared among the genes that showed (i) the best or worst Tx-Px alignment, (ii) the best/worst Px’-Px alignment, and (iii) the strongest or faintest improvement before versus after proteomization. To this end, we performed gene set enrichment set analysis (GSEA) on the Spearman *ρ* coefficients from the MS prediction, against the Gene Ontology (GO) term set.^49^ Of note, the RPPA version of the dataset could not be utilized for this purpose, as the RPPA panel only encompasses 454 antibodies, targeting a total of 398 unique genes.

(i) In the original dataset (Figure 2a), the best Tx-Px alignment (minimum mismatch) was found in a rather heterogeneous group of gene classes, mostly related to the cell structural apparatus, be it collagen, integrins, extracellular matrix or adhesion. Conversely, the worst Tx-Px alignment (maximum mismatch) was dominated by energy-related functions, most notably aerobic respiration. (ii) In Proteomizer’s prediction (Figure 2b), the best Px’-Px alignment (minimum mismatch) was strongly enriched by cytoskeletal functions, most notably related to actin and myosin. Of note, the GO “cellular respiration” class appeared among the top hits, despite it also listed among the most mismatched classes in the original dataset. Conversely, the worst performing classes (maximum mismatch) highlighted post-translational modifications, especially ubiquitination and transfer-RNAs (tRNAs). It is remarkable that “hard-to-predict” gene classes have so much to do with the cell machinery that is directly responsible for the Tx-Px mismatch in the first place. (iii) Finally, the classes that improved the most from proteomization (Figure 2c) were dominated by ribosomes and aerobic respiration. Conversely, the least-improved pool focused on transient processes, essentially cytokine-related terms (growth factor response, cytokine response, GTPase pathways, MAP kinase, etc.) and inflammation-related ones (inflammation response, symbiont entry into host, virus receptor activity, inflammation response, interferon pathway interleukin-2, etc.).

**Figure 2.**
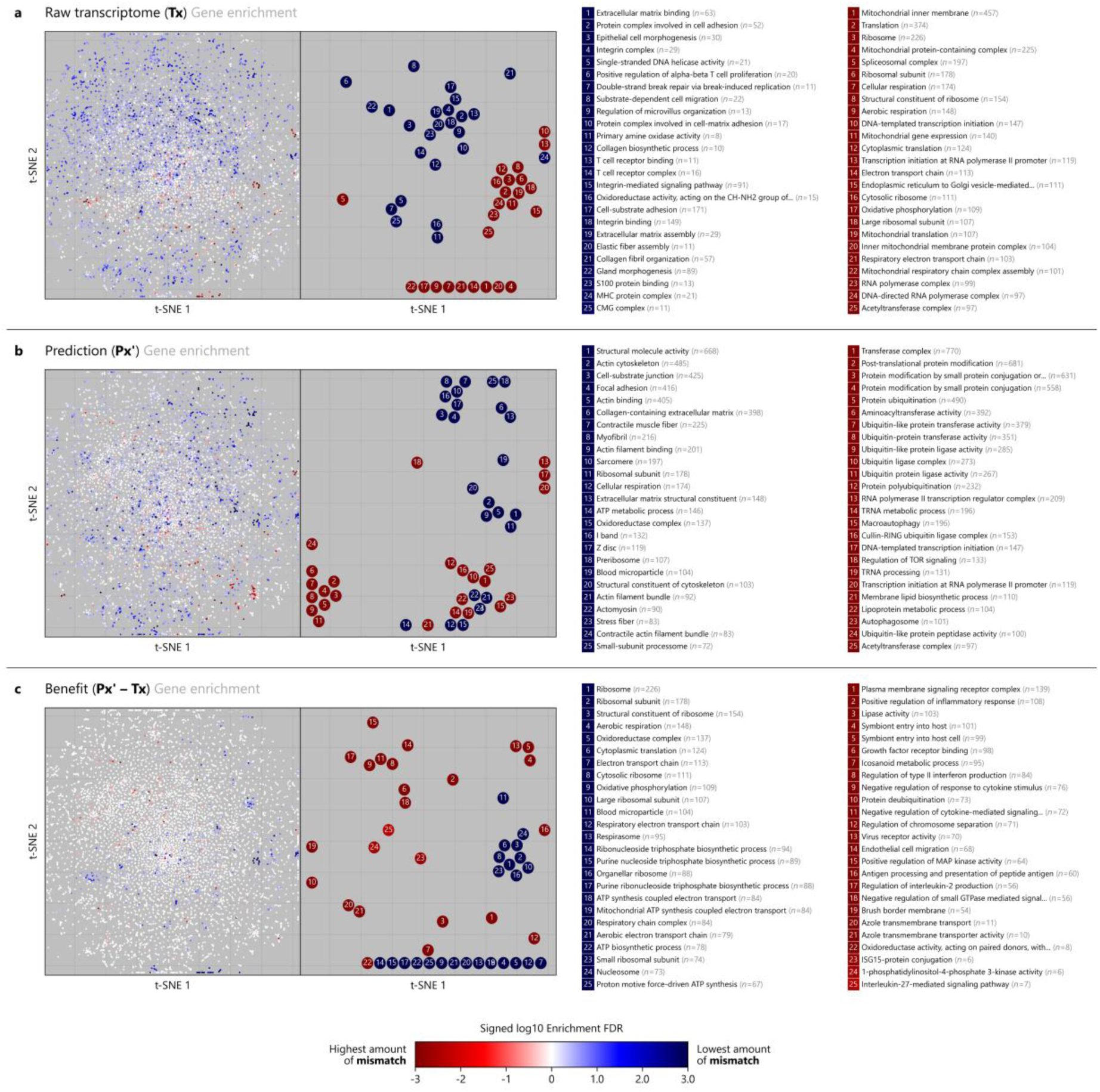
Gene set enrichment analysis of the transcriptomic-proteomic mismatch before proteomization, after proteomization, and in terms of differential benefit from proteomization, in the mass spectrometry version of the TCGA-CPTAC dataset. Proteomizer was applied to predict the mass spectrometry proteomic (Px) landscape, based on the transcriptomic (Tx) and miRNomic (Mx) ones, on 1,594 samples sourced from The Cancer Genome Atlas (TCGA) and Clinical Proteomic Tumor Analysis Consortium (CPTAC). For all genes, mismatch was quantified between raw Tx and ground-truth Px in terms of Spearman correlation coefficient *ρ*_1_, resulting in gene ranking (a). Next, mismatch was quantified between the predicted Px’ and the ground-truth Px *ρ*_2_, resulting in gene ranking (b). Finally, the proteomization benefit was determined as the difference between the two coefficients, *ρ*_2_ − *ρ*_1_, resulting in gene ranking (c). Next, gene set enrichment analysis was applied to the three gene rankings, against all Gene Ontology classes of sizes between 5 and 2,000 members. These resulted in false discovery rate (FDR) values being computed for each GO class, from each of the three gene rankings. (Column 1) FDR scores were plotted against a t-distributed stochastic neighbor embedding (t-SNE) plot, where dots represent individual gene ontology classes. The t-SNE was realized on the basis of their gene membership Boolean table, as a result, it showcases the localization of each GO class in the space of cell functions. For example, GO terms related to aerobic respiration colocalize at the bottom-right of the plot. (Column 2) The top 25 GO classes (strongest alignment, i.e., minimum mismatch) and the bottom 25 GO classes (weakest alignment, i.e., maximum mismatch) were individually annotated to highlight their localization, and the presence of biologically correlated clusters. (Columns 3-4) Finally, the top- and bottom-performing classes were individually listed, together with their respective gene numerosity. Coordinates are identical across all six plots. t-SNE parameters: perplexity=“auto”.

Overall, these results confirm that, not only is the mismatch problem remarkably variable across different gene classes, but some biological fields like cell energy and ribosome biology may particularly benefit from ML proteomization.

### Monte Carlo simulations for differential gene expression analysis

While correlation has been the go-to metric to quantify the Tx-Px mismatch throughout bioinformatic literature (Table 1), we argue that it does not necessarily convey biological relevance. In real-world applications, virtually all use cases of gene expression analysis involve differential analysis between two sample sets, differing by a certain “perturbation” —be it disease, treatment or stimulus. When biologists perform a differential Tx experiment between two samples, they are seeking a list of proteins that are significantly up- or downregulated as a consequence of the perturbation. Tx is effective as long as differential Tx is a good proxy for differential Px— or, more explicitly, the differentially expressed genes (DEGs) at Tx level match the “ground truth” DEGs at Px level.

In light of the major correlation improvements produced by proteomization, we sought to assess whether proteomization also brought forth greater accuracy in terms of differential expression analysis, i.e., DEG identification. For this purpose, we designed a novel Monte Carlo method that simulates a real-world scenario of differential expression analysis, on the basis of the TCGA-CPTAC datasets. The algorithm works as follows.

Before starting, samples within each tissue of origin undergo clustering, based on their Px landscape (assumed as the ground truth). Clustering is performed by *k*-mean clustering, where *k* is optimized by grid-searching the optimal silhouette score between 2 and 10. We selected unsupervised clustering, as it is the most established technique to separate omic samples by their overall biology, for example, it is widely used in single-cell Tx for the identification of different cell types,^50^ and in cancer to perform precision diagnostics.^51^ For instance, in the MS dataset: kidney samples divide almost perfectly the clear-cell and non-clear cell renal carcinomas. The effectiveness of this method was validated by principal component analysis (PCA), which showed that cluster labels generally colocalized together forming tidy, well-defined groups. Pairwise comparisons between samples of distinct biologies reasonably exemplify real-world use cases of differential Tx (as long as the samples are from the same organ of origin). It is very plausible that a biologist may be interested in finding out the DEGs separating healthy from cancerous, or small-cell from non-small-cell lung cancers; whereas, it would be much less plausible to apply differential analysis between two entirely different organs, say, liver versus uterus.

After establishing clusters, the algorithm proceeds in a randomized, iterative fashion.

1. For a given organ, say, pancreas, it randomly picks 2 clusters.
2. Next, it randomly selects 3 samples from cluster A, and 3 samples from cluster B. a
3. Next, for each gene, it applies Wald test (arguably the most robust and most common statistical test in differential expression analysis^52^) to compute a p-value between the Px readouts from group A, versus the corresponding Px readout from group B; the p-value is then thresholded into a label, which represents the gene’s DEG status: label “+1” if the gene is significantly (*p* < 0.05) upregulated in the A samples over the B samples; label “–1” if the gene is significantly (*p* < 0.05) downregulated; label “0” if the gene is not significantly differentially expressed between the two sample groups.

The p-value (and its cognate label) between the Px reads in group A versus the Px reads in group B corresponds to the ground truth (gt) differential analysis result. The same procedure is repeated by comparing the Tx reads in group A versus the Tx reads in group B, which corresponds to the current state of affairs (raw), where Tx is used as a proxy for Px-based differential expression analysis. Finally, it is repeated once again, comparing Proteomizer’s prediction Px’ in group A versus the Px’ in group B, which represents our proposed ML-based proteomization (pred). The general hypothesis is that gt’s p-value should be closer to pred’s p-value, whereas raw’s p-value should be farther away; equivalently GT’s label will agree with pred’s label more often than raw’s label agrees with GT’s label.

Finally, steps (1-3) are combinatorially repeated: for different statistical tests, namely Student’s t-test, Mann-Whitney/Wilcoxon rank-sum u-test and Wald w-test; for different p-value thresholds, namely *p* < 0.5, < 0.1, and *p* < 0.01; for all genes in the dataset; for 100 randomly selected combinations of A and B samples; for multiple sample sizes, going from 3, upwards; for all tissues of origin.

Because omics are subject to noise, a higher sample size brings along greater statistical power, resulting in more comparisons reaching statistical significance. At low sample sizes, it is trivial to predict DEGs, as almost all genes are non-significant due to limited power: therefore, a “dummy” classifier that always outputs a “0” label is almost always correct. This changes dramatically at higher sample sizes, especially considering that our p-values are not corrected for false discovery rate, making significant comparisons very likely at sample sizes ≥10.

At the end of the experiment, the benefit delivered by proteomization is calculated by assessing the mean Δ log p-value between Px’ and Px, against the Δ log p-value between Tx and Px; if the former is smaller than the latter, it implies that proteomization was beneficial. Equivalently, one could assess the average agreement between the labels (at, say, *p* < 0.05) of Px’ with Px, against the agreement between the labels of Tx with Px; if the former is larger than the latter, it implies that Proteomization was beneficial. The labels have the advantage of being more human-interpretable than a Δ log p-value, as they express a percentage of correctly classified genes (DEG vs non-DEG).

We applied our Monte Carlo method to assess Proteomizer’s predictions (Figure 3, Supplementary Figure 7). In both RPPA and MS, results appeared moderately successful for small sample sizes (*n* = 3-8), showcasing a mean p-value boost of 1.2-2.8×; and extremely successful at large sample sizes (*n* = 10-50), with a p-value boost reaching as high as 62.3×.

**Figure 3.**
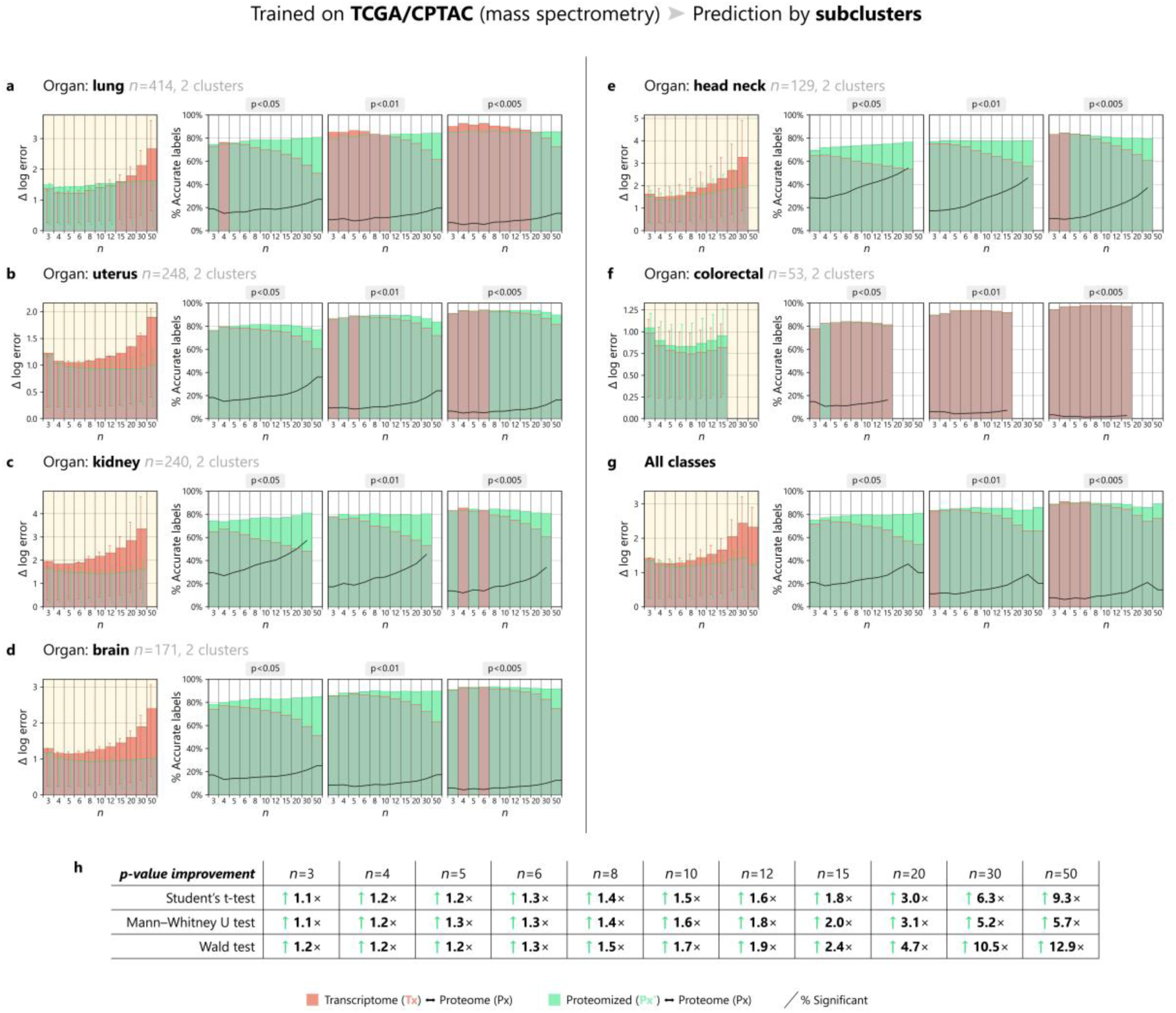
Monte Carlo simulation of differential expression analyses across sample clusters, in the mass spectrometry version of the TCGA-CPTAC dataset. Proteomizer was applied to predict the mass spectrometry proteomic (Px) landscape, based on the transcriptomic (Tx) and miRNomic (Mx) ones, on 1,594 samples sourced from The Cancer Genome Atlas (TCGA) and Clinical Proteomic Tumor Analysis Consortium (CPTAC). At the end of the training, three full landscapes were made available: (i) the raw Tx, (ii) the predicted Px’ and (iii) the ground truth Px. Next, a Monte Carlo method was applied across sample clusters within each tissue of origin. The rationale was to determine whether proteomization had been beneficial, with respect to the customary endpoint of gene expression tests, that is differential expression analysis; in less formal words, the task was to assess whether Px’ would do a better job at predicting the differentially expressed genes (DEGs) in Px, compared to Tx. While a complete description of the Monte Carlo method is available in the Materials and Methods, here follows a short summary. The analysis is repeated within the scope of each individual tissue of origin (a-f). First, samples are divided into 2-10 clusters by unsupervised machine learning (ML), specifically, by *k*-means clustering where *k* is optimized using the silhouette score. Next, the following iteration step is repeated 100 times, for varying sample size values *n* (x-axes in the figure). A number *n* of samples are randomly selected from one of the clusters, and the same number *n* of samples are randomly selected from a different cluster: they work as the experimental (exp) and control group (ctrl) respectively, of a hypothetical differential expression analysis experiment performed by a biologist. All genes undergo differential expression analysis by Wald test, comparing Txexp with Txctrl, Px’exp with Px’ctrl, and Pxexp with Pxctrl; these three comparisons yield a set of p-values for each gene. (Column 1) The p-values are log-transformed and assigned a sign, namely: “+” for the genes upregulated in the experimental group, “–” for the downregulated ones. The average difference between the log-transformed signed p-values of the Px’ with respect to Px constitutes the “Δ log error’ of Px’, and is depicted in green. Likewise, the average difference between the log-transformed signed p-values of the Tx with respect to Px constitutes the “Δ log error” of Tx, and is depicted in red. If the proteomization is beneficial, then the green “Δ log error” in Px’ is supposed to be lower than in Tx—thus, leaving an emerging silhouette of red color standing out over the green. (Column 2) The p-values in Tx, Px’ and Px are flattened into labels, according to the following rule: label “+1” if a gene is significantly (*p* < 0.05) upregulated in the experimental group; label “–1” if a gene is significantly (*p* < 0.05) downregulated in the experimental group; label “0” if the comparison is not significant. Then, the number of genes where Px’ agrees with Px is counted, and goes to constitute the “% accurate labels” for Px’, which is depicted in green. Likewise, the number of genes where Tx agrees with Px is counted, and goes to constitute the “% accurate labels” for Tx, which is depicted in red. If the proteomization is beneficial, then the green “% accurate labels” in Px’ is supposed to be lower than in Tx—thus, leaving an emerging silhouette of green color standing out over the red. The black line depicts the % of significant comparisons; of note, p-values were not corrected for multiple testing (i.e., no Bonferroni or Benjamini-Hochberg corrections were applied), thereby it is expected that the percentage of significant comparisons will be strongly inflated. (Columns 3-4) Same as column 2, except the p-value threshold is fixed at 0.01 and 0.005, respectively (instead of 0.05); this makes the scenario of a statistically significant comparison rarer. (g) Aggregate measures across all tissues of origin. (h) Relative p-value improvement (average p-value for Px’ / average p-value for Tx) at different sample sizes, and using different statistics, namely: Student’s t-test, Mann-Whitney/Wilcoxon rank-sum u-test and Wald w-test. Overall, proteomization had a beneficial effect in the mass spectrometry version of the TCGA-CPTAC dataset, yielding an average increase in Wald test “Δ log error” ranging from 1.2× (*n* = 3) up to 12.9× (*n* = 50). The benefit appears to be proportional to the sample size, being relatively marginal with small samples due to low statistical power, and becoming very robust at larger samples, where the number of incorrectly identified labels is reduced by a factor of ∼50% (panel g, columns 2-4). All tissues registered an improvement, with the sole exception of the colorectal. These results confirm that, as long as the tissues of interest are included in the training dataset, machine learning is an effective tool to alleviate the Tx-Px mismatch.

In light of the evidence that genes belonging to different biological processes display different degrees of improvement as a result of proteomization, we employed the validation test results to select the top 5% most improved genes, and we repeated the Monte Carlo method on the test dataset, using only these selected genes. Results show that average Wald test p-values improve by up to 3 orders of magnitude at small samples sizes (*n* = 3-8), and by up to 6 orders of magnitude at large sample sizes (*n* = 10-50) (Figure 4). This result indicates that ML-based proteomization can massively increase DEG identification accuracy for a select group of genes.

**Figure 4.**
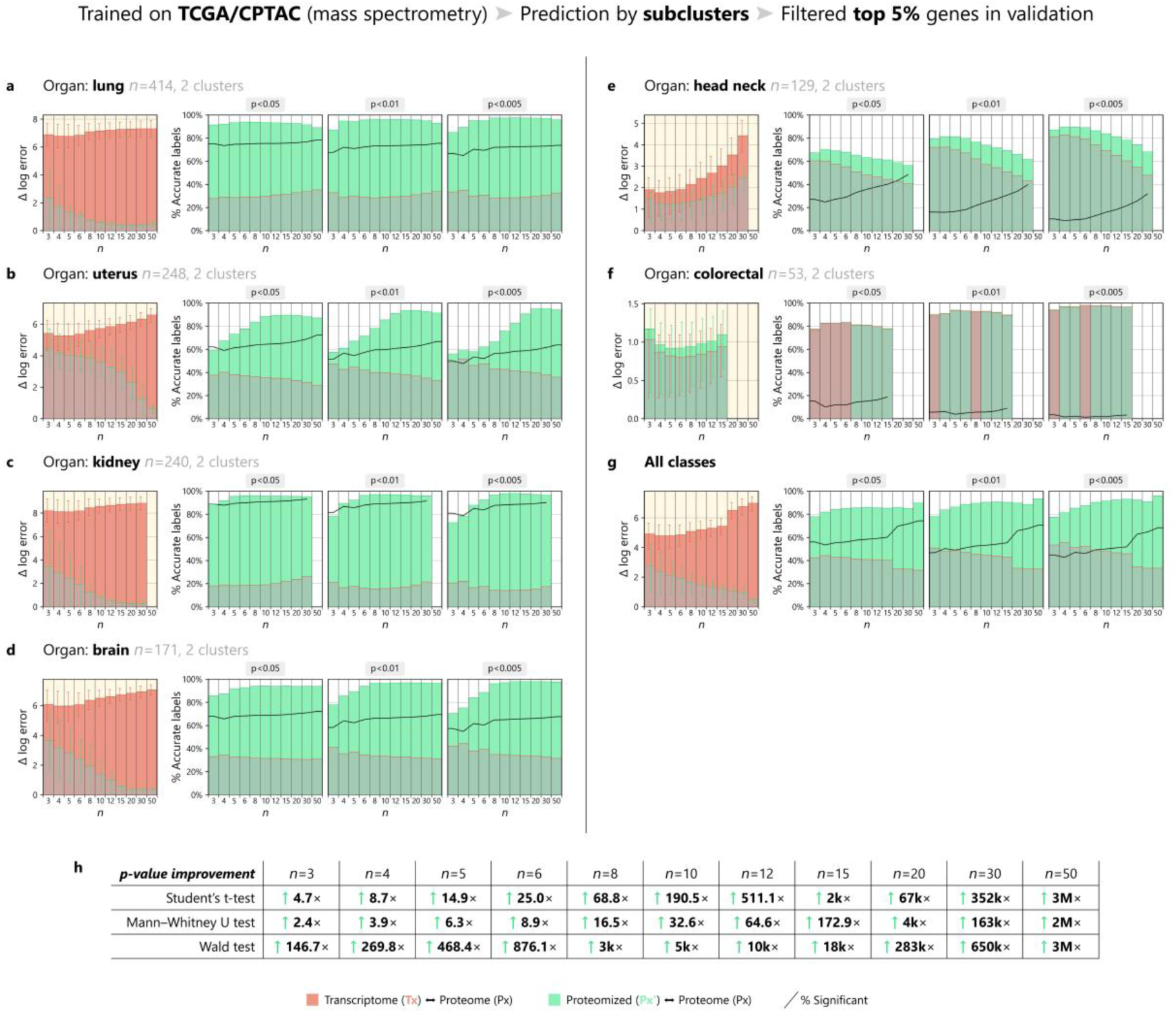
Monte Carlo simulation of differential expression analyses across sample clusters, in the mass spectrometry version of the TCGA-CPTAC dataset, filtering for the top 5% best performer genes in the validation dataset. The figure is obtained identically to Figure 3. The only difference is that genes were filtered, by selectively including the top 5% genes that benefited the most from proteomization in the validation dataset. Of note, the Monte Carlo method was applied on the test dataset prediction: the selection process was kept completely separate from the testing, to prevent data leakage and guarantee reproducibility. The best performer genes were selected by the average improvement in the “Δ log error” of Px’ achieved at Student’s t-test, in the sample sizes 3, 4, 5 and 8.

### Proteomization is not transferable across unseen tissues or data collection protocols

We then proceeded to examine whether Proteomizer was effective at predicting DEGs between sample clusters from a tissue type that was not utilized during training. First, we trained Proteomizer using a leave-one-group-out (LOGO) cross-validation structure: while classic stratified cross-validation ensures that all tissues of origin are equally represented across the cross-validation folds, LOGO includes all tissues but one for training and validation, and finally tests performance on the held-out tissue; the process is recursively repeated, holding out one tissue at a time. In the MS dataset, a total of 9 tissues were available in the dataset: hence, LOGO was repeated for a total of 9 folds; once trained across 8 tissues, the model was tasked to predict the Px landscape of the 9^th^ held-out tissue; after training, the Monte Carlo method was employed, to measure Proteomizer’s ability in identifying the DEGs across 2 or more subclusters from said 9^th^ tissue. Sadly, Proteomizer was not effective at identifying the DEGs, and its overall effect was either neutral or pejorative, irrespective of the tissue or the sample size (Figure 5).

**Figure 5.**
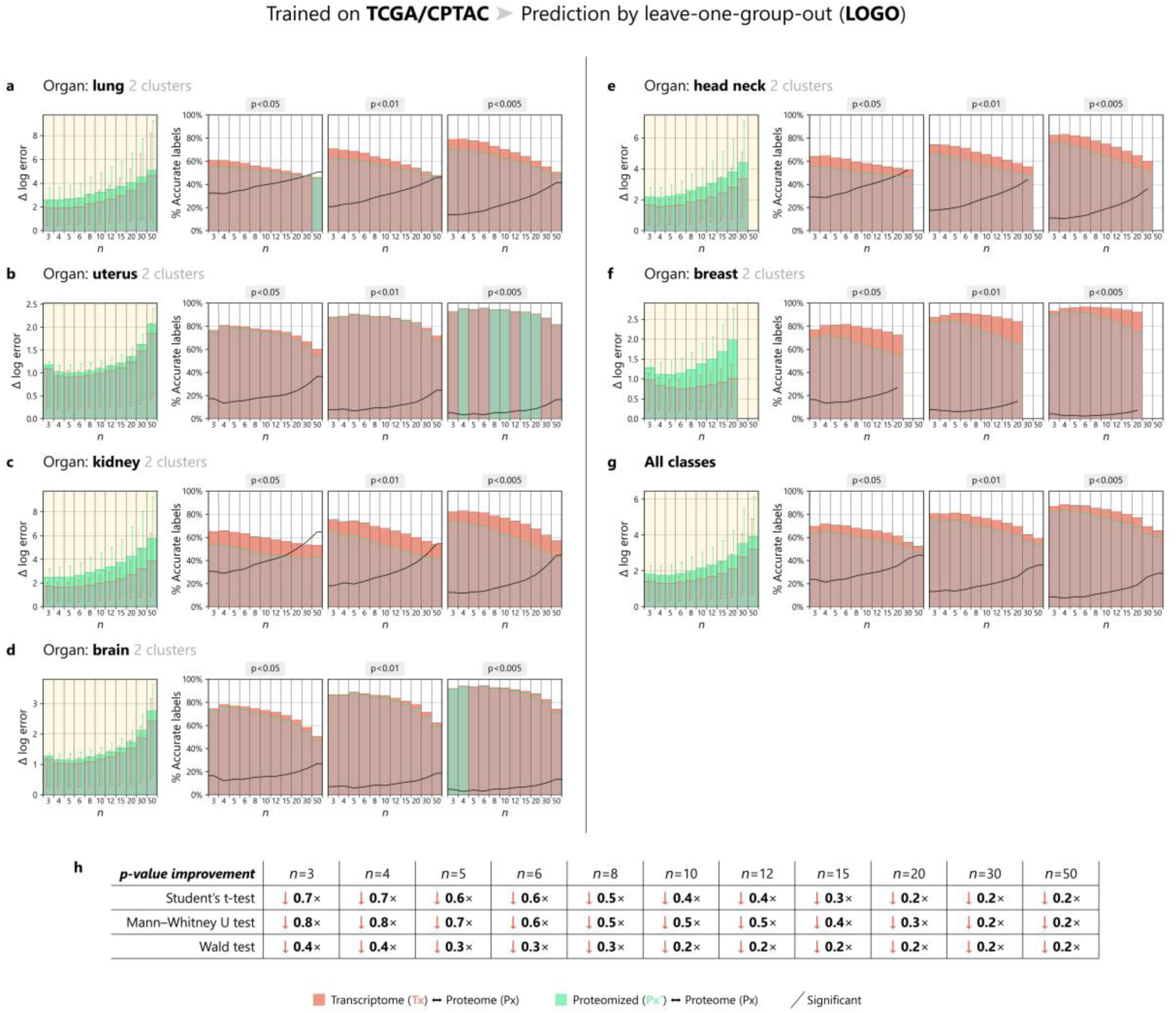
Monte Carlo simulation of differential expression analyses in a leave-one-group-out setting, in the mass spectrometry version of the TCGA-CPTAC dataset. Proteomizer was applied to predict the mass spectrometry proteomic (Px) landscape, based on the transcriptomic (Tx) and miRNomic (Mx) ones, on 1,594 samples sourced from The Cancer Genome Atlas (TCGA) and Clinical Proteomic Tumor Analysis Consortium (CPTAC). At the end of the training, three full landscapes were made available: (i) the raw Tx, (ii) the predicted Px’ and (iii) the ground truth Px. Next, a Monte Carlo method was applied in a leave-one-group-out (LOGO) fashion; LOGO consists of training the machine learning model on all tissues but one, and then testing the predictor on the held-out tissue. The Monte Carlo method was applied as described in Figure 3 (a full description can be found in Materials and Methods). Overall, proteomization had a net neutral or disruptive effect across all tested tissues, sample sizes and p-value thresholds. These results suggest that machine learning-based proteomization is not effective at translating to a completely novel tissue type.

We also assessed whether Proteomizer would be capable of making predictions on completely independent datasets sourced from Alzheimer’s disease Knowledge Portal^53^, namely: the Religious Orders Study/Memory and Aging Project (ROSMAP) study,^54,55^ the Mount Sinai Brain Bank (MSBB) study^44,56^ and the ROSMAP iPSC-Derived Induced Neurons (ROSMAP-IN) study.^57,58^ ROSMAP and MSBB were conducted on post-mortem brain samples in a cohort of aged subjects, some of whom received clinical or pathological diagnosis of Alzheimer’s disease (AD), or other dementia; instead, RISMAP-IN utilized neurons derived from induced pluripotent stem cells (iPSCs), sampled from ROSMAP subjects. After training Proteomizer on all the 9 tissues of origin in the MS version of the TCGA-CPTAC dataset (including the brain), we tasked it to predict the proteomic differences between healthy and AD subjects in 6 ROSMAP/MSBB sub-studies; unfortunately, Proteomizer was unsuccessful, yielding mostly neutral effects. Because neurodegeneration strongly impacts proteostasis, we repeated the experiment comparing male subjects versus female subjects, with similar results (Supplementary Figure 8).

Overall, these results suggest that proteomization is effective as long as the tissue in question has been included in the training cohort, and the multiomic measurements are made consistently between the dataset used for training and the dataset used for testing.

### PROTEOMIZER EXPLAINER is effective at identifying gene regulatory relations

Finally, we wished to investigate whether the Tx-Px mismatch could be exploited as a source of biological information, rather than as a mere problem. To this end, we applied explainable artificial intelligence (XAI), a ML approach that encompasses various families of “explainer” models to make ML models human-understandable. Their use is considered an integral part of today’s best practices in the field.^59^ In our case, the fundamental question was the following:

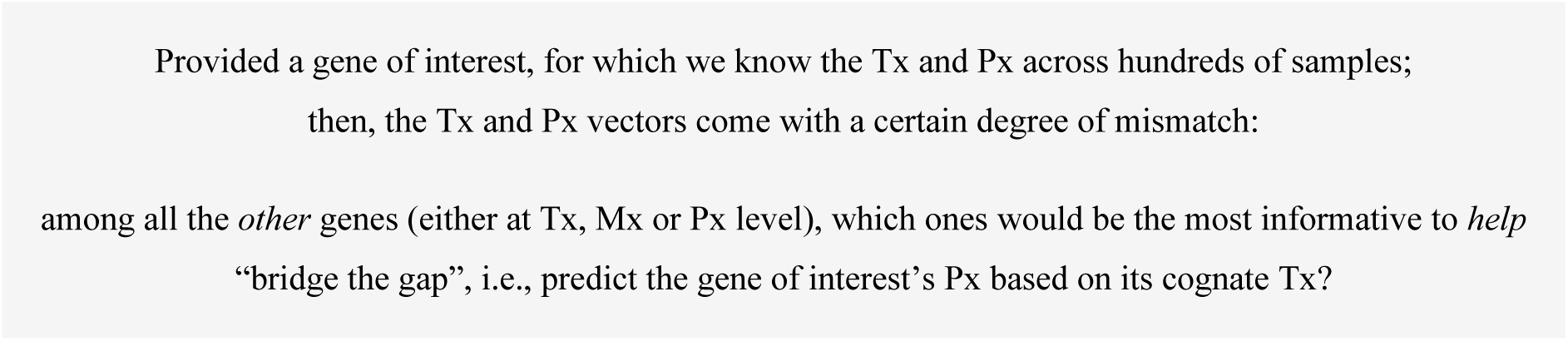

We will hereinafter refer to the gene of interest as the GOI, and the genes other-than-the-gene-of-interest as the helper genes, or the helper columns. The helper genes are essentially all non-GOI’s Tx and Px columns in the dataset, plus all the columns in the Mx dataset.

From a more biological perspective, the rationale is the following. If a helper gene is effective at bridging the information gap between the GOI’s Tx and the GOI’s Px, two explanations are possible, other than chance: either the helper is *<u>coregulated</u>* with the GOI, implying that their similarity is just an epiphenomenal correlation; or, the helper is a *<u>regulator</u>* of the GOI, embodying a causal relation. For example, a good helper gene may be a ubiquitin ligase that is causally responsible for a GOI’s accelerated turnover.

A list of 100 GOIs was employed for this purpose (further elaborated below). The training was repeated both in “direct” mode, i.e., trying to predict the GOI’s Px based on its Tx; and in “reverse” mode, i.e., trying to predict the GOI’s Tx based on its Px. Of note, the information “gap” is conceptually the same in direct and reverse. Overall, the project was organized in 6 tasks:

- task 1: starting from the GOI’s Tx, and helpers’ Tx, predicting the GOI’s Px (direct);
- task 2: starting from the GOI’s Tx, and helpers’ Px, predicting the GOI’s Px (direct);
- task 3: starting from the GOI’s Tx, and helpers’ Mx, predicting the GOI’s Px (direct);
- task 4: starting from the GOI’s Px, and helpers’ Tx, predicting the GOI’s Tx (reverse);
- task 5: starting from the GOI’s Px, and helpers’ Px, predicting the GOI’s Tx (reverse);
- task 6: starting from the GOI’s Px, and helpers’ Mx, predicting the GOI’s Tx (reverse).

We compared three explainer architectures, namely: Extreme Gradient Boosting (XGBoost)^60^ importance scores, Shapley Additive exPlanations (SHAP),^61^ and a novel approach called Information Gain. Each of the three explainers produced a ranking of the helpers, based on how important they were in explaining the GOI’s Tx-Px mismatch. The top-ranked helpers are likely to be regulators of the GOI, or genes that are co-regulated with the GOI.

To validate the biological meaningfulness of our predictions, we wished to cross-compare our top-ranked helpers against a set known gene-gene or miRNA-gene relations. To this end, we created the Biology Mega Graph, a knowledge graph encompassing 322,255,334 known regulatory relations, which is part of our VLab ML library.^62^ The graph was created by harmonizing together 11 biological repositories, namely: ComplexPortal,^63^ human DEPhOsphorylation Database (DEPOD),^64^ Gene Ontology,^65^ microT (miRBase and MirGeneDB versions),^66^ miRBase,^67^ NCBI Gene,^68^ PhosphoSitePlus,^69^ Signor,^70^ STRING^71^ and TarBase.^72^ Harmonization was conducted using a well-defined vocabulary of 69 edge labels, which express the biological effect associated with the edge (e.g. “gene *A* phosphorylates gene *B*”). The structure and standardized vocabulary of the Biology Mega Graph is recapitulated in Supplementary Table 1.

As the Biology Mega Graph aggregates mostly experiment-based information, a comparison between our predictions and Biology Mega Graph entries can be meaningful as long as the GOIs have sufficient literature coverage. For this reason, we chose to test the explainers on a set of 100 GOIs, selected for being the most annotated ones in terms of number of PubMed publications (a complete list can be found in Figure 8). The rationale was that, if the explainer proved effective at recognizing the interactors for these highly characterized genes, then, Proteomizer explainer will perform just as well on unannotated “ghost genes” too, for the simple reason that its training source (TCGA-CPTAC) is literature-agnostic.

First, we measured the alignment between the rankings computed by each explainer, and the known gene-gene or miRNA-gene annotations in the Biology Mega Graph. Alignment was quantified in terms of receiver operating characteristic area under the curve (ROC-AUC), which measures the general enrichment for the known gene-gene and miRNA-gene relations towards the top part of the ML-predicted ranking (i.e., it does not focus exclusively on the top-ranked items).

In the three direct tasks (1-3), Tx-based and Px-based helpers generally achieved AUCs in the 55-60% range, for example, “post-translationally modifies” yielded 55.9%, while “phosphorylates” obtained 56.6%. Excitingly, miRNA-based functions consistently reached AUCs of 70-75%. As for the explainer choice, XGBoost and Information Gain generally achieved the highest number of top performances in this task, without a clear winner (Supplementary Figure 9).

In the three reverse tasks (4-6), scores were generally higher, suggesting (unsurprisingly) that Px is more informative of the state of the tissue than Tx. Tx-based and Px-based helpers achieved AUCs of 60-65%, for example, “post-translationally modifies” hit 62.5%, whereas “phosphorylates” reached 63.9%. Once again, miRNA helpers outperformed the regular genes, with miRNA-based terms averaging remarkable AUCs of 75-80%. Among the three explainers, Information Gain secured most of the top performances in the Tx-based domain, whereas XGBoost was the best performer in the Px- and Mx tasks (Figure 6, Supplementary Figure 10).

**Figure 6.**
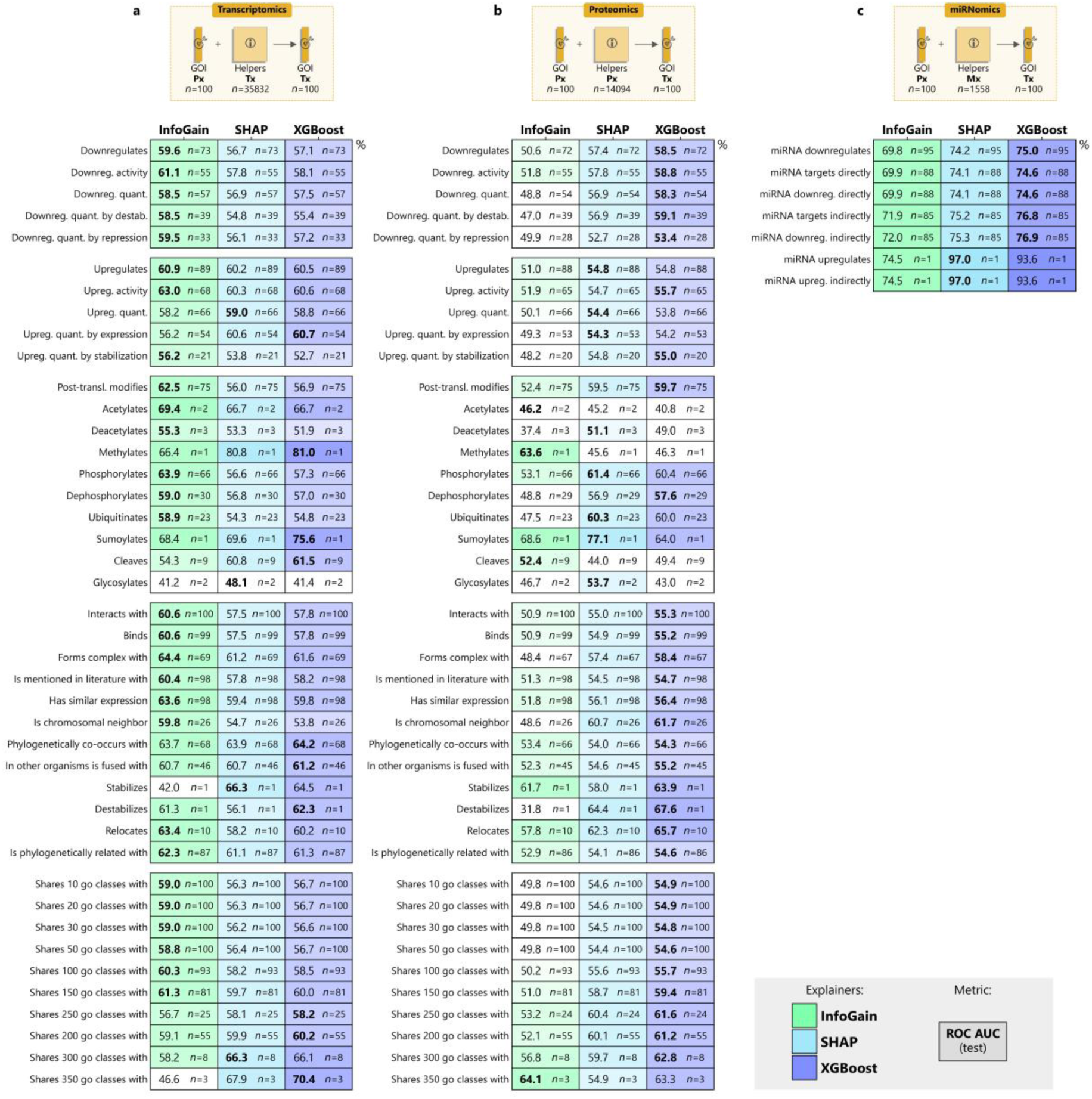
Proteomizer explainer ROC-AUC scores in the reverse tasks (proteome to transcriptome), benchmarked on the gene-gene and microRNA-gene relations from the Biology Mega Graph. The plot shows the receiver operating characteristic area under the curve (ROC-AUC) scores, obtained by comparing the importance scores from three Proteomizer explainers, against the known gene-gene and microRNA-gene relations in scientific literature. The Proteomizer importance scores were generated from a model, that was tasked to predict the RNA sequencing-based transcriptome (Tx) for 100 highly annotated genes of interest (GOIs), from their respective mass spectrometry-based proteome (Px) (reverse task), across 1,594 samples from 9 tissues of origin. Specifically, the importance score was determined for each of the helper genes (i.e., non-GOI genes) using three different explainers. (i) Information Gain (InfoGain, column 1) consists of training an independent eXtreme Gradient Boosting (XGBoost) model for each GOI-helper pair, where the GOI’s Px column and the helper column were provided as the input, and the GOI’s Tx as the target; then, taking the mean absolute error (MAE) across samples, with an inverted sign, as a score. (ii) XGBoost (column 2) consists of training an independent XGBoost model for each domain-helper pair, whereby the GOI’s Tx and all helpers in a domain—either whole Tx, whole Px, or whole miRNome (Mx)—were provided as the input, and GOI Px as the target; then, taking the XGBoost importance scores. (iii) SHapley Additive exPlanations (SHAP, column 3) is identical to (ii), but SHAP scores were used instead of the XGBoost importance ones. Finally, the importance scores from every explainer were compared against the known GOI-helper relations from literature, for each of 47 relation types, for each of the 100 GOIs. Such relations were sourced from the Biology Mega Graph (Supplementary Table 1), a knowledge graph which aggregates 320 million curated gene-gene and microRNA-gene relations from 11 biological repositories. The agreement between importance scores and literature was calculated in terms of ROC-AUC, in a direction-invariant way—e.g., the (*AKT*, *GSK3*) “phosphorylates” pair is treated as existing, if either AKT phosphorylates GSK3, or if GSK3 phosphorylates AKT. The top section of each table represents the task domain. (a) From the GOI’s Tx and each helper gene’s Tx, predicting the GOI’s Px. (b) From the GOI’s Tx and each helper gene’s Px, predicting the GOI’s Px. (c) From the GOI’s Tx and each helper miRNA’s Mx, predicting the GOI’s Px. The bottom section of each column expresses the ROC-AUC and sample numerosity for each of the 47 biological relation types, namely 42 gene-gene (a, b) and 7 miRNA-gene relations (c). The remaining terms were dropped due to insufficient number of examples. Numerosity (*n*) expresses how many of the 100 genes had at least one annotation of that corresponding relation type (e.g., if AKT phosphorylates some other proteins or is phosphorylated by some other proteins, then it will be counted as 1). The ROC-AUC of the best performing explainer for each task is shown in bold.

So far, we have shown that Proteomizer explainer is effective at yielding a *general* enrichment of known causal regulatory gene-gene and miRNA-gene relations. However, its intended final use is to prioritize only a few such regulators, as real-world biologists can realistically test in the lab only a few top hits (say, top 10-50). To assess this use case, we investigated if the explainers were capable of a major enrichment of known annotations, specifically in the top part of the ranking; that is, without taking into account the enrichment in the remainder part of the helper dataset. To do so, we measured the proportion of known GOI interactors among the explainer’s top lister helpers, versus the proportion of known GOI interactors genome-wide, for each of the 100 GOIs, and for each interaction modality in the Biology Mega Graph (Figure 7, Supplementary Figure 11). Most of the examined interaction modalities yielded an enrichment of 5-50× in the top 10 hits. For example, “upregulates” had a general prevalence of 0.06%, which increased to 3.5% in Proteomizer explainer’s top 10 hits (58×). Similarly, “post-translationally modifies” rose from 0.05% to 2.0% (40×); and “miRNA downregulates” rose from 3.04% to 17.3% (5.7×). In this task, SHAP outperformed the other architectures, suggesting it may be the preferrable choice for highlighting top-hit genes (as opposed to enriching the whole gene landscape).

**Figure 7.**
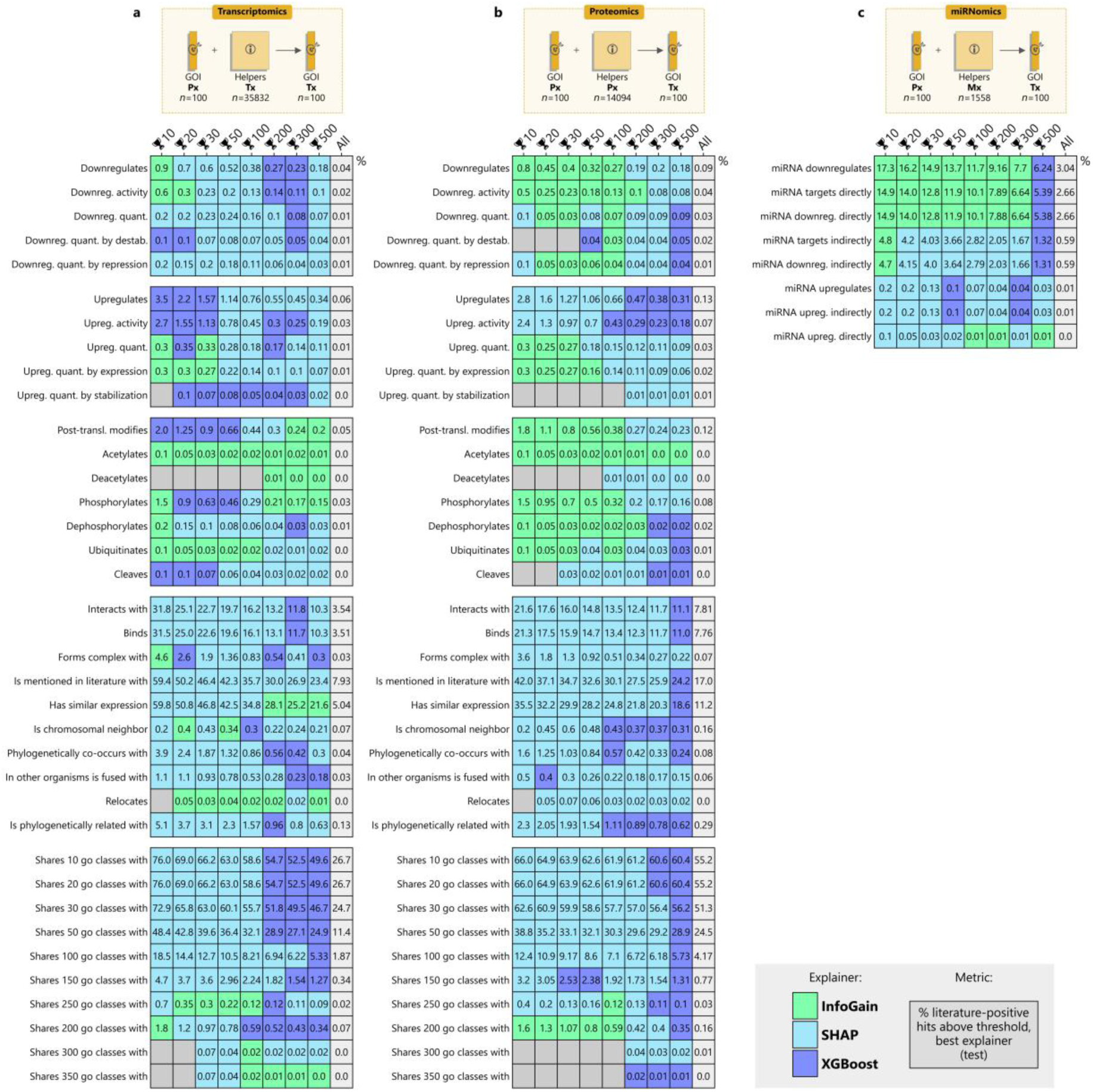
Proteomizer explainer top-hit positive fractions in the reverse tasks (proteome to transcriptome), benchmarked against the gene-gene and microRNA-gene relations in the Biology Mega Graph. The plot shows the fraction of top hits by three Proteomizer explainers, that are also known in the current scientific literature. The Proteomizer importance scores were generated from a model, that was tasked to predict the respective RNA sequencing-based transcriptome (Tx) of 100 highly annotated genes of interest (GOIs), from their mass spectrometry-based proteome (Px) (direct task), across 1,594 samples from 9 tissues of origin. Specifically, the importance score was determined for each of the helper genes (i.e., non-GOI genes) using three different explainers: Information Gain (InfoGain, green), eXtreme Gradient Boosting (XGBoost, blue), SHapley Additive exPlanations (SHAP, cyan)—see Figure 6 for details. Next, the importance scores were compared against the known GOI-helper relations from literature, for each of 47 relation types, for each of the 100 GOIs. Such relations were sourced from the Biology Mega Graph (Supplementary Table 1), a knowledge graph which aggregates 320 million curated gene-gene and miRNA-gene relations from 11 biological repositories. The agreement between the importance scores and literature were calculated in terms of top-hit positive fraction, in a direction-invariant way—e.g., the (*AKT*, *GSK3*) “phosphorylates” pair is treated as existing, if either AKT phosphorylates GSK3, or if GSK3 phosphorylates AKT. Essentially, for each of the GOIs, the explainer generated a raking of the helper genes; the ranking was cut at a variable threshold (🏆 *n*), starting from the top 5,000 and moving up to the top 10; this list of “predicted interactors” was compared with the list of known interactors from the Biology Mega Graph, and the percentage of genes in common between the two was counted (whereby 0% indicates that none of the top *n* hits are known interactors in a given biological modality, and 100% indicates that all of the top *n* genes are known interactors. The unthresholded percentage (All) is also displayed for reference, to highlight the enrichment observed among the top hits. The top section of each table represents the task domain. (a) From the GOI’s Px and each helper gene’s Tx, predicting the GOI’s Tx. (b) From the GOI’s Px and each helper gene’s Px, predicting the GOI’s Tx. (c) From the GOI’s Px and each helper miRNA’s Mx, predicting the GOI’s Tx. The bottom section of each table represents the top-hit positive fractions at varying thresholds. Specifically, the fraction was measured from the best-performing explainer, i.e., the one that yielded the highest fraction; the identity of such explainer varied on a case-by-case basis, and is expressed in a color code.

**Figure 8.**
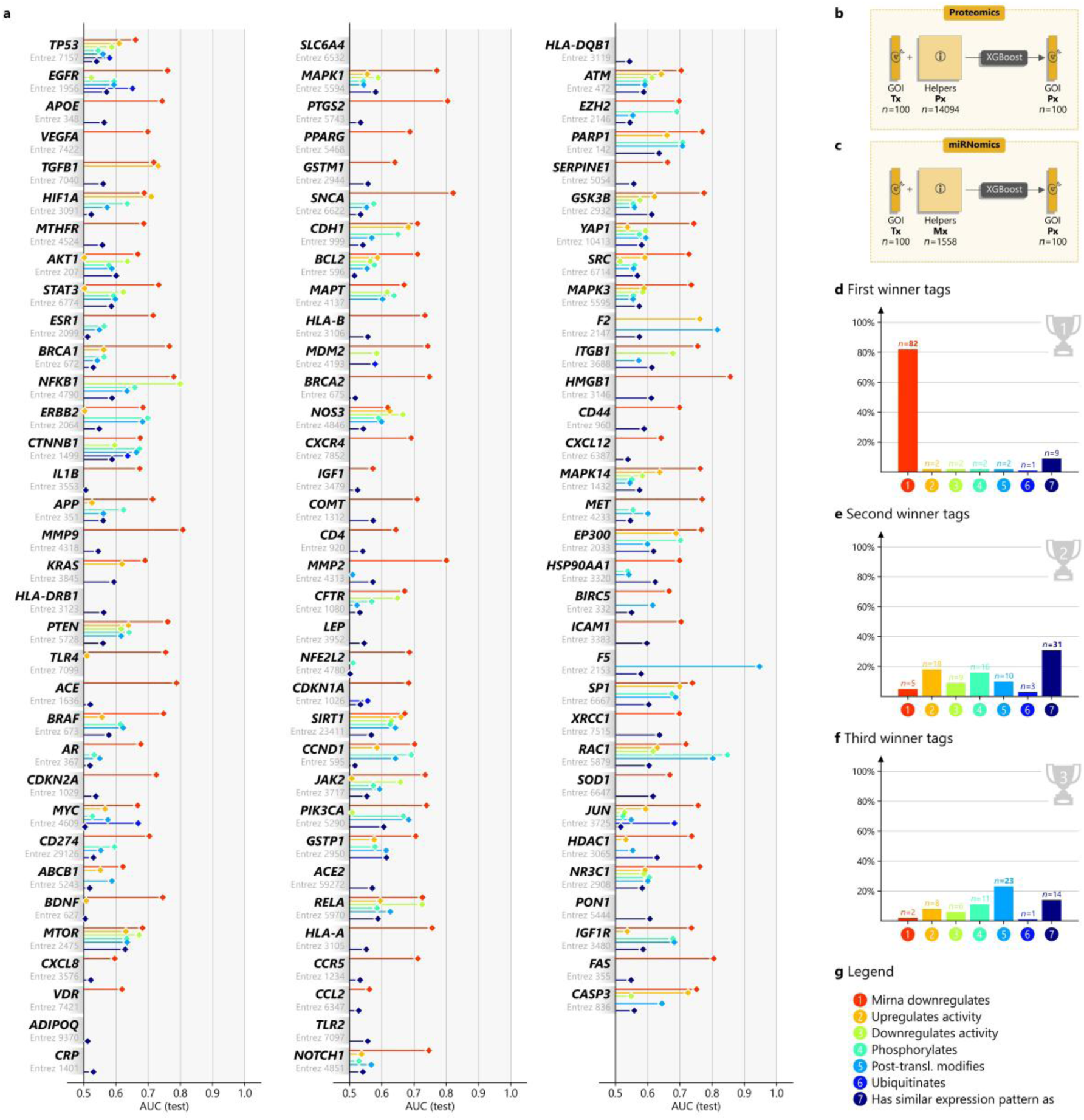
Proteomizer explainer XGBoost importance scores in the direct tasks (transcriptome to proteome), benchmarked against 7 relations in the Biology Mega Graph. The plot shows the single-gene detail of the receiver operating characteristic area under the curve (ROC-AUC) scores, obtained by comparing importance scores from three Proteomizer explainers, against 7 known gene-gene and miRNA-gene relation types in scientific literature, involving helper genes and microRNAs (miRNAs). Such relations types include 6 causal relation types, namely “mirna downregulates”, “upregulates activity”, “downregulates activity”, “phosphorylates”, “post-translational modifications”, “ubiquitinates”; and 1 correlational relation type, namely “has similar expression pattern as”. The Proteomizer importance scores were generated from a model, that was tasked to predict the mass spectrometry-based proteome (Px) of 100 highly annotated genes of interest (GOIs), from their respective RNA sequencing-based transcriptome (Tx) (direct task), across 1,594 samples from 9 tissues of origin. Specifically, the importance score was determined for each of the helper genes (i.e., non-GOI genes) using the eXtreme Gradient Boosting (XGBoost) explainer—see Figure 6 for details. Next, the importance scores were compared against the known GOI-helper relations from literature, for each of selected 7 relation types, for each of the 100 GOIs. Such relations were sourced from the Biology Mega Graph (Supplementary Table 1), a knowledge graph which aggregates 320 million curated gene-gene and miRNA-gene relations from 11 biological repositories. The agreement between importance scores and literature were calculated in terms of ROC-AUC, in a direction-invariant way—e.g., the (*AKT*, *GSK3*) “phosphorylates” pair is treated as existing, if either AKT phosphorylates GSK3, or if GSK3 phosphorylates AKT. (a) 100 ROC-AUC plots, referred to each of the 100 GOIs; bars compare the enrichment of the 7 relation types, for the Proteomizer XGBoost explainer predictions referred to each GOI. Please observe that the x-axis scale goes from 0.5 to 1.0 (AUCs below 0.5 are not shown). (b, c) Task overview: the 7 selected relation predictions involve 6 tasks of the type “from the GOI Tx and each helper gene Tx, predicting the GOI Px”. The only exception is “mirna downregulates”, which is of the type “from the GOI Tx and each helper miRNA Mx, predicting the GOI Px”. (d-f) Winner plot, depicting which relation type achieved the highest ROC-AUC in a given GOI, for all the GOIs (the maximum being 100); below, which relation type achieved the second-highest ROC-AUC, and third-highest ROC-AUC. (g) Color code for 7 relation prediction tasks. We can observe that “mirna downregulates” was ranked as the most enriched task for 82 of the 100 GOIs, suggesting that Proteomizer explainer is remarkably effective at picking up miRNAs. A possible criticism to Proteomizer explainer is the fact that it may pick up genes that are simply coregulated with the GOI. For this reason, the correlational relation type “has similar expression pattern as” was included in the assessment: while it ranked #2 overall, it did so with a great margin from “mirna downregulates”, achieving only 9 out of 100 first places; additionally, in the second-ranked winners, the other causative relations achieved comparable levels to the correlational relation type.

Finally, we wished to assess the overall contribution of Tx, Px and Mx to the prediction of the GOIs’ mismatch. To pursue this question, we trained an XGBoost model for each of the 100 GOIs, comparing the following helper dataset combinations: Tx, Px, Mx, Tx+Px, Tx+Mx, Px+Mx, Tx+Mx+Px (Table 4). Without ML, the mean GOI Tx-Px correlation was *ρ* = 0.346. The ML from the Tx domain increased this figure to 0.579; Mx yielded a comparable 0.559; while Px boosted it to a whopping 0.778, implying that Px is the most informative of the three omics. It is somewhat surprising that the 1,558 miRNAs conveyed as much information on the GOIs’ mismatch as the 34,322 Tx genes; additionally, Tx and Mx conveyed overlapping information as no synergy was observed between the two of them, considering that Tx+Mx yielded a mere *ρ* = 0.580 (almost identical to that of the two channels taken separately). Overall, these results provide a harmonized measurement of how much each of the three omics is informative regarding the Tx-Px mismatch in general, and suggest that Mx contains a high “density” of information regarding the Tx-Px mismatch.

**Table 4.**
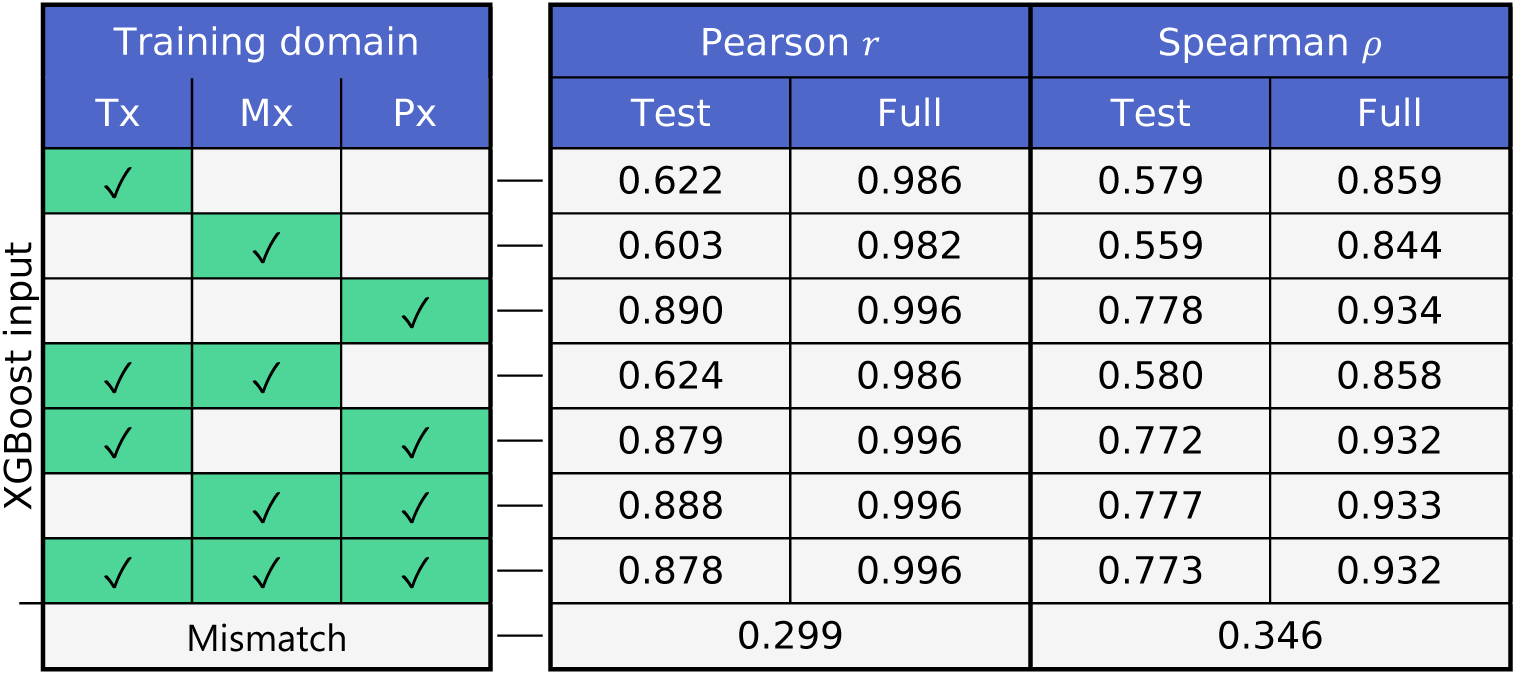
Performance from Proteomizer XGBoost explainer as a function of the input domain included in the training, in the direct tasks (transcriptome to proteome) An eXtreme Gradient Boosting (XGBoost) model was tasked to predict the mass spectrometry-based proteome (Px) of 100 highly annotated genes of interest (GOIs), from their respective RNA sequencing-based transcriptome (Tx) (direct task), across 1,594 samples from 9 tissues of origin. Alongside the GOI’s Tx, one or more genome-wide helper domains were passed as input: Tx, miRNomics (Mx), Px, the combination of Tx and Mx, the combination of Tx and Px, the combination of Mx and Px, and finally, all of the domains combined. The training was repeated individually for each of the 100 GOI, across the 7 different input epochs (rows). Performance was quantified in terms of Tx-Px correlation across the samples, averaged for all 100 GOIs. Specifically, four metrics were used: Pearson correlation coefficient *r* using the prediction from the test dataset (test), or from the entire dataset(full); Spearman correlation coefficient *ρ* using the prediction from the test dataset (test), or from the entire dataset(full). The raw Tx-Px mismatch is displayed in the bottom raw. Of note, this analysis was carried out while excluding all microRNAs (miRNAs) from the Tx: as explained in the Discussion, miRNAs are generally picked up by standard Tx, though with lower accuracy than Mx; in order to measure the contribution of messenger RNAs (mRNAs) separately from miRNAs, the miRNAs readouts in Tx were dropped, reducing the Tx reads from 35,833 down to 34,322. Overall, Px appears to be by far the most informative source of information to predict a GOI’s Tx, from its respective Px. Interestingly, the contribution from is substantially similar to that of Mx, despite the number of reads being tremendously different (34,322 and 1,558 reads, respectively). Also, Tx and Mx information is not synergistic, implying that the same information on the cell’s state is somehow shared between the two of them.

**Table 5.**
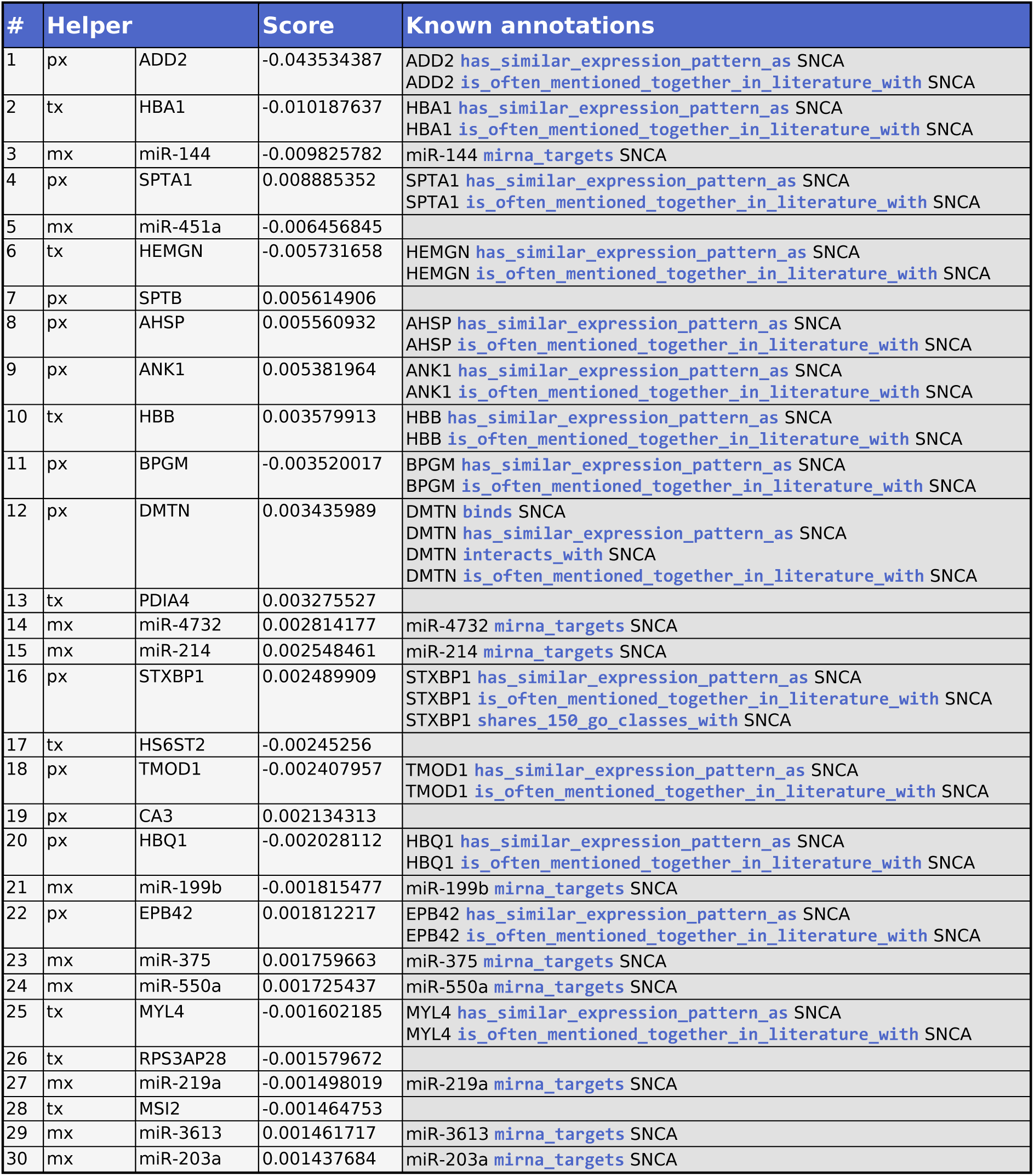
Proteomizer explainer results for alpha-synuclein, using the SHAP explainer architecture in the direct tasks (transcriptome to proteome) The table lists the helper genes that resulted to be the most informative at predicting the mass spectrometry-based proteomic levels of alpha-synuclein (*SNCA* gene), given its matched RNA sequencing-based transcriptomic levels across 1,594 samples from 9 tissues of origin, according to the SHapley Additive exPlanations (SHAP) explainer. The “#” column represents the helper gene’s rank, based on the absolute SHAP score. The “Helper” column expresses the helper gene’s domain (tx: transcriptomics; mx: miRNomics; px: proteomics) and standardized gene symbol. The “Score” column expresses the SHAP signed score. The “Known annotations” list the already known interactions between the helper gene and alpha-synuclein, in addition to their modality, as they appear in the Biology Mega Graph.

**Table 6.**
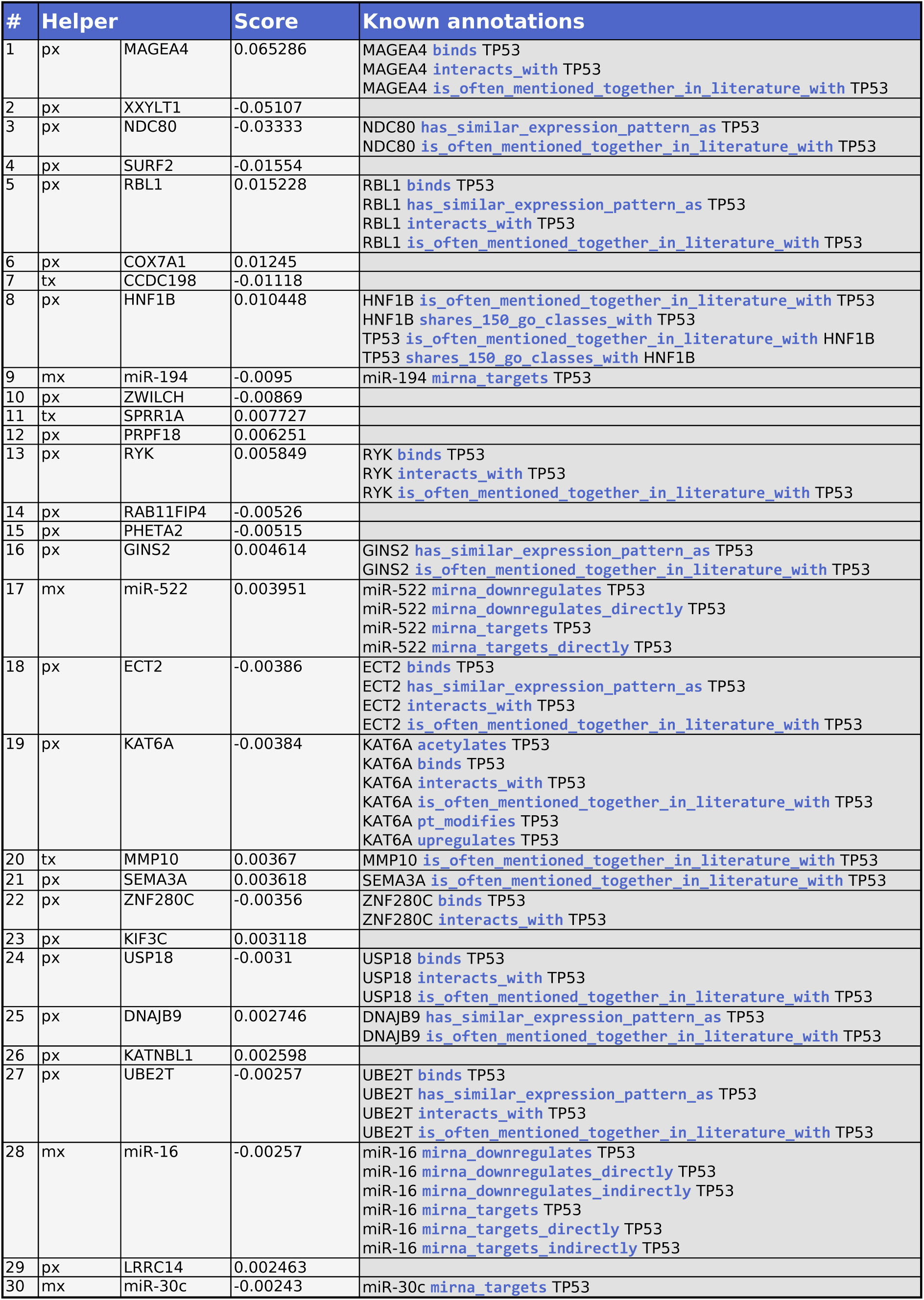
Proteomizer explainer results for p53, using the SHAP architecture in the direct tasks (transcriptome to proteome) The table lists the helper genes that resulted to be the most informative at predicting the mass spectrometry-based proteomic levels of p53 (*TP53* gene), given its matched RNA sequencing-based transcriptomic levels across 1,594 samples from 9 tissues of origin, according to the SHapley Additive exPlanations (SHAP) explainer. The “#” column represents the helper gene’s rank, based on the absolute SHAP score. The “Helper” column expresses the helper gene’s domain (tx: transcriptomics; mx: miRNomics; px: proteomics) and standardized gene symbol. The “Score” column expresses the SHAP signed score. The “Known annotations” list the already known interactions between the helper gene and p53, in addition to their modality, as they appear in the Biology Mega Graph.

**Table 7.** Performance quantification of Proteomizer, compared to existing proteomization architectures. The table offers a comparison between Proteomizer and all the main proteomization architecture available in literature, with respect to their performance and machine learning approach. Performance is quantified using correlation coefficients, specifically the Pearson *r* and the Spearman *ρ*, calculated across the samples and averaged for all genes. The arrow (“➤”) separates the before-after values: on the left side, the correlation value in the raw dataset is reported, i.e., the correlation between the raw transcriptome (Tx) and the ground-truth proteome (Px); on the right side, the correlation value in the prediction is reported, i.e., the correlation between the predicted proteome (Px’) and the ground-truth Px. Papers are sorted chronologically by their publication date. The “Input” column showcases which datasets were utilized to train the model, and their sample size (*n*); datasets include The Cancer Genome Atlas (TCGA), Clinical Proteomic Tumor Analysis Consortium (CPTAC) and the Religious Orders Study/Memory and Aging Project (ROSMAP); for Proteomizer, the mass spectrometry version is assumed as the reference. Finally, the “Model” column highlights which regressor architecture was employed to make the predictions.

| Date | Publication | Pearson $r$<br>(samples) | Sparman $\rho$<br>(samples) | Input | Model |
| --- | --- | --- | --- | --- | --- |
| 2019.12 | Li et al. | 0.38 ► 0.53 | N/A | TCGA-CPTAC<br>($n = 182$) | Random forest |
| 2020.02 | Magnusson et al. | 0.21 ► 0.79-<br>0.94 | N/A | CD4<sup>+</sup> T-helper<br>cells ($n = 7$) | Linear model |
| 2022.02 | Tasaki et al. | 0.08 ► 0.18 | N/A | ROSMAP<br>($n = 384$) | Variational<br>autoencoder (VAE) |
| 2022.11 | Srivastava et al. | 0.47 ► 0.60 | N/A | TCGA-CPTAC<br>($n = 958$) | Linear regression,<br>elastic net,<br>random forest |
| 2024.07 | Cranney and Meyer | N/A | N/A | TCGA-CPTAC<br>($n = 1,256$) | Deep learning |
| 2025.06 | Proteomizer | 0.27 ► 0.68 | 0.31 ► 0.54 | TCGA-CPTAC<br>($n = 1,594$) | Deep learning |

### PROTEOMIZER EXPLAINER is particularly effective at identifying micro-RNAs targeting a given gene of interest

One could argue that Proteomizer explainer may theoretically pick up both causal and coregulation relations. Among the 49 terms explored, while the majority represented causal biological relations (e.g., “post-translationally modifies”, “phosphorylates”, “upregulates”, “relocates, interacts with”, etc.), 4 terms were included that expressed coregulatory, non-causal relations (“has similar expression”, “is chromosomal neighbor”, “phylogenetically co-occurs with”, “in other organisms is fused with”). Remarkably, the AUCs from the best-performing causal relations such as “mirna downregulates” (72-77%) were markedly higher than those from the 4 coregulation terms (60-65%). This was true for both direct and reverse tasks. It suggests that, although some coregulation may be picked up by the Proteomizer explainers as a confounding variable, its effect size does not prevail over the causal relations, most notably the miRNA-related ones. These results generally confirm that Proteomizer explainer is an effective tool to highlight the genes—and especially the miRNAs—that play a role in the regulation of a GOI.

Next, we opted to assess, at single-GOI resolution, which biological terms were the most predictive of its Tx-Px mismatch, among a selection of 6 causal terms (“mirna downregulates”, “interacts with”, “post-translational modifications”, “phosphorylates”, “ubiquitinates”, “relocates”) and 1 coregulation term (“has similar expression pattern as”). Remarkably, the most predictive term turned out to be “mirna downregulates” in 82 out of 100 GOIs, whereas “has similar expression pattern as” won in only 9 cases. Conversely, the second-most predictive term was “has similar expression pattern as” in 31 of 100 GOIs, with the causal terms covering the remainder 69 GOIs in somewhat comparable proportions. These results further confirm that Proteomizer explainer is an effective approach to pick up miRNA regulators for a given GOI, and is only moderately influenced by coregulatory epiphenomena.

### PROTEOMIZER EXPLAINER can be applied to highlight novel regulators of a given gene of interest

So far, we have provided computational evidence that Proteomizer explainer is capable of identifying a GOI’s regulatory partners, with only a minority of the hits being due to epiphenomenal coregulation. Furthermore, its literature-agnostic architecture makes it ideal to identify novel, uncharacterized regulatory relations. To validate such hypothesis, let’s examine some representative predictions for two GOIs of clinical significance. In light of the different explainers’ performance in top hit prioritization, the SHAP explainer will be employed.

As a first example GOI, we will consider alpha-synuclein (αS, *SNCA* gene), regarded as the central protein of Parkinson’s disease.^73,74^ Among its 30 top-ranked interactors predicted by Proteomizer explainer, 23 had already been annotated to share at least one type of interaction with αS according to the Biology Mega Graph, whereas 7 appeared novel. These include the following:

- *miR-451a* (hit #5) has been reported as a differentially expressed gene in multiple transcriptomic experiments on neurodegeneration, including serum from early-stage alpha-synucleinopathies,^75^ leukocytes from amyotrophic lateral sclerosis,^76^ liquor from Alzheimer’s disease^77^ and brain from Huntington’s disease.^78^ While a specific action on αS has never been reported (nor excluded), this represents a favorable substrate for such hypothesis, due the extensive mechanistic similarities among neurodegenerative diseases.
- *SPTB* (hit #7) encodes a subunit of spectrins, cytoskeletal components involved in membrane organization. Spectrins have been variously associated with neurodegeneration and psychiatric disorders. Notably, the pathology-specific phosphorylated species of αS was reported to bind spectrins and disrupt their function in synaptic organization: this mechanism was hypothesized— among many others—as a possible explanation for αS-mediated toxicity.^79^
- *PDIA4* (hit #13) encodes a protein disulfide isomerase, a class of endoplasmic reticulum chaperones involved in protein folding. Failure of the protein folding system and subsequent endoplasmic reticulum stress are a core mechanism in neurodegeneration; while disulfide isomerases have been repeatedly implicated as protective to PD, existing PD studies seem to have focused on *PDIA1* and *PDIA3*, while *PDIA4* has been engaged more by the cancer community, and appears understudied in the context of neurodegeneration.^80^

As an alternative example, let’s consider p53 (*TP53* gene), which was the first oncosuppressor gene to be identified, in 1979. It acts as a downstream endpoint for numerous cell-cycle-suppressor pathways, representing a variety of biological mechanisms ranging from DNA damage to hypoxia, oxidative stress and many others. Being the most frequently mutated gene in human cancer (∼50%), and its mutation being a marker for worse prognosis, it has been nicknamed the “holy grail” of cancer.^81,82^ Among the top 30 hits according to Proteomizer explainer, 18 had already been associated with *TP53* in the Biology Mega Grah. The top novel ones are the following:

- *XXYLT1* (hit #2) was originally characterized in 2012 as a UDP-xylose:α-xyloside α1,3-xylosyltransferase.^83^ Subsequent evidence highlighted that its deletion promotes lung cancer by inducing cell proliferation while inhibiting apoptosis^84^; and that its mRNA levels are downregulated, while methylation levels are upregulated, in lung adenocarcinoma over normal lung tissue. Although a direct nexus with p53 has never been highlighted, the high percentage of p53 mutations in lung adenocarcinoma (also around 50%^82^) implies that a downstream involvement is very likely.
- *SURF2* (hit #4) was functionally characterized only in 2024, as it was found to be a physical interactor of free 5S ribosomal subunit; ribosomes have been pointed out as a cancer chemotherapy target, because their downregulation leads to nucleolar stress, which in turn triggers p53-mediated cell cycle arrest. The study found that SURF2 inhibits MDM2, which is a known, major inhibitor of p53.^85^
- *COX7A1* (hit #6) encodes a subunit of mitochondrial complex IV. Its overexpression blocks cell proliferation in lung cancer cells by impairing autophagy.^86^ Additionally, it has been linked to lung cancer’s sensitivity to cysteine deprivation, suggesting a possible role as a therapeutic target.^87^
- *CCDC198* (hit #7) was only characterized in 2023, and renamed as *FAME*. Its main role was reported to be energy balance and iron metabolism. Notably, yeast-two-hybrid screen revealed physical interaction with known cell-cycle regulators, acting on MDM2, and thereby influencing p53 levels; for this reason, the authors postulated a role in cell cycle regulation.^88^

Very similar considerations apply to the rest of the novel hits (*ZWILCH*, *SPRR1A*, *PRPF18*, *RAB11FIP4*, *PHETA2*, *KIF3C*, *KATNBL1*, *LRRC14*). Overall, the top hits for both example GOIs appear enriched with known interactors for the given GOI—which is expected, considering that *SNCA* and *TP53* are two of the most widely studied genes in the history of biology. Interestingly though, these known hits are intermingled with novel predicted regulators, many of which show robust theoretical grounds, that corroborate a hypothetical regulatory role upon the GOI. These results suggest that Proteomizer explainer is indeed effective at detecting novel interaction or regulatory partners for a given GOI, justifying its usefulness for today’s real-world biologists.

## Discussion

In the past 20 years, gene expression analysis by Tx has become one of the most widely adopted tools in biologists’ arsenal. It allows to reliably quantify mRNA levels across all genes in a sample, enabling investigators to make differential comparisons across two or more conditions—be them treated-untreated, healthy-diseased, and so forth. However, this idea comes with a fundamental flaw: for most biologically active genes, it is the proteins (not the RNAs) that act out biological functions. Still, when biologists deploy Tx, they are implicitly utilizing mRNAs as a proxy for protein quantity measurement. While the latter could be measured directly through Px, this is rarely done due to its higher cost and complexity over Tx. For example, when a researcher establishes that protein-folding-related genes are upregulated in Parkinson’s disease substantia nigra, they usually do so based on mRNA data, *implying* that those markers must be elevated also at protein level. ^89^ After all, one may think: if a gene is upregulated at mRNA level, those extra mRNA copies will *obviously* result in a higher number of protein copies—it’s the central dogma of Molecular Biology.^90^ Regrettably, this ground assumption proves incorrect. The few papers that quantified Tx-Px correlation reported a humble *r* = 0.4-0.7 across genes, and a ghastly *r* = 0.1-0.5 across different samples. Indeed, the existence of such mismatch is justified by a large number of evolutionary, biological and to a much smaller extent technical justifications, as previously discussed.

The increasing availability of matched Tx and Px datasets has prompted the bioinformatic community to explore whether a sample’s Px landscape may be inferred from information that is diffusely represented in the Tx landscape; for this task, we coined the neologism of “proteomizing” the transcriptome. A few past bioinformatic papers have attempted this endeavor. For the most part, they utilized shallow learning techniques and a very limited number of samples (<200);^11,35,91,92^ only very recently, three papers overcame such limitations, namely Srivastava *et al*. (2022),^2^ Tasaki *et al.* (2022)^3^ and Cranney and Meyer (2024).^46^ However, even these studies came with significant limitations: most crucially, while most of them provided a performance characterization in terms of ML metrics, the biological characterization was largely missing. As a result, their effectiveness on *practical* biological use cases was never investigated.

In the present work, we introduced PROTEOMIZER, a novel, deep-learning based tool that not only outperforms the previously published ML papers in the Tx-to-Px task, but much more importantly, comes with data-driven assessment of its effectiveness in two real-world biological applications: (i) differential expression analysis and (ii) the identification causal regulators of a given GOI, most notably, miRNAs. In the next paragraphs, we will briefly discuss Proteomizer’s strengths and limits, in various respects.

Let’s start from the dataset choice: despite impressive progress in the past 5 years, availability of matched Tx-Px data is still extremely limited, and to the best of our knowledge, the only large-scale, multi-organ dataset of the kind is TCGA-CPTAC. While its existence is an invaluable asset to the bioinformatic community, it is important to mention the limitations that come with such dataset, for our specific purposes. First, all measurements were conducted in bulk tissue, as single-cell proteomics still remains an ongoing frontier;^7^ this implies that cell type proportions may vary from one sample to another, for example, as a result of aging^93^ or inflammation, adding a layer of noise to across all measurements. Second, 99.6% of the RPPA samples and 77.2% of the MS samples were of cancerous nature, implying the widespread presence of driver and acquired genomic mutations, which may alter the functioning of proteostasis (e.g., reducing the efficacy of miRNA-mediated translation repression^94^); these can also include large-scale chromosomal rearrangements which, in extreme cases, may grossly modify the cell functioning^95^ (a notorious example being HeLa cells, showcasing 80-110 chromosomes^96^). Finally, while TMT offers one of the largest dynamic intervals of all Px approaches,^97^ it is still impacted by reproducibility limits, even in state-of-the-art facilities; for example, 32 CPTAC ovarian samples that were processed at Pacific Northwest National Laboratory, and independently re-processed at Johns Hopkins University barely correlated, at a Pearson score of 0.59.^11^

The omic choice also deserves some attention: Proteomizer was trained from Tx and Mx. Differently from the CPTAC DREAM challenge,^41^ we did not include genomics information, that is, the cancer-related mutations that were present in each sample. This choice was made deliberately, and consistently with most papers in this space,^2–4,46^ as the scope of Proteomizer is not specific to cancer biology, but rather, it is meant as a general-purpose tool that maps Tx into Px. However, the presence of mutated genes in our training samples indeed acts as a confounding factor, and the addition of genomics may prove valuable as a future expansion, especially for the explainer part. Conversely, a crucial (and so far, unprecedented) strength of Proteomizer is the inclusion of Mx. Most other proteomization papers studied miRNAs from their Tx readouts,^46^ which is questionable: Tx protocols are optimized to selectively pick up mRNAs (not miRNAs), leveraging the fact that only mRNAs are capped by the 5’ 7-methylguanylate cap.^98^ Likewise in Tx, library preparation and sequencing depth are optimized for RNA species much longer than miRNAs (which are only 18-25 nucleotide-long). For these reasons, miRNAs should be detected using a specific miRNA-sequencing protocol (i.e., Mx), which was provided among the many multiomic assays in TCGA.

Moving on to the ML metrics, Proteomizer achieved the best performance available to this day in a multi-tissue proteomization task (Table 4) by a remarkable margin, scoring 0.68 in Pearson *r* and 0.54 in Spearman *ρ* correlation coefficients (all data are referred to the MS version, in the test dataset). The closest example is that of Srivastava *et al.*, which achieved a Pearson value across samples of 0.599.^2^ Two noticeable factors motivate such gap: on one hand, the increasing availability of training examples; on the other, the relatively simple models employed by most other papers, relying mostly on linear models up to as recently as 2022.^2,4^ Of course, even Proteomizer’s *ρ* = 0.68 is still a far cry from perfect correlation. However, incomplete predictability is easy to justify biologically, as only a subset of genes are thought to have feedback loops, conveying their protein abundance information back into the Tx landscape—which is the fundamental assumption underlying the proteomization task.

Architecturally speaking, Proteomizer’s deep neural network may be further developed. For example, considering the high amount of noise affecting Tx and Px data, a denoiser may be beneficial (although one could argue that the existing implementation, coupling an excess in model complexity with early stop to avert overfitting, should effectively behave as a denoiser itself). Alternatively, an encoder-decoder structure could be beneficial to distil a low-dimensionality, high-expressivity embedding of the tissue state, which may boost performance. On the other hand, both of these tools were employed by Tasaki *et al.* on their AD Knowledge Portal-based predictions, and still, their performance was limited to *r* = 0.18.^3^ Overall, these data suggest that training data play a much heavier role than the architectural choice, consistently with our observed marginal performance wiggle at varying hyperparameter combinations, and with an ongoing trend in the ML community calling for more “data-driven AI”.

Further to the prediction score itself, an important advancement in Proteomizer lies in the Monte Carlo method to validate its prediction. Previous papers generally did not assess whether the increase in correlation coefficients actually brought along improved accuracy in identifying DEGs. Notable exceptions were Magnusson *et al.*, who focused on time-resolved DEGs during Th1 cell differentiation^4^; and Tasaki *et al.*, who reported that the DEGs from the proteomized dataset matched more closely those from the Px, although only for the specific case of AD versus control, and with unsatisfactory performance.^3^ Proteomizer represents the first instance where proteomization benefit was systematically measured in terms of DEG detection against the Px ground truth, showcasing a p-value boost between 1.2× and 12.9× across all genes, and as large as 3 to 6 orders of magnitude in the top 5% of the genes.

As regards gene set enrichment, we reported the highest raw Tx-Px correlation in structural genes, and the worst correlation in metabolism-related functions; after proteomization, the best correlation was in ribosomal, cytoskeletal and cell respiration terms, whereas the worst was in ubiquitin and transfer RNAs (tRNAs); finally, the maximum benefit was in aerobic respiration terms, while the least benefit was observed for transitory responses such as the inflammatory. This is partially in line with existing literature, barring a certain degree of inconsistency among the various papers. Early findings on *S. cerevisiae* highlighted metabolism, energy and protein synthesis among the best correlated (least mismatched) functions.^99^ Conversely, CPTAC-based Li *et al.* showed that, after proteomization, metabolism displayed the highest correlation scores (consistently with Proteomizer), whereas the minimum ones belonged to ribosome, spliceosome, proteasome, and neurodegeneration classes; the maximum benefit was observed for metabolism (consistently with Proteomizer).^11^ Conversely, CPTAC-based Srivastava *et al.* reported the highest raw correlations for membrane proteins and cell junctions, and the poorest in large multi-protein complexes.^2^ This partial lack of agreement may be due to different tissues being used for each study, different versions of GO annotations being used,^100^ and to the widespread practice of off-the-shelf enrichment algorithms. Despite being quick and convenient, these usually do not ensure that the universe of the provided data is identical to the one used in the ontology: errors can arise due to Entrez ID versioning, symbol ambiguity and improper naming conversions. Proteomizer uses a custom enrichment algorithm which enforces the highest standards of rigorousness in this respect.

While Proteomizer was remarkably successful at improving correlation scores and DEG detection in tissue types that had been included among the training examples, it failed to do so on novel tissues, be them from the same CPTAC repository, or from the independent AD Knowledge Portal source. The general concept of training a proteomization architecture on one dataset while tasking it to make predictions on a different one has been attempted before by Magnusson *et al.* and Tasaki *et al*., yielding mostly negative results.^3,4^ Magnusson *et al.* trained one of their linear models on time-resolved, splicing-variant-resolved datasets of Th1, T-reg and B lymphocytes; later, they tried to apply their predictors to different tissues from the human protein atlas,^101^ reporting marginal improvements. They concluded that “the model should be applied in a cell-specific manner given the low correlation in mixed tissue samples”.^4^ Instead, Tasaki *et al.* trained their model on the ROSMAP study, and later applied the predictor to the MSBB one. Both of these studies combine healthy and AD aged brains, though they were carried out by independent groups, using different brain regions, equipment, experimental protocol and data processing. On top of that, both aging^93^ and neurodegeneration^102^ are known for affecting proteostasis, further exacerbating the Tx-Px mismatch issue and making it more unpredictable. Again, their model only marginally increased MSBB’s correlation, from 0.09 to 0.14.^3^ The most likely explanation is that this is an intrinsic limitation of the technique: proteomization can only be effective as long as the model has seen examples of the tissue type in question during training, and the experimental protocol is consistent between the training dataset and the dataset used to make predictions. As a future development, it would be interesting to check if this limitation may be overridden by creating a new ML architecture that takes as input a “supplement of information” regarding the natural state of the tissue in question.

Finally, we employed three explainer architectures to detect which mRNAs, proteins and miRNAs were the most informative to predict the Tx-Px mismatch for 100 highly annotated GOIs. The idea of using XAI on proteomization tasks is not a new one. Indeed, most of the mentioned papers applied either random forest feature importance scores^2^, SHAP^2,3,46^ or Boruta^2^. However, none provided a biological validation for their predictions: they restricted their efforts to merely presenting the explainer results, without assessing their reliability or novelty. For example, Srivastava *et al.* utilized the importance scores to create subgraphs centered around each GOI; however, they restricted the scope of their explainer to the proteins which were *already* known for being interactors of each GOI, based on either CORUM or STRING. As a result, such results could not be utilized to discover *novel* gene-gene regulatory pathways.^2^

Importantly, none of the other papers quantified the relative importance of causal relations, compared to mere correlations. Conversely, our analyses compared 26 causal relations (e.g., “post-translationally modifies”, “phosphorylates”, “upregulates”, “relocates”, etc.) against 4 correlative relations (“has similar expression”, “is chromosomal neighbor”, “phylogenetically co-occurs with”, “in other organisms is fused with”), and showed that the best-performing causal terms generally outperformed the correlative ones. This result confirms that, although indeed some coregulated genes are picked up by the analysis, their presence is quantitatively limited in favor of the cause-effect biological relations. Proteomizer explainer proved remarkably proficient at predicting miRNA-gene relations, reaching a ROC AUC of 0.75. Provided the remarkable biological relevance of miRNAs in clinically relevant contexts,^103^ this positions Proteomizer explainer as a potential source for the discovery of novel miRNAs, especially for ghost genes.

As a final remark, we would like to stress how different (and more promising) this investigation approach is, compared to more traditional methods like coexpression. The latter is based on the guilt-by-association (GBA) assumption, by which, if two genes are coregulated, they are (slightly) more likely to share some of their biological functions.^104^ Conversely, our method measures the information contribution that each gene or miRNA conveys to explain the mismatch between the GOI’s Tx and Px readouts, thus focusing exclusively on what happens at post-transcriptomic level. For this reason, Proteomizer explainer is particularly effective at picking up novel, causal actions between genes, in a completely literature-agnostic fashion.

## Materials and methods

### Virtual Lab

Proteomizer was created using our Virtual Lab (VLab) ML framework.^62^ It was developed in Python 3.9.19, using the Anaconda v23.7.4 distribution to manage libraries and environments, and the PyCharm 2022.2.5 (Professional Edition) environment. The ML code relied on the PyTorch (torch) and Scikit-learn (sklearn) frameworks. All training was conducted using Nvidia CUDA^105^ v12.3.2 to leverage GPU acceleration for computational efficiency.

### TCGA-CPTAC datasets

The Proteomizer training dataset was sourced from The Cancer Genome Atlas (TCGA)^37^ and Clinical Proteomic Tumor Analysis Consortium (CPTAC).^38,39^ Specifically, the RNA sequencing (RNAseq), microRNA sequencing (miRNAseq) and RPPA data were accessed through the NIH Genomic Data Commons (GDC) data portal (portal.gdc.cancer.gov), version 2.^47^ Files were filtered by the “experimental strategy” field, selecting “RNA-Seq”, “miRNA-Seq” and “RPPA”, respectively; and by the “Data Type” field, selecting “Gene Expression Quantification”, “miRNA Expression Quantification” and “Protein Expression Quantification”, respectively. The MS data were accessed through the NIH Proteomic Data Commons (PDC) data portal (proteomic.datacommons.cancer.gov/pdc/).^48^ Files were selected by the “Data Category” field, selecting “Protein Assembly”, and by the “Analytical Fraction” field, selecting “Proteome”. From both file selections, a File manifest, Biospecimens manifest, and a Clinical manifest were generated. The number of retrieved samples was the following: 25,192 for RNAseq, 17,469 for miRNAseq, 7,906 for RPPA, 5,228 MS (September 2024).

Samples were intersected across the different channels, in terms of their “Sample ID” field: the intersection of RNAseq ∩ miRNAseq ∩ RPPA yielded 7,338 sample IDs (version 1); the intersection of RNAseq ∩ miRNAseq ∩ MS yielded 1,957 sample IDs (version 2). Duplicates (0.28%) were dropped. For both versions, data corresponding to the selected sample IDs were downloaded using the GDC data transfer tool^106^; for PDC, a Python script downloaded each file by the HTTP link provided in the file manifest, using the Python requests library. A dataframe was assembled from each downloaded file: for RNAseq, the tpm_unstranded column was used (in line with established transcriptomics guidelines, suggesting to utilize transcripts per million mapped reads, TPM^107,108^); for miRNAseq, the reads_per_million_miRNA_mapped column was used; for RPPA, the protein_expression column was used; for MS, the Log Ratio columns were used for each of the reporter ions (“126C”, “127N”, “127C”, “128N”, “128C”, “129N”, “129C”, “130N”, “130C”, “131N”, “131C”, excluding the POOL).

Next, namespace conversions were applied:

- RNAseq data were originally expressed as 60,660 Ensembl IDs; these were demultiplexed into 39,467 Entrez IDs; synonymous IDs were aggregated by sum; all-0 reads were dropped, resulting in 35,979 approved reads in version 1, and 35,833 approved reads in version 2.
- miRNAseq data were originally expressed as 1,877 miRNA names; names were mapped to the namespace of miRBase^67^; 31 unmatched names were dropped; synonymous names were aggregated by sum, and all-0 reads were dropped, resulting in 1,609 approved reads in version 1, and 1,558 approved reads in version 2.
- RPPA data were left untouched, expressed in terms of 454 antigen IDs (AGIDs).
- MS data were originally expressed as 14,569 Entrez IDs; these were updated by replacing obsolete IDs with the current ones, and mapped to the namespace of the NCBI Gene dataset; 1 unmatched Entrez ID was dropped. MS data contained 34.30% not-a-number (NaN) values, due to undetected proteins. As MS values are expressed in terms of log-ratio, those values would correspond to a readout of −∞; the smallest non-NaN value reported was –28.38; as a result, to cap those at a finite value, all NaN’s were replaced with the arbitrary value of –30.0; the effect of such capping is visible in Supplementary Figure 4. Finally, synonymous IDs were aggregated by mean; all-flat reads were dropped, resulting in 14,093 approved reads.

Next, sample metadata were generated, combining columns from the biospecimen manifest and the clinical manifests, from GDC and PDC. Notably, the tissue_or_organ_of_origin column was harmonised, to make sure it reflected the biology of the samples, by applying the transformation depicted in Supplementary Table 2. Purposefully, tissue groups were left relatively broad: this was done to guarantee that, later on, the Monte Carlo method would be able to detect at least two biological clusters within each group.

Finally, the RNAseq, miRNAseq and RPPA data from version 1 were aggregated into an object of the MultiOmicDataset class, from our VLab library. Likewise, the RNAseq, miRNAseq and MS data from version 2 were aggregated into another MultiOmicDataset object. These objects will henceforth be referred to as the RPPA Proteomizer dataset, and the MS Proteomizer dataset, respectively. Their final composition is depicted in Table 2.

### Pre-processing

We applied pre-processing to make data distribution more suitable for ML processing.

- For Tx and Mx, after mapping genes into the Entrez namespace, we re-applied the normalization by transcripts per million mapped reads (TPM), as recommended by Tx guidelines.^107,108^ Next, we log-transformed the data to ensure that all reads would occupy the same order of magnitude.

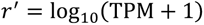

- RPPA Px is measured based on antibody binding, using a panel of antibodies specific for different proteins (454 antibodies in the case of TCGA). Reads are expressed in terms of an optical score in an arbitrary scale. Notably, values are not cross-comparable from a protein to another, as they are assessed using different antibodies: for instance, if a sample displays 1.2 for EGFR and 2.4 for BRCA1, it does not mean that EGFR is present in half the nanogram quantity of BRCA1; each protein is on a scale of its own. In order to make the Px reads interpretable across different proteins, we leveraged the large size of our dataset: we converted the reads into Z-scores, calculated gene-centrically across all samples. As a result, each RPPA value expresses how many standard deviations it is off, from the mean value across all the tissues. This way, results are not only uniformly scaled, but they also become human-interpretable. we maxed the Z-scores at ±5 (or at ±6), to avoid excessive weight on outliers.

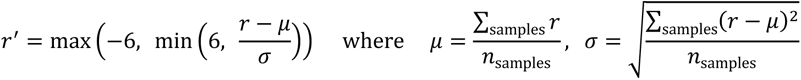

- Similar considerations apply for mass spectrometry Px. Because CPTAC utilized TMT, peaks for each sample are expressed as a log-ratio against the peak produced by a common reference sample called “POOL”. The latter is a mixture of a high number of cancer biopsies from a diverse set of organs; as a result, it contains non-zero amounts of virtually all human proteins.^109^ It follows that, like in the RPPA case, each protein is expressed in a scale that is completely independent of that of other proteins. To account for this problem, we applied the same pre-processing procedure as for RPPA. Additionally, undetected proteins (which would correspond to a log-ratio of −∞) were capped at a log-ratio of –50.

Distribution density plots confirmed that the pre-processing transformations made data remarkably more manageable, and closer to a normal distribution, in both the RPPA version (Supplementary Figure 3) and the mass spectrometry version (Supplementary Figure 4) of the dataset.

### Multi-layer perceptron

For both versions of the Proteomizer dataset, a multi-layer perceptron (MLP) model was trained on a multi-variable regression task, to predict the Px landscape, provided the transcriptomic Tx and Mx ones. Samples (examples) were split into 10 cross-validation (CV) folds, each composed of 80% training examples, 10% validation examples and 10% test examples. Splitting was conducted in a stratified way, with respect to the tissue of origin. The MLP was instantiated from the PyTorch Sequential class; it was trained using the AdamW optimiser, and the MSELoss (mean squared error, MSE) loss. To prevent overfitting, early stopping was employed, based on the MSE in the validation dataset. At each prediction, grid search was automatically conducted by VLab to select the best-performing hyperparameters in the validation set. At the end of the grid search, a final “output” training epoch was performed in a 2-set split, namely 90% training examples and 10% validation examples, using the optimal hyperparameter combination: the output epoch is meant as the best prediction for the model, and can be used for all purposes other than measuring ML metric.

For each grid search epoch, three predicted proteomes (Px’) are generated: (1) a test dataset version, which is a concatenation of the model’s output corresponding to the 10% test dataset examples, repeated across the 10 CV folds so as to cover the entirety of the dataset. (2) A validation dataset version, which is created in the same way (except in the “output” epoch, where validation is not available). (3) A training dataset version, which is created by averaging the model’s output from the training datasets across the 10 CV folds; averaging is required, as the training dataset occupies 80% of the sample space in each fold (or 90% in output mode); as a result, each sample will yield the average across 8 predicted reads (9 in output mode).

### Gene set enrichment analysis

At the end of the MLP training, output predictions underwent gene set enrichment analysis (GSEA). The analysis was carried out separately for each of these three vectors: (1) the Spearman correlation coefficient *ρ* between Tx and Px; (2) the *ρ* between the predicted Px’ and the Px; (3) the proteomization benefit, measured as the arithmetic difference between (2) and (1). The target classes were sourced from Gene Ontology,^49^ as previously described. GSEA computation was conducted using the GSEApy Python library, retrieving four values for each GO term: an enrichment score, a normalized enrichment score, a p-value and false discovery rate (FDR). The top 25 and bottom 25 GO terms with respect to FDR were plotted on a t-distributed stochastic neighbor embedding (t-SNE) plot, conducted on the membership matrix of the GO dataset (Figure 2).

### Monte Carlo method for differential expression analysis simulation

The analysis takes in as inputs the MLP result from the best grid search epoch, consisting of three versions of the predicted Px’, namely a training dataset prediction, a test dataset prediction, and a training dataset prediction; and the original multiomic dataset, consisting of Tx, Mx and Px. Samples within each tissue of origin were clustered by *k*-means clustering on the Px portion, representing the “ground truth” of our prediction; clustering was repeated for a range of *k* values from 2 to 10, and the best epoch was selected by the silhouette score. Clusters with less than 20 samples were dropped.

Next, the following subroutine was repeatedly executed. Within the tissue of origin *t* (say, liver), a number *n* of samples were randomly selected from one of the clusters, representing the “experimental group” of a hypothetical differential expression analysis experiment; the same number *n* of samples were selected from a different cluster, representing its matched “control group”. Wald test was applied to all genes in the dataset: for every gene, the Px’ reads 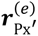 in the experimental group (a vector of length equal to *n*) and the Px’ reads s 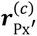 in the control group (another vector of length equal to *n*) were compared by Wald test; each Wald test returned a *W*_Px_′ statistic and a p-value *p*_Px_′ for every gene, expressing how differentially expressed the gene is in the experimental group samples, compared to the control group samples. The same procedure was carried out for Tx, and for Px. Next, two metrics were computed to quantify the Px’-Px agreement, and the Tx-Px agreement.

- Δ log *p*: a continuous metric, which quantifies the difference between the signed, log-transformed p-values. A “signed” p-value indicates a “+” sign if the gene is upregulated in the experimental group, a “–” sign if the gene is downregulated. The rationale is that, ideally, p-values in the Px’ should be similar to those in the ground truth Px, whereas the p-values in Tx should be relatively distant from those in Px. By averaging out the Δ*p*_Px_′_, Px_ across all genes in the dataset, we are effectively measuring the distance between the significance scores in the predicted Px’ and those in the ground truth Px; similarly, by averaging out the Δ*p*_Tx, Px_ across all genes in the dataset, we are effectively measuring the distance between the significance scores in the raw Tx and those in the ground truth Px. Proteomization is beneficial if the former number is smaller than the latter.

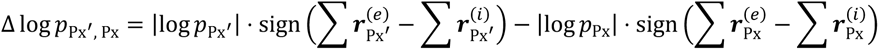

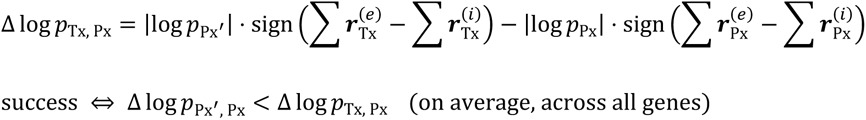

- Label agreement (LA): a discrete metric, which assesses whether the thresholded p-values agree between Px’ and Px, or between Tx and Px; the thresholded p-value is expressed in the form of a label *l*, which is assigned as follows: “+1” if a gene is significantly (*p* < 0.05) upregulated in the experimental group; “–1” if it is significantly (*p* < 0.05) downregulated; “0” if the comparison is not significant. The label agreement LA between two domains (e.g., between Px’ and Px) is defined as equal to 1.0 if the two labels are identical; 0.0 if that is not the case. By averaging out the LA_Px_′_, Px_ across all genes in the dataset, we are effectively measuring how often the predicted Px’ correctly “guesses” the differentially expressed genes from the Px; similarly, by averaging out the LA_Tx, Px_ across all genes, we are effectively measuring how often the raw Tx correctly “guesses” the differentially expressed genes from the ground truth Px; proteomization is beneficial if the former number is greater than the latter.

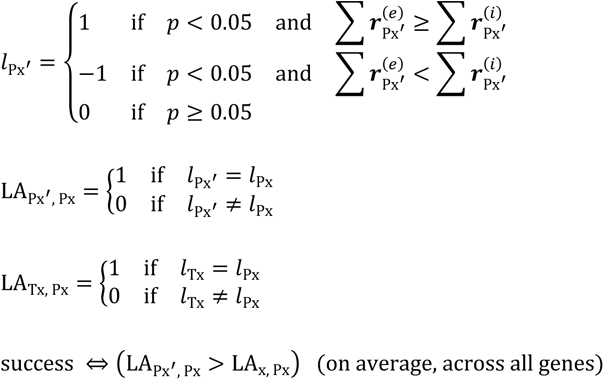

The subroutine was repeated for a range of sample sizes *n*, namely 3, 4, 5, 6, 8, 10, 12, 15, 20, 30, 50; combinatorially, for a range of statistical tests (Student’s t-test, Mann-Whitney/Wilcoxon rank-sum u-test, Wald w-test); and combinatorially, for all tissues of origin available in the dataset (liver, kidney, pancreas, etc.). For every iteration of the subroutine:

- the average Δ log *p*_Px_′_, Px_ was plotted against the average Δ log *p*_Tx, Px_ : the idea being that, if proteomization was beneficial, then the former should smaller than the latter;
- similarly, the average LA_Px_′_, Px_ was plotted against the average LA_Tx, Px_ : the idea being that, if proteomization was beneficial, then the former should be larger than the latter.

### Leave-one-group-out training

The MLP regressor was launched, as previously described. However, instead of employing a stratified CV where all tissues are equally represented across all CV folds, a leave-one-group-out (LOGO) cross-validation was

employed: in each fold, all samples were present except for those belonging to one specific tissue of origin. The training was repeated while rotating the held-out tissue across all tissues in the dataset: it follows that the number of LOGO folds is exactly identical to the number of tissues in the dataset.

### ROSMAP and MSBB datasets

The four Alzheimer’s disease (AD) multiomic datasets were sourced from the Religious Orders Study/Memory and Aging Project (ROSMAP) study (batch 1 and 2)^54,55^; the Mount Sinai Brain Bank (MSBB)^44,56^ study; and the ROSMAP iPSC-Derived Induced Neurons (ROSMAP-IN).^57,58^ Specifically, they were accessed through the AD Knowledge Portal,^53^ after submitting a request for access through the Synapse.org portal,^110^ which was approved (ID: 9603055). Files were filtered by the “Study” field, selecting the following values: “MSBB”, “ROSMAP” and “ROSMAP-IN”. Next, they were manually download from the portal. RNAseq, miRNAseq and MS were processed analogously to TCGA and CPTAC, with minor variations due to the specific format of each study. Four versions of the dataset were created, one for each of the afore-mentioned studies. Data were aggregated into MultiOmicDataset objects, counting 38,897 reads by 209 samples in ROSMAP batch 1; 36,130 reads by 110 samples in ROSMAP batch 2; 27,722 reads by 184 samples in MSBB; 20,384 reads by 36 samples in ROSMAP RI. The example results showcased in Supplementary Figure 8 were generated using the ROSMAP batch 1; no significant differences were observed using the other three datasets.

In all analyses involving both the TCGA-CPTAC MS dataset and one of the ROSMAP/MSBB datasets, only the reads in common between both datasets were utilized; the remainder reads were dropped.

### Proteomizer explainers

Three architectures were employed to predict which genes (either in Tx, Px or Mx) are the most “helpful” (hence the name, “helper” genes) in explaining the Tx-Px mismatch for a given target gene of interest (GOI). The analysis was repeated across 100 GOIs, selected for having the highest number of references in the PubMed repository. Each explainer was run on the following six tasks:

- Three “direct” tasks:
  1. From the GOI’s Tx, across all helper genes in the Tx space (other than the GOI itself), predict the GOI’s Px.
  2. From the GOI’s Tx, across all helper proteins in the Px space (other than the GOI itself), predict the GOI’s Px.
  3. From the GOI’s Tx, across all helper miRNAs in the Mx space (other than the GOI itself), predict the GOI’s Px.
- Three “reverse” tasks:
  4. From the GOI’s Px, across all helper genes in the Tx space (other than the GOI itself), predict the GOI’s Tx.
  5. From the GOI’s Px, across all helper proteins in the Px space, predict the GOI’s Tx.
  6. From the GOI’s Px, across all helper miRNAs in the Mx space, predict the GOI’s Tx.

The three explainer architectures were: Extreme Gradient Boosting (XGBoost)^60^ importance scores, Shapley Additive exPlanations (SHAP),^61^ and a novel approach that we named “Information Gain”. XGBoost is primarily a ML architecture, compatible with both classification and regression tasks, that is based on an ensemble of decision trees; however, it differs from random forests as the trees are trained subsequently, whereby each tree corrects the errors made by the preceding ones. It is popular in the ML community for its remarkable robustness and flexibility, as it often obtains top-performing performance across a diverse variety of ML problems. Additionally, it comes with in-built explainability, as the model output includes an unsigned importance score for each of the features provided in training.^60^ Conversely, SHAP is purely an explainer; it is a game theory-based method that can be applied *post-hoc* to a trained ML model, and returns signed importance scores for each of the provided features. It works by measuring the average marginal contribution of a feature to the model’s prediction, yielding a signed importance score.^61^ Finally, Information Gain is a novel explainer architecture that works by training an independent ML model for every feature in the dataset, alongside the “origin” omic vector, while targeting the “destination” omic vector; it then utilizes the models’ performance as a proxy for the importance of its associated feature.

The next paragraphs will describe the exact ML task on which the three explainer channels were trained.

In the XGBoost importance and SHAP channels, one XGBoost model was trained for every (task, GOI) tuple, resulting in 600 models being trained. Each model was fed with a training dataset that comprised the origin omic of the GOI (i.e., a column vector), concatenated with the whole helper omic full matrix; the target vector comprised the destination omic of the GOI.

More formally, be **Dataset** the TCGA-CPTAC dataset; be *n* the number of samples in the dataset (equal to 1,594) and be *m*_Tx_, *m*_Mx_ and *m*_Px_ the number of columns, corresponding to the Tx, Mx and Px readouts in each sample, respectively:

**Tx** ∈ ℝ^*n*×*m*Tx^ where *m*_Tx_ = 35,833

**Mx** ∈ ℝ^*n*×*m*Mx^ where *m*_Mx_ = 1,558

**Px** ∈ ℝ^*n*×*m*Px^ where *m*_Px_ = 14,094

**Dataset** = **Tx** || **Mx** || **Px** ∈ ℝ^*n*×*m*^ where *m* = *m*_Tx_ + *m*_Mx_ + *m*_Px_ = 51,485

Be 𝒢 our set of 100 GOIs; be matrix 𝑿 the training dataset, and vector 𝒀 the target vector of a ML model.

Then, for each GOI, 6 XGBoost models were trained on the following 𝑿 and 𝒀:

∀𝑔 ∈ 𝒢:

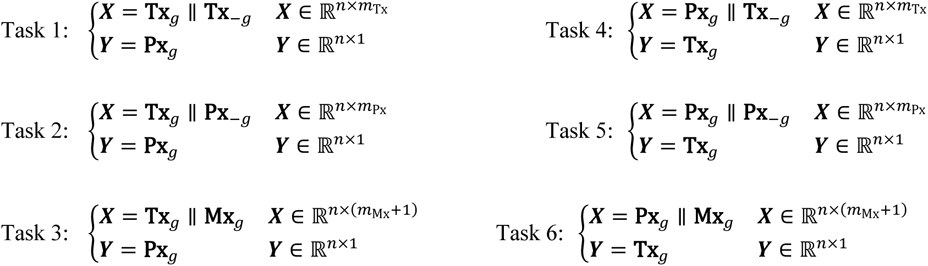

All training epochs were conducted in a 10-fold cross-validation setting, using early stop in the validation dataset to prevent overfitting; hyperparameters were automatically optimized by VLab by means of a grid search. At the end of each training, XGBoost feature importance scores were collected, alongside SHAP values importance scores for the test dataset prediction, the validation dataset prediction, and the training dataset prediction; the test dataset version of the importance scores was used for subsequent comparison with the Biology Mega Graph.

Conversely, in the Information Gain channel, one XGBoost model was trained for every (task, GOI, helper gene) tuple, resulting in 5.148.300 models being trained. Each model was fed with a training dataset that comprised the origin omic of the GOI (i.e., a column vector), concatenated with helper omic of a single helper gene (i.e., another column vector); the target vector comprised the destination omic of the GOI.

More formally:

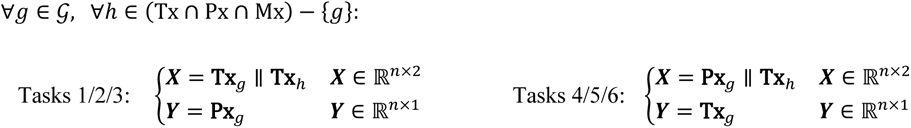

All training epochs were conducted in a 10-fold cross-validation setting, using early stop in the validation dataset to prevent overfitting; hyperparameters were automatically optimized by VLab by means of a grid search. At the end of each training, the effectiveness of the provided helper was quantified in terms of the mean absolute error (MAE); the test dataset version of the performance scores was used for subsequent comparison with the Biology Mega Graph (0).

Overall, all three explainers answer the same question: which gene’s Tx, gene’s Px or miRNA’s Mx are most informative for the GOI’s Tx-Px mismatch level?

### Biology Mega Graph

11 relational datasets, encompassing gene-gene or miRNA-gene relations, were downloaded from their respective sources and parsed, namely: ComplexPortal,^63^ human DEPhOsphorylation Database (DEPOD),^64^ GO,^65^ microT (MirBase and MirGeneDB versions),^66^ Mirbase,^67^ NCBI Gene,^68^ PhosphoSitePlus,^69^ Signor,^70^ STRING^71^ and TarBase.^72^ GO was parsed as previously described. Relation labels underwent manual standardization and curation, to make them cross-compatible across the different data sources. All genes were converted from their original namespaces into the NCBI Entrez namespace, and miRNA names into the miRBase namespace^67^; unmatched names were dropped; synonymous entries were aggregated by a logical *OR*. The edge labels were harmonized into a common, fixed vocabulary. The final result was a knowledge graph encompassing 69 levels (one for every biological interaction modality) and 322,255,334 known regulatory relations, among 193,384 gene Entrez IDs and 3,537 miRNAs; its exact composition is depicted in Supplementary Table 1.

### Hardware

All development was conducted on a high-performance mainframe computer, featuring Nvidia 4090 GPU and Intel i9-14900K 3.20 GHz CPU.

### Data availability

The VLab library^62^ source code, including the full GhostBuster module, can be found in the following repository: https://github.com/giuliodeangeli/vlab

## Acknowledgements

The authors would like to express their sincere gratitude to all those who provided invaluable advice, insightful discussions, and constructive feedback throughout this project. Their expertise and guidance greatly contributed to the development of this work. In particular, we acknowledge the contributions of Cristiano Peron, Antonio Cisternino, Pietro Barbiero, Niccolò Pancino, Michelangelo Diligenti, Lorenzo Giusti, Donato Crisostomi, Giuseppe Marra, Gabriele Ciravegna, Francesco Giannini, Aviva Tolkovski and Giorgio Vivacqua.

This research was supported by funding from the Cambridge Trust and the Medical Research Council Doctoral Training Partnership (MRC DTP). We acknowledge their generous support in facilitating this study.

## Author contributions

G.D. was responsible for dataset preparation, machine learning model development and training, explainability analysis, literature review, manuscript writing.

M.G.S. and P.L. provided conceptual guidance and contributed to manuscript revisions.

## Statements

### TCGA/CPTAC studies

The results shown here are in whole or part based upon data generated by the TCGA Research Network: https://www.cancer.gov/tcga.

Data used in this publication were generated in part by the Clinical Proteomic Tumor Analysis Consortium (CPTAC).

### ROSMAP study

The results published here are in whole or in part based on data obtained from the AD Knowledge Portal (https://adknowledgeportal.org).

Study data were provided by the Rush Alzheimer’s Disease Center, Rush University Medical Center, Chicago. Data collection was supported through funding by NIA grants P30AG10161 (ROS), R01AG15819 (ROSMAP; genomics and RNAseq), R01AG17917 (MAP), R01AG30146, R01AG36042 (5hC methylation, ATACseq), RC2AG036547 (H3K9Ac), R01AG36836 (RNAseq), R01AG48015 (monocyte RNAseq)

RF1AG57473 (single nucleus RNAseq), U01AG32984 (genomic and whole exome sequencing), U01AG46152 (ROSMAP AMP-AD, targeted proteomics), U01AG46161(TMT proteomics), U01AG61356 (whole genome sequencing, targeted proteomics, ROSMAP AMP-AD), the Illinois Department of Public Health (ROSMAP), and the Translational Genomics Research Institute (genomic).

TMT Proteomics: Study data were provided through the Accelerating Medicine Partnership for AD (U01AG046161 and U01AG061357) based on samples provided by the Rush Alzheimer’s Disease Center, Rush University Medical Center, Chicago. Data collection was supported through funding by NIA grants P30AG10161, R01AG15819, R01AG17917, R01AG30146, R01AG36836, U01AG32984, U01AG46152, the Illinois Department of Public Health, and the Translational Genomics Research Institute.

RNA-seq Bulk Brain: “Annie J. Lee, Yiyi Ma, Lei Yu, Robert J. Dawe, Cristin McCabe, Konstantinos Arfanakis, Richard Mayeux, David A. Bennett, Hans-Ulrich Klein, and Philip L. De Jager. Multi-region brain transcriptomes uncover two subtypes of aging individuals with differences in Alzheimer’s disease risk and the impact of APOEε4. bioRxiv 2021”

#### MSBB study

The results published here are in whole or in part based on data obtained from the AD Knowledge Portal (https://adknowledgeportal.org/). These data were generated from postmortem brain tissue collected through the Mount Sinai VA Medical Center Brain Bank and were provided by Dr. Eric Schadt from Mount Sinai School of Medicine.

TMT Proteomics: The results published here are in whole or in part based on data obtained from the AD Knowledge Portal (https://adknowledgeportal.org/). These data were provided by Dr. Levey from Emory University based on postmortem brain tissue collected through the Mount Sinai VA Medical Center Brain Bank provided by Dr. Eric Schadt from Mount Sinai School of Medicine.

#### ROSMAP-IN

The results published here are in whole or in part based on data obtained from the AD Knowledge Portal (https://adknowledgeportal.org). Study data were provided by the Young-Pearse Lab of the Ann Romney Center for Neurologic Disease at Brigham and Women’s Hospital (BWH), Boston. Stem cell lines used in this study were generated through collaboration between BWH, the New York Stem Cell Foundation, and the Rush Alzheimer’s Disease Center, Rush University Medical Center, Chicago.

## Supplementary Figures

**Supplementary Figure 1.**
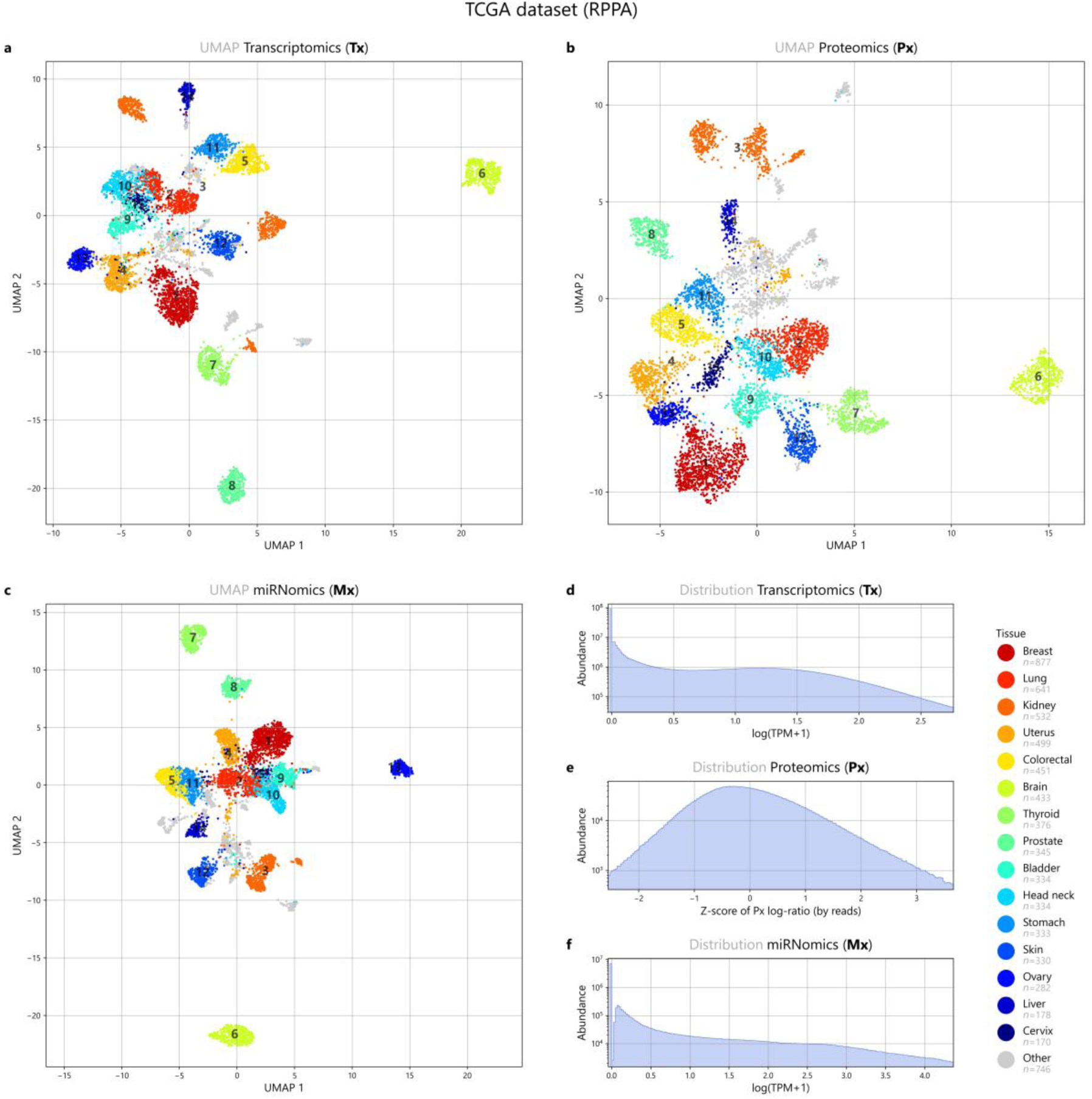
Dimensionality reduction plot in the RPPA version of the TCGA dataset. This plot represents a dimensionality-reduced overview of the dataset that was used to train Proteomizer—specifically, the version where protein measurements were conducted using reverse-phase protein array (RPPA). (a-c) Uniform manifold approximation and projection (UMAP) was applied to the 7.019 samples, which represent human patient biopsies from healthy or cancerous tissues, from 30+ tissues of origin (see Table 2); it was repeated for each of the three omics of interest: transcriptomics (Tx, 35,979 reads), miRNomics (Mx, 1,609 reads) or RPPA proteomics (Px, 454 reads), aggregated by their Entrez ID or miRBase standard name. Color coding conveys the 15 of the 30+ tissues of origin of the samples (for metastatic cancer, this corresponds to the primary histological origin of the tumor, e.g., colorectal metastasis in the liver would qualify as colorectal). The UMAP plot highlights how in all three omics, taken individually, samples make up clearly demarked clusters, which generally correspond to one specific tissue of origin. This suggests that the dataset was assembled correctly, and real biological signal is present in the data. (d-f) Distribution profile, in log-transformed coordinates, for each of the three omics of interest. UMAP parameters: n_neighbors=80, min_dist=0.51.

**Supplementary Figure 2.**
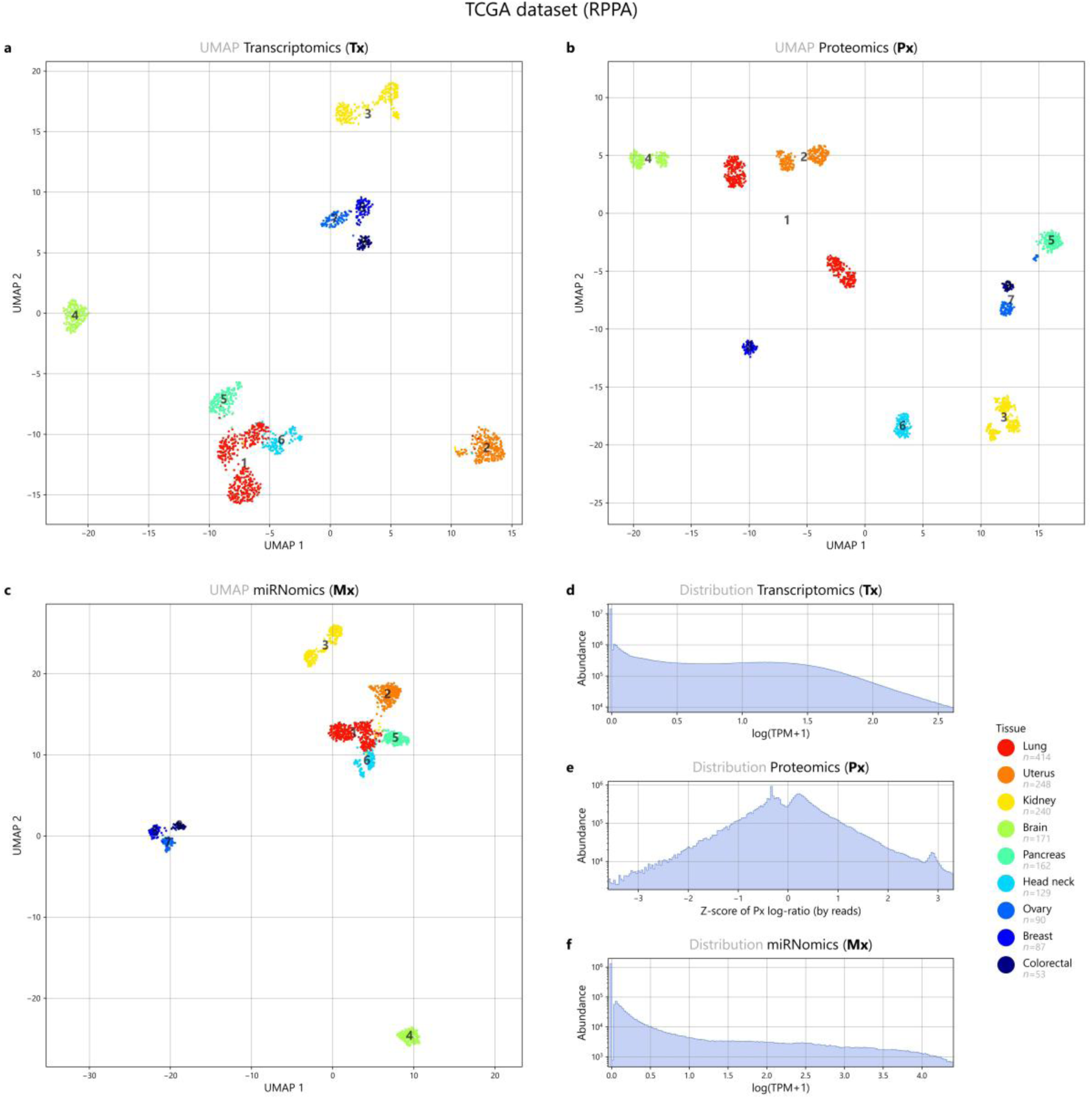
Dimensionality reduction plot in the mass spectrometry version of the TCGA- CPTAC dataset. This plot represents a dimensionality-reduced overview of the dataset that was used to train Proteomizer—specifically, the version where protein measurements were conducted using mass spectrometry tandem mass tag (TMT). (a-c) Uniform manifold approximation and projection (UMAP) was applied to the 1,594 samples, which represent human patient biopsies from healthy or cancerous tissues, from 30+ tissues of origin (see Table 2); it was repeated for each of the three omics of interest: transcriptomics (Tx, 35,833 reads), miRNomics (Mx, 1,558 reads) or TMT proteomics (Px, 14,094 reads), aggregated by their Entrez ID or miRBase standard name. Color coding conveys the 9 tissues of origin of the samples (for metastatic cancer, this corresponds to the primary histological origin of the tumor, e.g., colorectal metastasis in the liver would qualify as colorectal). The UMAP plot highlights how in all three omics, taken individually, samples make up clearly demarked clusters, which generally correspond to one specific tissue of origin. This suggests that the dataset was assembled correctly, and real biological signal is present in the data. (d-f) Distribution profile, in log-transformed coordinates, for each of the three omics of interest. UMAP parameters: n_neighbors=55, min_dist=0.50.

**Supplementary Figure 3.**
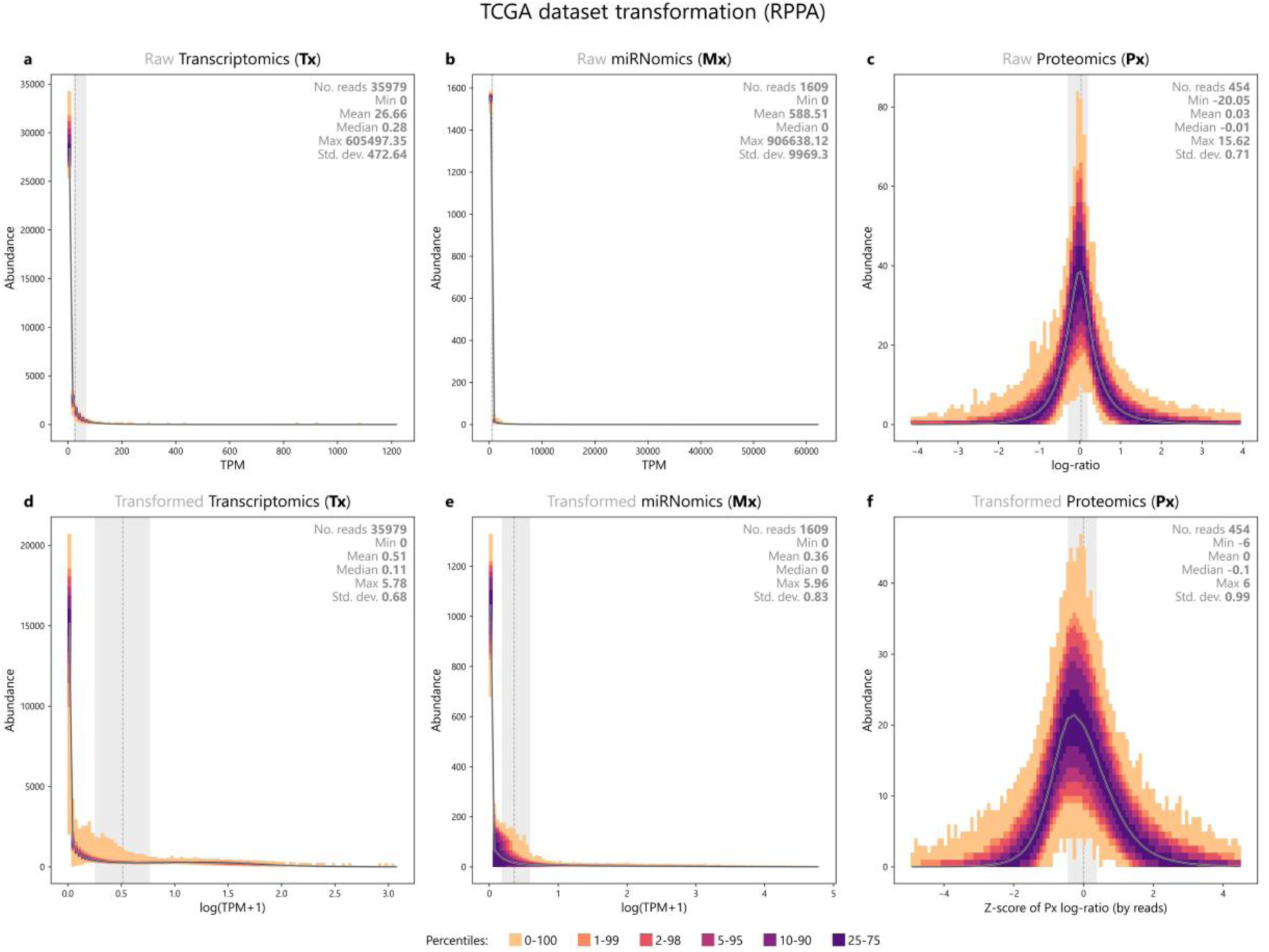
Density plot showcasing the effects of the pre-processing step, on the RPPA version of the TCGA dataset. Density plot showcasing the effects of the pre-processing step, on the RPPA version of the TCGA dataset This plot represents the pre-processing step of the dataset used to train Proteomizer—specifically, the version where protein measurements were conducted using reverse-phase protein array (RPPA). It showcases a before-after comparison of the effects of pre-processing, in terms of data distribution within each of the three omics of interest. Raw data were sourced from The Cancer Genome Atlas (TCGA). A dataset was assembled, made up of 7.019 samples, and 38,042 features. Samples represent human patient biopsies from healthy or cancerous tissues, from 30+ tissues of origin (see Table 2). Reads represent multiomic measurements by either transcriptomics (Tx, 35,979 reads), miRNomics (Mx, 1,609 reads) or RPPA proteomics (Px, 454 reads), aggregated by their Entrez ID or miRBase standard name. The top row depicts the original TCGA data, where Px is expressed in terms of RPPA optic score, while Tx and Mx data are expressed as transcripts per million mapped reads (TPM). The bottom row depicts the pre-processed versions thereof, where Px data underwent Z-scoring capped at ±5, while Tx and Mx data underwent log-transformation (after adding 1, to avoid log of 0). Colors represent a density plot, where lighter tones embrace the entire dataset, while darker tones depict data subsets progressively closer to the mode, for each value interval. The overall median is represented as a dark grey dashed line, and the ±σ confidence interval is show as a light grey background. RPPA raw data were already remarkably close to a normal distribution, consistently with the fact that they are an optic measurement from an antibody-based assay; as a result, Z-transformation had little effect on their overall landscape. Tx and Mx originally followed a descending monotonic Zipf’s distribution, aligning with their typical pattern; while the majority of genes have 0 or near-0 expression, the log transformation effectively reduced the order-of-magnitude span occupied by the distribution.

**Supplementary Figure 4.**
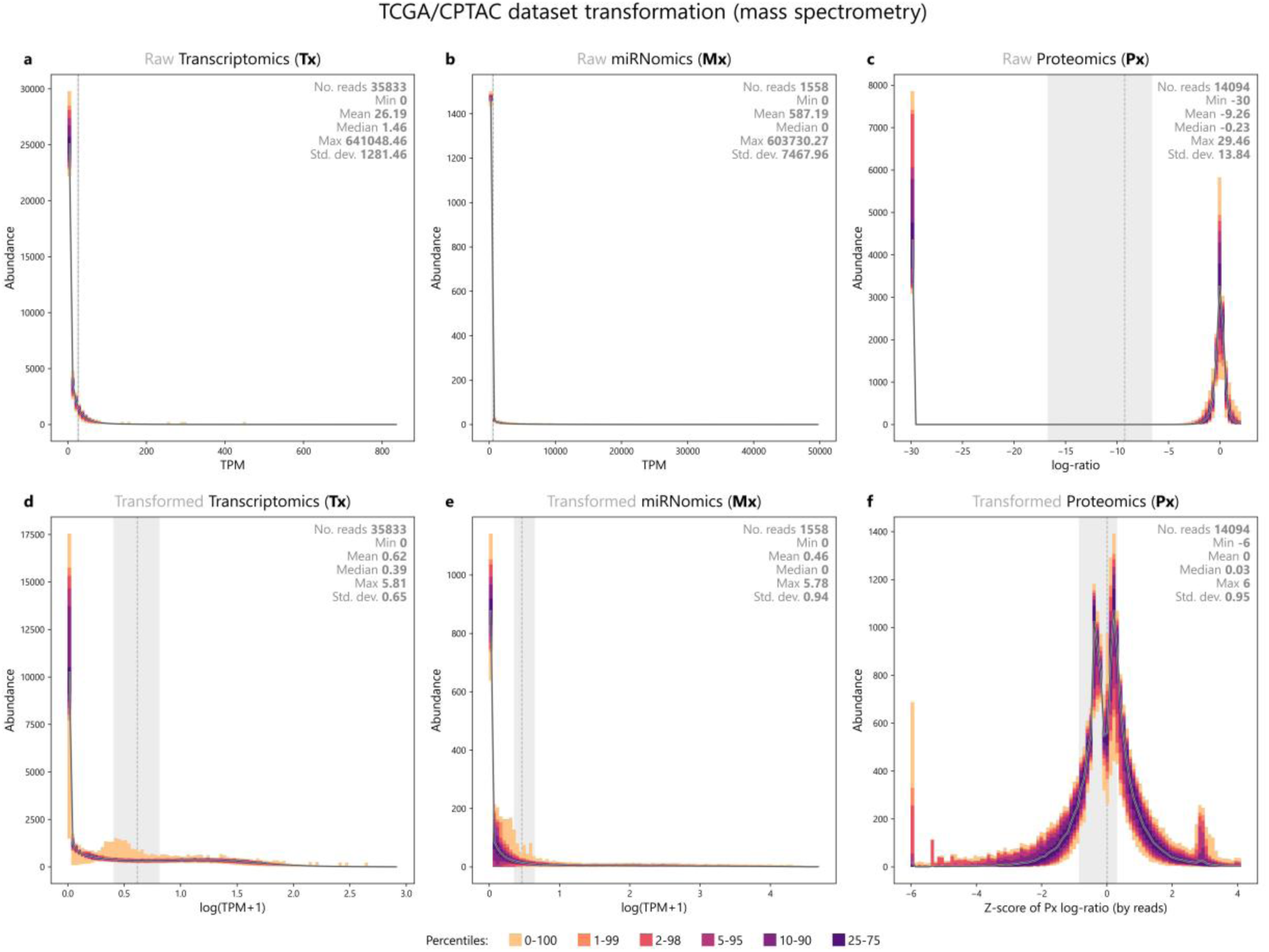
Density plot showcasing the effects of the pre-processing step, on the mass spectrometry version of the TCGA-CPTAC dataset. This plot represents the pre-processing step of the dataset used to train Proteomizer—specifically, the version where protein measurements were conducted using mass spectrometry tandem mass tag (TMT). It showcases a before-after comparison of the effects of pre-processing, in terms of data distribution within each of the three omics of interest. Raw data were sourced from The Cancer Genome Atlas (TCGA) and Clinical Proteomic Tumor Analysis Consortium (CPTAC). A dataset was assembled, made up of 1.594 samples, and 51,485 features. Samples represent human patient biopsies from healthy or cancerous tissue, from 9 tissues of origin (see Table 2). Reads represent multiomic measurements by either transcriptomics (Tx, 35,833 reads), miRNomics (Mx, 1,558 reads) or RPPA proteomics (Px, 14,094 reads), aggregated by their Entrez ID or miRBase standard name. The top row depicts the original TCGA data, where Px is expressed in terms of TMT log-ratio over the shared reference sample (POOL), while Tx and Mx data are expressed as transcripts per million mapped reads (TPM). The bottom row depicts the pre- processed versions thereof, where Px data underwent Z-scoring capped at ±5, while Tx and Mx data underwent log transformation (after adding 1, to avoid log of 0). Colors represent a density plot, where lighter tones embrace the entire dataset, while darker tones depict data subsets progressively closer to the mode, for each value interval. The overall median is represented as a dark grey dashed line, and the ±σ confidence interval is show as a light grey background. The log-ratio nature of Px expression measurements determined a noteworthy problem. Positive numbers represent a protein concentration that is higher than POOL; zero represents a concentration that is equivalent to POOL; and negative numbers represent a concentration lower than POOL; then, how to encode for the case of zero-expression (undetected protein), considering that log 0^+^ diverges to −∞? In the original CPTAC data, this case was represented as a not-a-number (NaN), which would be incompatible with subsequent machine learning. To solve this hurdle, we applied an arbitrary limit at –30.0 to the log-ratio distribution; such value was selected based on the fact that the minimum non-NaN value was –28.38. Since “zero” is, in general, the most frequently occurring expression level, the peak at –30.0 is clearly visible in panel (c); after applying Z-transformation, the peak that used to be at –30.0 appears to move remarkably close to 0.0: this is because the proteins with no expression in a given sample are often scarcely expressed across the majority of tissues, which puts both their mean and standard deviation levels at near-zero values. Overall, the before-after comparison of the Px landscape confirms that Z-scoring is remarkably effective at rendering the data closer to a Gaussian distribution, thereby more manageable from a machine learning perspective. Tx and Mx originally followed a descending monotonic Zipf’s distribution, aligning with their typical pattern; while the majority of genes have 0 or near-0 expression, the log transformation effectively reduced the order-of-magnitude span occupied by the distribution.

**Supplementary Figure 5.**
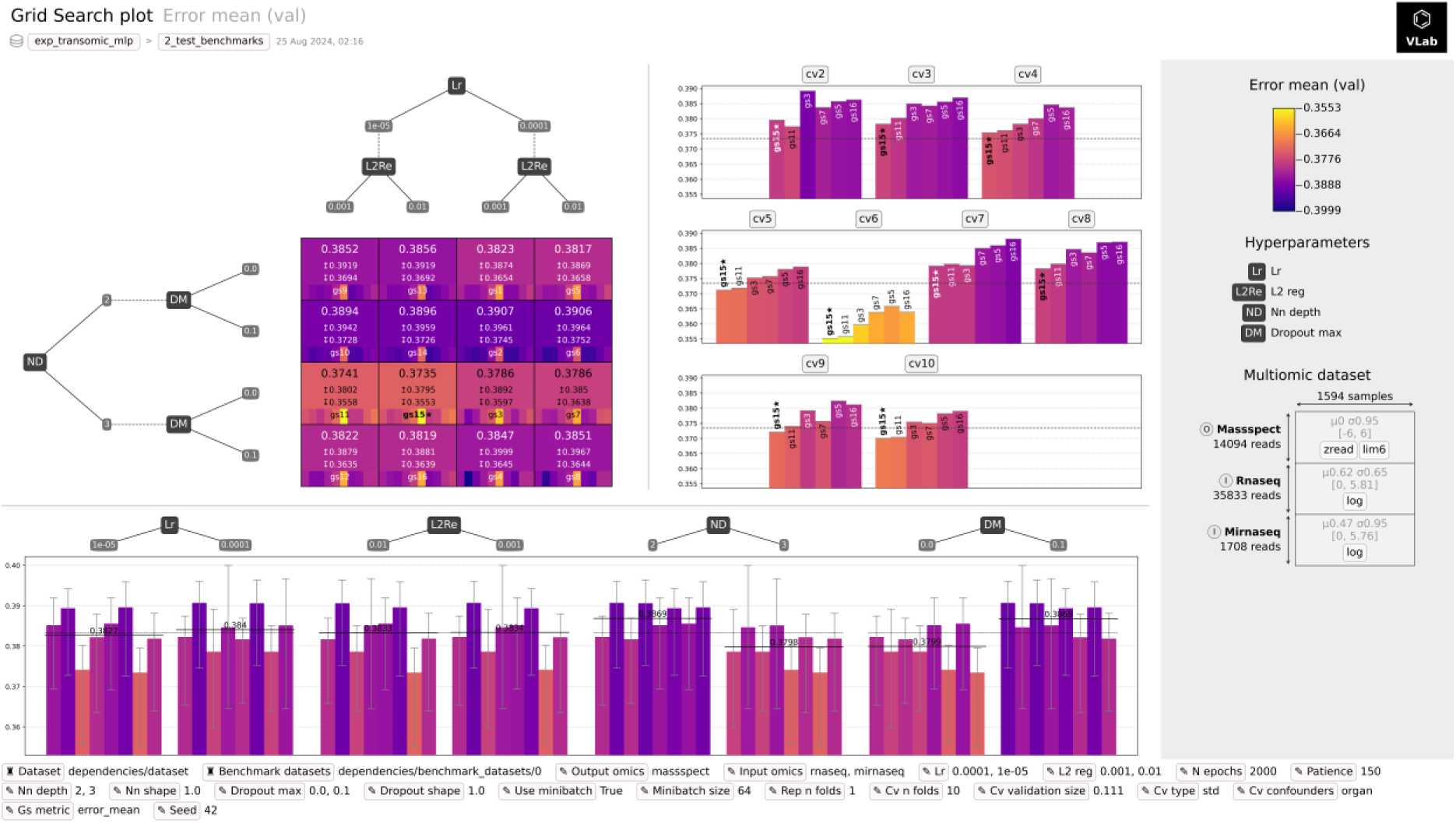
Grid search results for the proteomization task, on the mass spectrometry version of the TCGA-CPTAC dataset. This is an example plot from the Virtual Lab (VLab) machine learning framework. This plot represents a grid search for the best hyperparameter combination, in a multi-layer perceptron architecture tasked at predicting the proteomic landscape (Px) from the transcriptomic (Tx) and miRNomic (Mx) ones, in a 10-fold stratified cross-validation setting. Specifically, 16 hyperparameters combinations were searched, involving learning rate (Lr), L2 regularization (L2Re), network depth (ND) and maximum dropout share (DM). Performance is measured in terms of mean absolute error (MAE) in the validation dataset. The experiment was launched on the mass spectrometry version of the Proteomizer dataset, where Px values are expressed in the form of Z-scores, i.e., in terms of “number of standard deviations”. The best-performing grid search achieved a MAE of 0.375 standard deviations.

**Supplementary Figure 6.**
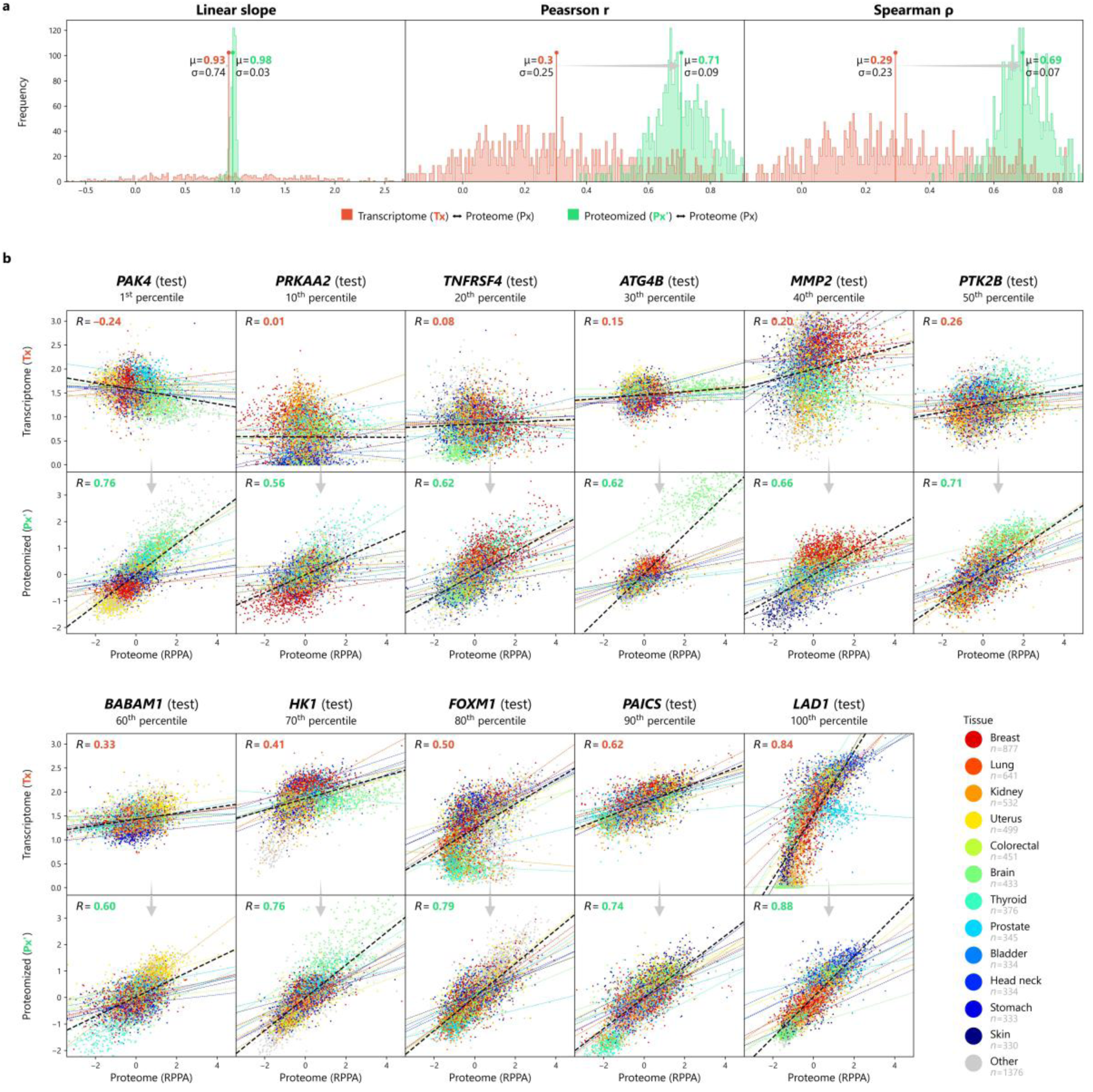
Correlation metrics before and after proteomization, in the RPPA version of the TCGA dataset. Proteomizer was applied to predict the reverse-phase protein array (RPPA) proteomic (Px) landscape, based on the transcriptomic (Tx) and miRNomic (Mx) landscape, on 7,019 samples sourced from The Cancer Genome Atlas (TCGA). After training, the predicted Px’ in the test dataset was compared with the original Tx and Px. (a-c) Three correlation metrics—namely the linear slope, the Pearson *r* correlation coefficient and the Spearman *ρ* correlation coefficient—were applied between raw Tx and Px (red), and between predicted Px’ and Px (green), for each of the 452 genes with both Tx and Px availability; the distribution profiles show the effect of proteomization, in a before-after fashion. (d) Scatter plots of the 7,019 samples for 11 exemplary genes, depicting the ground truth Px coordinates on the x-axis, and the raw Tx (top) or the predicted Px’ (bottom) on the y-axis. Genes were sampled in a diverse range of Tx-Px mismatch values, from the most mismatched (1^st^ percentile) to the most aligned (100^th^ percentile). Coordinates are identical from one plot to another. Colors indicate the tissue of origin (see Table 2). A linear regression line was computed in each plot, both for the entire gene set (black dashed line), and for individual organs (solid colored lines).

**Supplementary Figure 7.**
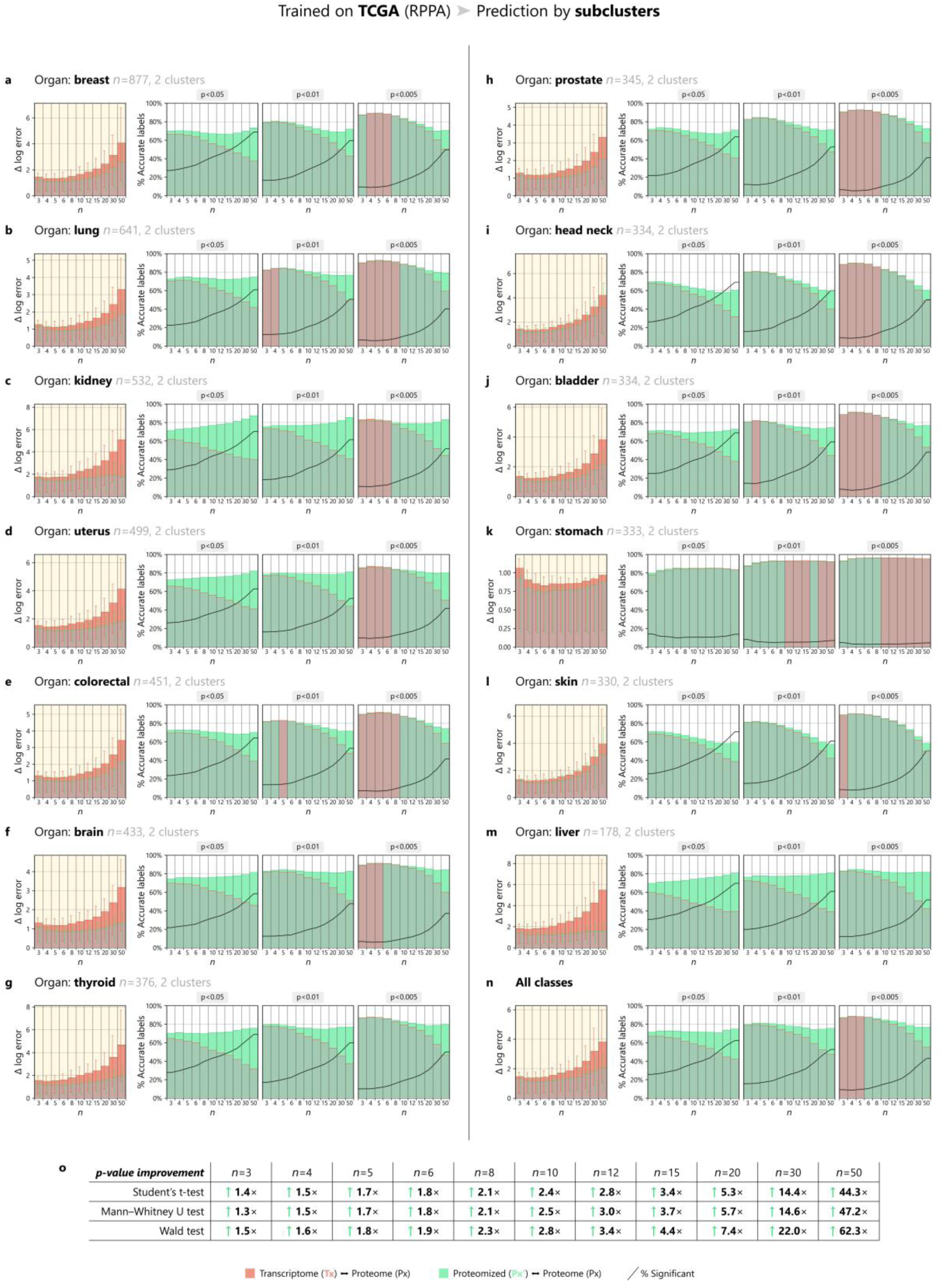
Monte Carlo simulation of differential expression analyses across sample clusters, in the RPPA version of the TCGA dataset. Proteomizer was applied to predict the reverse phase protein array (RPPA) proteomic (Px) landscape, based on the transcriptomic (Tx) and miRNomic (Mx) ones, on 7,019 samples sourced from The Cancer Genome Atlas (TCGA). At the end of the training, three full landscapes were made available: (i) the raw Tx, (ii) the predicted Px’ and (iii) the ground truth Px. Next, a Monte Carlo method was applied across sample clusters within each tissue of origin. The rationale was to determine whether proteomization had been beneficial, with respect to the customary endpoint of gene expression tests, that is differential expression analysis; in less formal words, the task was to assess whether Px’ would do a better job at predicting the differentially expressed genes (DEGs) in Px, compared to Tx. While a complete description of the Monte Carlo method is available in Materials and Methods, here follows a short summary. The analysis is repeated within the scope of each individual tissue of origin (a-m; note: 15 tissues are not shown here due to space constraints). First, samples are divided into 2-10 clusters by unsupervised machine learning (ML), specifically, by *k*- means clustering where *k* is optimized using the silhouette score. Next, the following iteration step is repeated 100 times, for varying sample size values *n* (x-axes in the figure). A number *n* of samples are randomly selected from one of the clusters, and the same number *n* of samples are randomly selected from a different cluster: they work as the experimental (exp) and control group (ctrl) respectively, of a hypothetical differential expression analysis experiment performed by a biologist. All genes undergo differential expression analysis by Wald test, comparing Txexp with Txctrl, Px’exp with Px’ctrl, and Pxexp with Pxctrl; these three comparisons yield a set of p-values for each gene. (Column 1) The p-values are log-transformed and assigned a sign, namely: “+” for the genes upregulated in the experimental group, “–” for the downregulated ones. The average difference between the log-transformed signed p-values of the Px’ with respect to Px constitutes the “Δ log error” of Px’, and is depicted in green. Likewise, the average difference between the log-transformed signed p-values of the Tx with respect to Px constitutes the “Δ log error” of Tx, and is depicted in red. If the proteomization is beneficial, then the green “Δ log error” in Px’ is supposed to be lower than in Tx—thus, leaving an emerging silhouette of red color standing out over the green. (Column 2) The p-values in Tx, Px’ and Px are flattened into labels, according to the following rule: label “+1” if a gene is significantly (*p* < 0.05) upregulated in the experimental group; label “–1” if a gene is significantly (*p* < 0.05) downregulated in the experimental group; label “0” if the comparison is not significant. Then, the number of genes where Px’ agrees with Px (in terms of labels) is counted, and goes to constitute the “% accurate labels” for Px’, which is depicted in green. Likewise, the number of genes where Tx agrees with Px is counted, and goes to constitute the “% accurate labels” for Tx, which is depicted in red. If the proteomization is beneficial, then the green “% accurate labels” in Px’ is supposed to be lower than in Tx—thus, leaving an emerging silhouette of green color standing out over the red. The black line depicts the % of significant comparisons; of note, p-values were not corrected for multiple testing (i.e., no Bonferroni or Benjamini-Hochberg corrections were applied), thereby it is expected that the percentage of significant comparisons will be strongly inflated. (Columns 3-4) Same as column 2, except the p-value threshold is fixed at 0.01 and 0.005, respectively (instead of 0.05); this makes the scenario of a statistically significant comparison rarer. (g) Aggregate measures across all tissues of origin. (h) Relative p-value improvement (average p-value for Px’ / average p-value for Tx) at different sample sizes, and using different statistics, namely: Student’s t-test, Mann-Whitney/Wilcoxon rank-sum u-test and Wald w-test. Overall, proteomization had a beneficial effect in the RPPA version of the TCGA dataset, yielding an average increase in Wald test “Δ log error” ranging from 1.5× (*n* = 3) up to 62.3× (*n* = 50). The benefit appears to be proportional to the sample size, being relatively marginal (or even slightly pejorative) with small samples due to low statistical power, and becoming very robust at larger samples, where the number of incorrectly identified labels is reduced by a factor of ∼50% (panel n, columns 2-4). The effect was beneficial across all tissues of origin (including the 15 not shown here due to space limitations). These results confirm that, as long as the tissues of interest are included in the training dataset, machine learning is an effective tool to alleviate the Tx-Px mismatch.

**Supplementary Figure 8.**
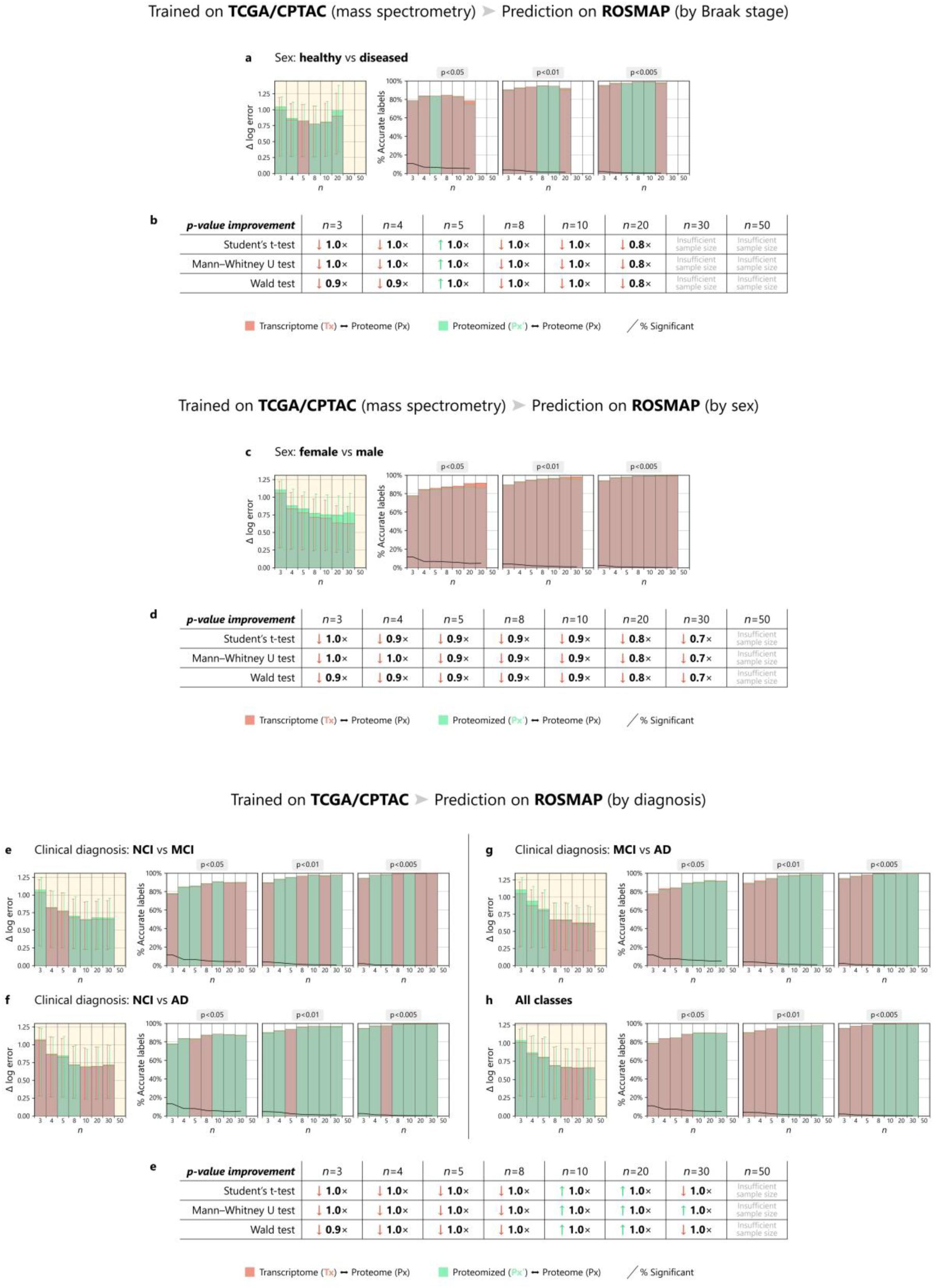
Monte Carlo simulation of differential expression analyses across sample clusters, trained on the mass spectrometry version of the TCGA-CPTAC dataset, predicting on the ROSMAP batch 1 dataset. Proteomizer was applied to predict the mass spectrometry proteomic (Px) landscape, based on the transcriptomic (Tx) ones, on 1,594 samples sourced from The Cancer Genome Atlas (TCGA) and Clinical Proteomic Tumor Analysis Consortium (CPTAC). At the end of the training, the model was used to infer the Px of 209 samples from the Religious Orders Study/Memory and Aging Project (ROSMAP) batch 1 study, based on the corresponding Tx samples. Only the 14,041 genes in common between the TCGA-CPTAC and ROSMAP datasets were utilized. As a result, three full landscapes were made available: (i) the raw Tx, (ii) the predicted Px’ and (iii) the ground truth Px (all three referred to ROSMAP). Next, a Monte Carlo method was applied across sample clusters within each tissue of origin. The Monte Carlo method was applied as described in Figure 3 (a full description can be found in Materials and Methods). Overall, proteomization had a net neutral or disruptive effect across all tested tissues, sample sizes and p-value thresholds. These results suggest that machine learning-based proteomization is not effective at translating to a completely independent dataset, where the Tx and Px were generated by two different research groups using different protocols and equipment.

**Supplementary Figure 9.**
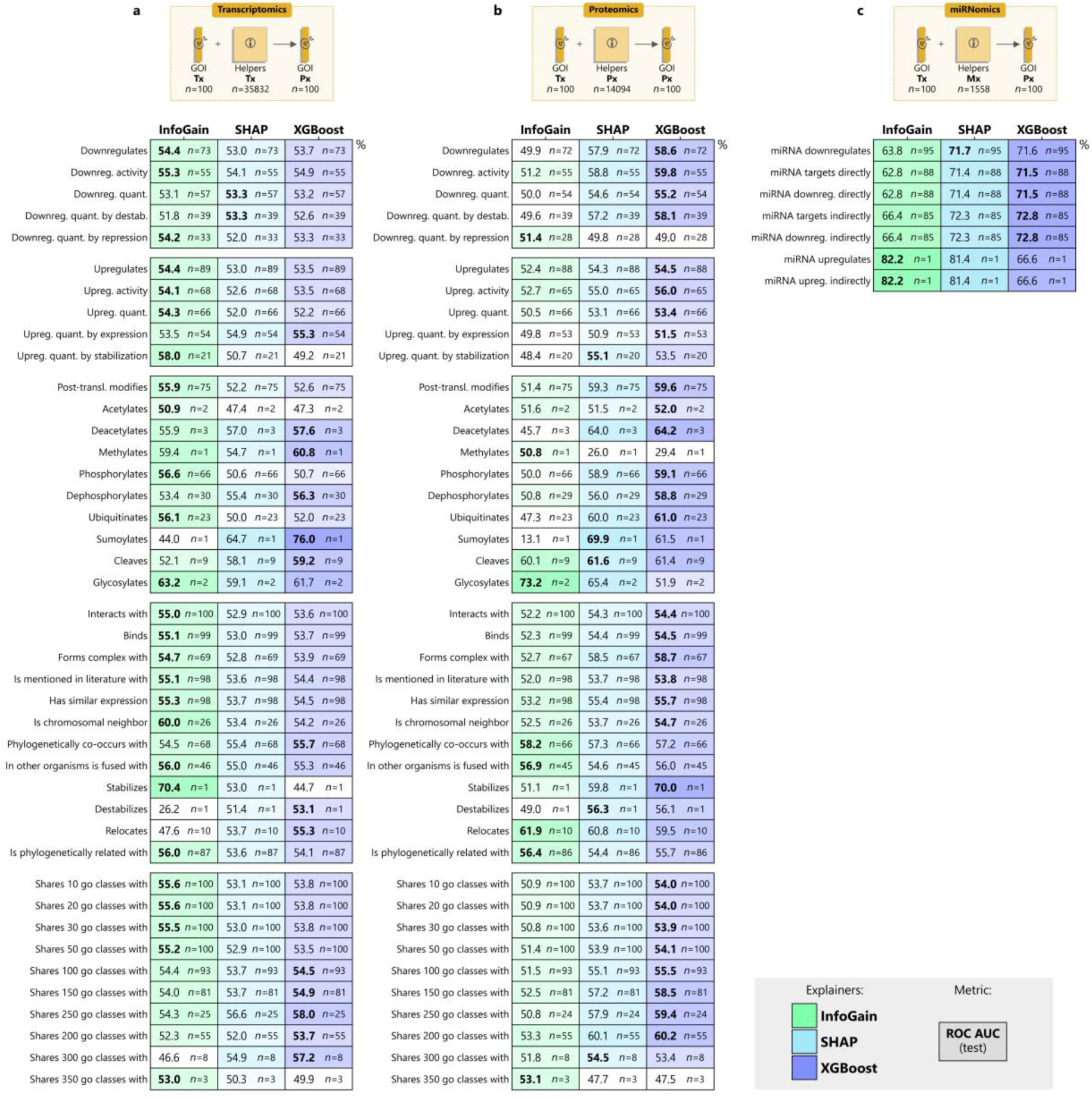
Proteomizer explainer ROC-AUC scores in the direct tasks (transcriptome to proteome), benchmarked on the gene-gene and microRNA-gene relations from the Biology Mega Graph. The plot shows the receiver operating characteristic area under the curve (ROC-AUC) scores, obtained by comparing the importance scores from three Proteomizer explainers, against the known gene-gene and microRNA-gene relations in scientific literature. The Proteomizer importance scores were generated from a model, that was tasked to predict the mass spectrometry-based proteome (Px) of 100 highly annotated genes of interest (GOIs), from their respective RNA sequencing-based transcriptome (Tx) (direct task), across 1,594 samples from 9 tissues of origin. Specifically, the importance score was determined for each of the helper genes (i.e., non-GOI genes) using three different explainers. (i) Information Gain (InfoGain, column 1) consists of training an independent eXtreme Gradient Boosting (XGBoost) model for each GOI-helper pair, where the GOI’s Tx column and the helper column were provided as the input, and the GOI’s Px as the target; then, taking the mean absolute error (MAE) across samples, with an inverted sign, as a score. (ii) XGBoost (column 2) consists of training an independent XGBoost model for each domain-helper pair, whereby the GOI’s Tx and all helpers in a domain—either whole Tx, whole Px, or whole miRNome (Mx)—were provided as the input, and GOI Px as the target; then, taking the XGBoost importance scores. (iii) SHapley Additive exPlanations (SHAP, column 3) is identical to (ii), but SHAP scores were used instead of the XGBoost importance ones. Finally, the importance scores from every explainer were compared against the known GOI-helper relations from literature, for each of 47 relation types, for each of the 100 GOIs. Such relations were sourced from the Biology Mega Graph (Supplementary Table 1), a knowledge graph which aggregates 320 million curated gene-gene and microRNA-gene relations from 11 biological repositories. The agreement between importance scores and literature was calculated in terms of ROC-AUC, in a direction-invariant way—e.g., the (*AKT*, *GSK3*) “phosphorylates” pair is treated as existing, if either AKT phosphorylates GSK3, or if GSK3 phosphorylates AKT. The top section of each column represents the task domain. (a) From the GOI’s Tx and each helper gene’s Tx, predicting the GOI’s Px. (b) From the GOI’s Tx and each helper gene’s Px, predicting the GOI’s Px. (c) From the GOI’s Tx and each helper miRNA’s Mx, predicting the GOI’s Px. The bottom section of each table expresses the ROC-AUC and sample numerosity for each of the 49 biological relation types, namely 42 gene-gene (a, b) and 7 miRNA-gene relations (c). The remaining terms were dropped due to insufficient number of examples. Numerosity (*n*) expresses how many of the 100 genes had at least one annotation of that corresponding relation type (e.g., if AKT phosphorylates some other proteins or is phosphorylated by some other proteins, then it will be counted as 1). The ROC-AUC of the best performing explainer for each task is shown in bold.

**Supplementary Figure 10.**
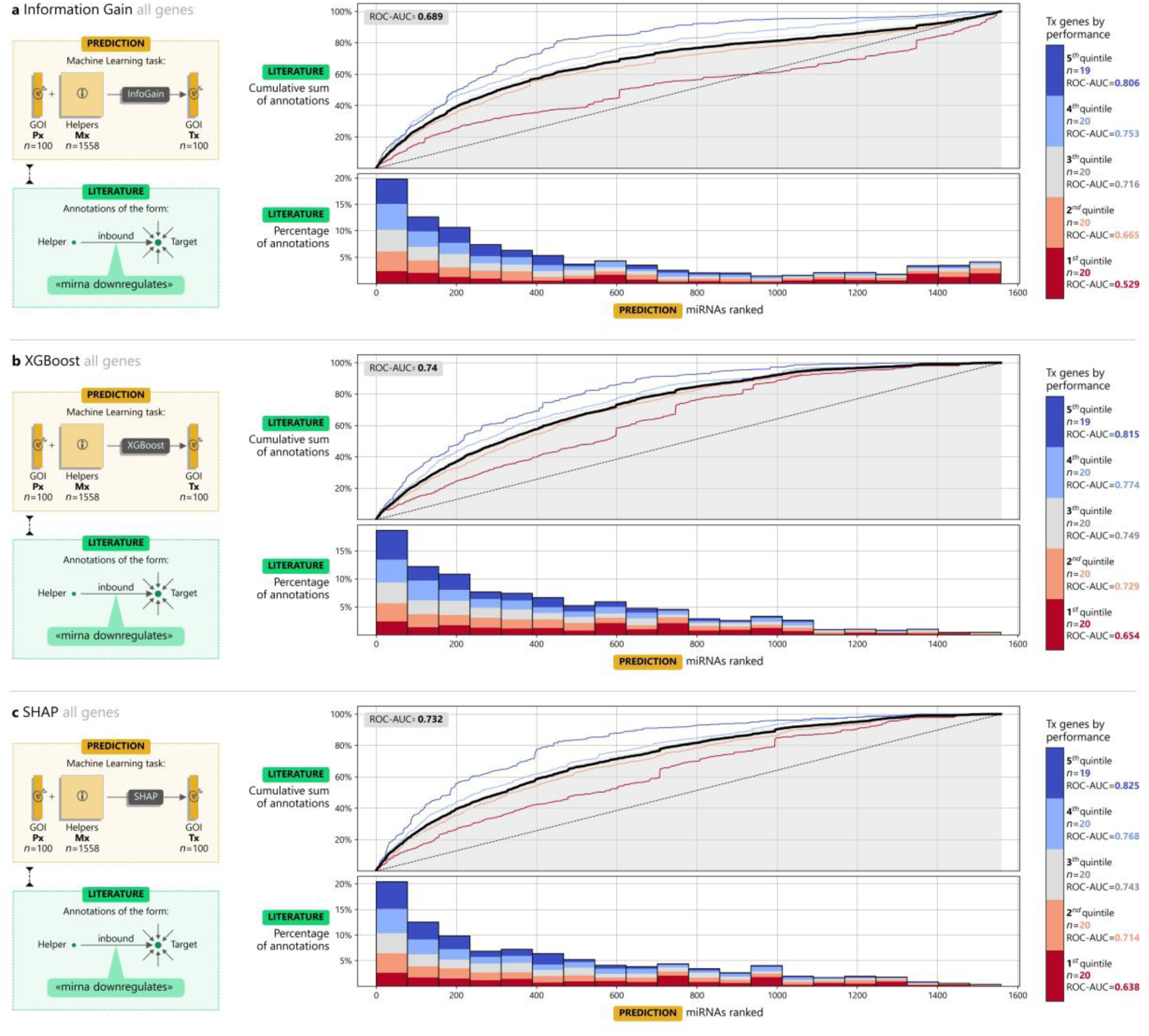
Proteomizer explainer cumulative gain curve in the reverse tasks (proteome to transcriptome), benchmarked against the “mirna downregulates” relation in the Biology Mega Graph. The plot shows the cumulative gain curve, obtained by comparing the importance scores from three Proteomizer explainers, against the known microRNA (miRNA) to gene downregulation relations in scientific literature. The Proteomizer importance scores were generated from a model, that was tasked to predict the RNA sequencing-based transcriptome (Tx) for 100 highly annotated genes of interest (GOIs), from their respective mass spectrometry-based proteome (Px) (reverse task), across 1,594 samples from 9 tissues of origin. Specifically, the importance score was determined for each of the helper genes (i.e., non-GOI genes) using three different explainers: Information Gain (InfoGain, row a), eXtreme Gradient Boosting (XGBoost, row b), SHapley Additive exPlanations (SHAP, row c)—see Figure 6 for details. Next, the importance scores were compared against the known miRNA-gene relations from literature. Such relations were sourced from the Biology Mega Graph (Supplementary Table 1) a knowledge graph which aggregates 320 million curated gene-gene and miRNA-gene relations from 11 biological repositories. Specifically, the relations marked as “miRNA downregulates” were used, which were sourced from TarBase. In each row in the figure, the left section represents the task (yellow), and the comparison with the Biology Mega Graph (green). The center section is a gain cumulative curve, which represents the average cumulative gain curve across the 100 GOIs: for each of the GOIs, the 1,558 miRNAs in the Proteomizer dataset were ranked, based on the predicted likelihood of them downregulating the GOI, according to Proteomizer explainer; next, a cumulative sum was computed, treating as 1.0 the miRNA-GOI pairs which existed in the “mirna downregulates” level of the Biology Mega Graph, and as 0.0 those that did not exist; the cumulative sum plots were averaged out across the 100 GOIs, and binned into a histogram; a color code was applied based on the average receiver operating characteristic area under the curve (ROC-AUC) score of each miRNA, to enable the reader to distinguish the cumulative gain curve of poorly predictable miRNAs (red) from that of highly predictable ones (blue). The right section shows the color code legend, and miRNA numerosity in each of the 5 ROC-AUC bins.

**Supplementary Figure 11.**
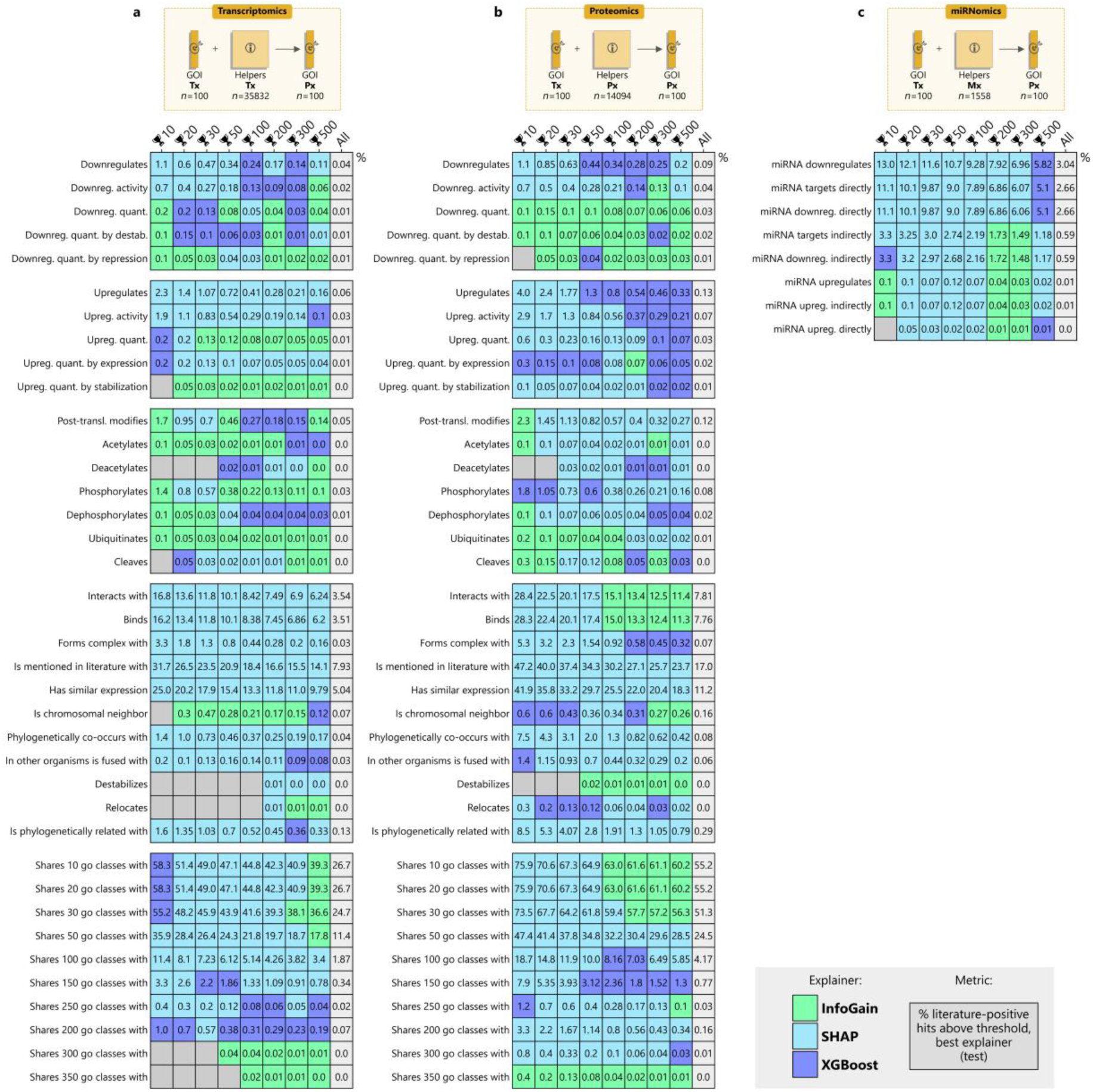
Proteomizer explainer top-hit positive fractions in the direct tasks (transcriptome to proteome), benchmarked against the gene-gene and microRNA-gene relations in the Biology Mega Graph. The plot shows the fraction of top hits by three Proteomizer explainers, that are also known in the current scientific literature. The Proteomizer importance scores were generated from a model, that was tasked to predict the mass spectrometry-based proteome (Px) of 100 highly annotated genes of interest (GOIs), from their respective RNA sequencing-based transcriptome (Tx) (direct task), across 1,594 samples from 9 tissues of origin. Specifically, the importance score was determined for each of the helper genes (i.e., non-GOI genes) using three different explainers: Information Gain (InfoGain, green), eXtreme Gradient Boosting (XGBoost, blue), SHapley Additive exPlanations (SHAP, cyan). Next, the importance scores were compared against the known GOI-helper relations from literature, for each of 47 relation types, for each of the 100 GOIs. Such relations were sourced from the Biology Mega Graph (Supplementary Table 1), a knowledge graph which aggregates 320 million curated gene-gene and miRNA-gene relations from 11 biological repositories. The agreement between the importance scores and literature were calculated in terms of top-hit positive fraction, in a direction-invariant way—e.g., the (*AKT*, *GSK3*) “phosphorylates” pair is treated as existing, if either AKT phosphorylates GSK3, or if GSK3 phosphorylates AKT. Essentially, for each of the GOIs, the explainer generated a raking of the helper genes; the ranking was cut at a variable threshold (🏆 *n*), starting from the top 5,000 and moving up to the top 10; this list of “predicted interactors” was compared with the list of known interactors from the Biology Mega Graph, and the percentage of genes in common between the two was counted (whereby 0% indicates that none of the top *n* hits are known interactors in a given biological modality, and 100% indicates that all of the top *n* genes are known interactors. The unthresholded percentage (All) is also displayed for reference, to highlight the enrichment observed among the top hits. The top section of each table represents the task domain. (a) From the GOI’s Tx and each helper gene’s Tx, predicting the GOI’s Px. (b) From the GOI’s Tx and each helper gene’s Px, predicting the GOI’s Px. (c) From the GOI’s Tx and each helper miRNA’s Mx, predicting the GOI’s Px. The bottom section of each table represents the top-hit positive fractions at varying thresholds. Specifically, the fraction was measured from the best-performing explainer, i.e., the one that yielded the highest fraction; the identity of such explainer varied on a case-by-case basis, and is expressed in a color code.

## Supplementary Tables

**Supplementary Table 1.** Structure of the Biology Mega Graph. The Biology Mega Graph is a knowledge graph, i.e., a network made up of multiple layers representing edges of different types. It is composed of 196,921 nodes, 193,384 of which represent human genes, and 3,537 represent human microRNAs (miRNAs); and 322,255,334 edges in 69 levels. It was assembled from 11 sources, namely: Gene Ontology (GO, number of edges *n* = 229,299,929), microT-miRBase (microT2, *n* = 41,677,849), Search Tool for the Retrieval of Interacting Genes/Proteins (STRING, *n* = 35,311,526), microT-MirGeneDB (microT1, *n* = 11,525,038), TarBase (*n* = 2,910,516), Signor (*n* = 455,412 for single genes, and *n* = 148,932 for protein complexes, cplx), ComplexPortal (CplxPortal, *n* = 533,136), NCBI Gene (*n* = 193,384), PhosphoSitePlus (PhosphoSite, *n* = 42,846), miRBase (*n* = 4,801), human DEPhOsphorylation Database (DEPOD, *n* = 3,033). The graph is also available for *M. musculus*.

| # | Modality | Numerosity | # | Modality | Numerosity |
| --- | --- | --- | --- | --- | --- |
| 1 | ADP-ribosylates | Signor $n = 12$ | 32 | mirna downregulates | TarBase $n = 727,058$ |
| 2 | acetylates | Signor $n = 2,857$ | 33 | mirna downregulates directly | TarBase $n = 587,834$ |
| 3 | activates | Signor $n = 20$ | 34 | mirna downregulates indirectly | TarBase $n = 139,224$ |
| 4 | binds | Signor-cplx $n = 49,644$<br>Signor $n = 129,009$<br>CplxPortal $n = 177,712$<br>STRING $n = 6,156,420$ | 35 | mirna targets | TarBase $n = 727,629$<br>microT1 $n = 11,525,038$<br>microT2 $n = 41,677,849$ |
| 5 | carboxylates | Signor $n = 42$ | 36 | mirna targets directly | TarBase $n = 587,909$ |
| 6 | cleaves | Signor $n = 682$ | 37 | mirna targets indirectly | TarBase $n = 139,720$ |
| 7 | deacetylates | Signor $n = 143$ | 38 | mirna upregulates | TarBase $n = 571$ |
| 8 | deglycosylates | Signor $n = 7$ | 39 | mirna upregulates directly | TarBase $n = 75$ |
| 9 | demethylates | Signor $n = 457$ | 40 | mirna upregulates indirectly | TarBase $n = 496$ |
| 10 | dephosphorylates | DEPOD $n = 1,011$<br>Signor $n = 1,768$ | 41 | neddylates | Signor $n = 20$ |
| 11 | destabilizes | Signor $n = 69$ | 42 | nitrosylates | Signor $n = 1$ |
| 12 | desumoylates | Signor $n = 6$ | 43 | oxidates | Signor $n = 1$ |
| 13 | deubiquitinates | Signor $n = 156$ | 44 | palmitoylates | Signor $n = 15$ |
| 14 | downregulates | Signor $n = 21,644$ | 45 | phosphorylates | PhosphoSite $n = 14,282$<br>Signor $n = 20,743$ |
| 15 | downregulates activity | Signor $n = 11,396$ | 46 | phylogenetically co-occurs with | STRING $n = 219,906$ |
| 16 | downregulates quantity | Signor $n = 5,684$ | 47 | post-translationally modifies | DEPOD $n = 1,011$<br>PhosphoSite $n = 14,282$<br>Signor $n = 31,058$ |
| 17 | downregulates quantity by destabilization | Signor $n = 3,742$ | 48 | relocates | Signor $n = 1,681$ |
| 18 | downregulates quantity by repression | Signor $n = 1,529$ | 49 | shares 1 go class with | GO $n = 56,538,421$ |
| 19 | forms complex with | Signor-cplx $n = 49,644$<br>Signor $n = 129,009$<br>Signor-cplx $n = 177,712$ | 50 | shares 10 go classes with | GO $n = 56,538,421$ |
| 20 | glycosylates | Signor $n = 67$ | 51 | shares 100 go classes with | GO $n = 543,148$ |
| 21 | has similar expression pattern as | StringDB $n = 9,475,554$ | 52 | shares 150 go classes with | GO $n = 52,880$ |
| 22 | hydroxylates | Signor $n = 65$ | 53 | shares 20 go classes with | GO $n = 56,538,421$ |
| 23 | in other organisms is fused with | STRING $n = 59,452$ | 54 | shares 200 go classes with | GO $n = 7,228$ |
| 24 | inhibits | Signor $n = 2$ | 55 | shares 250 go classes with | GO $n = 1,276$ |
| 25 | interacts with | DEPOD $n = 1,011$<br>PhosphoSite $n = 14,282$<br>Signor-Cplx $n = 49,644$<br>Signor $n = 170,289$<br>CplxPortal $n = 177,712$<br>STRING $n = 6,156,420$ | 56 | shares 30 go classes with | GO $n = 48,136,204$ |
| 26 | is chromosomically neighborhood | STRING $n = 541,066$ | 57 | shares 300 go classes with | GO $n = 292$ |
| 27 | is identical | miRBase $n = 4,801$<br>NCBI Gene $n = 193,384$ | 58 | shares 350 go classes with | GO $n = 44$ |
| 28 | is mentioned in literature with | STRING $n = 11,882,284$ | 59 | shares 400 go classes with | GO $n = 4$ |
| 29 | is phylogenetically related with | STRING $n = 820,424$ | 60 | shares 50 go classes with | GO $n = 10,943,590$ |
| 30 | lipidates | Signor $n = 6$ | 61 | stabilizes | Signor $n = 129$ |
| 31 | methylates | Signor $n = 460$ | 62 | sumoylates | Signor $n = 65$ |
| | | | 63 | tyrosinates | Signor $n = 25$ |
| | | | 64 | ubiquitinates | Signor $n = 3,450$ |
| | | | 65 | upregulates | Signor $n = 35,486$ |
| | | | 66 | upregulates activity | Signor $n = 21,430$ |
| | | | 67 | upregulates quantity | Signor $n = 6,482$ |
| | | | 68 | upregulates quantity by expression | Signor $n = 3,414$ |
| | | | 69 | upregulates quantity by stabilization | Signor $n = 1,223$ |

**Supplementary Table 2.** Standardization of the tissue or organ of origin, in the TCGA dataset. The table displays how the “tissue_or_organ_of_origin” column was standardised, in the sample metadata from The Cancer Genome Atlas (TCGA). Tissues with similar (or identical) histology were aggregated together (e.g., “brain”, “brain stem”, “cerebellum”,…). In contrast, categories were left purposefully broad to guarantee that at least two separate biologies would be present, in preparation for differential expression analysis across sub-clusters.

| Original | Standardised | Original | Standardised | Original | Standardised |
| --- | --- | --- | --- | --- | --- |
| abdomen | - | head, face or neck | head neck | overlapping lesion of retroperitoneum and peritoneum | - |
| adrenal gland | adrenal | head of pancreas | pancreas | palate | head neck |
| ampulla of vater | bile ducts | heart | heart | pancreas | pancreas |
| anal canal | anus | hematopoietic system | blood | pancreatic ductal adenocarcinoma | pancreas |
| anterior floor of mouth | head neck | hepatic flexure of colon | colorectal | parietal lobe | brain |
| anterior mediastinum | - | hypopharynx | head neck | pelvic bones, sacrum, coccyx and associated joints | bone |
| anterior wall of bladder | bladder | intestinal tract | colorectal | pelvic lymph nodes | lymphatic |
| anus | anus | intra-abdominal lymph nodes | lymphatic | pelvis | bone |
| aortic body and other paraganglia | - | intrahepatic bile duct | bile ducts | peripheral nerves and autonomic nervous system of thorax | - |
| ascending colon | colorectal | intrathoracic lymph nodes | lymphatic | peripheral nerves and autonomic nervous system of trunk | - |
| base of tongue | head neck | isthmus uteri | uterus | peripheral nerves and autonomic nervous system of upper limb and shoulder | - |
| bladder | bladder | jejunum | small intestine | peritoneum | peritoneum |
| bladder neck | bladder | kidney | kidney | pharynx | head neck |
| blood | blood | larynx | head neck | pleura | pleura |
| body of pancreas | pancreas | lateral wall of bladder | bladder | posterior mediastinum | - |
| body of stomach | stomach | lesser curvature of stomach | stomach | posterior wall of bladder | bladder |
| bone | bone | lip | head neck | posterior wall of oropharynx | head neck |
| bone marrow | blood | liver | liver | prostate gland | prostate |
| bones of skull and face and associated joints | bone | long bones of lower limb and associated joints | bone | pylorus | stomach |
| border of tongue | head neck | long bones of upper limb, scapula and associated joints | bone | rectosigmoid junction | colorectal |
| brain | brain | lower gum | head neck | rectum | colorectal |
| brain stem | brain | lower lobe, lung | lung | retromolar area | head neck |
| breast | breast | lower third of esophagus | esophagus | retroperitoneum | - |
| cardia | stomach | lower-inner quadrant of breast | breast | short bones of lower limb and associated joints | bone |
| cecum | colorectal | lower-outer quadrant of breast | breast | sigmoid colon | colorectal |
| cerebellum | brain | lung | lung | skin | skin |
| cerebrum | brain | lung adenocarcinoma | lung | skin of lower limb and hip | skin |
| cervix uteri | cervix | lung squamous cell carcinoma | lung | skin of upper limb and shoulder | skin |
| cheek mucosa | head neck | lymph node | lymphatic | small intestine | small intestine |
| choroid | choroid | lymph nodes of axilla or arm | lymphatic | specified parts of peritoneum | peritoneum |
| ciliary body | - | lymph nodes of head, face and neck | lymphatic | spermatic cord | - |
| clear cell renal cell carcinoma | kidney | lymph nodes of inguinal region or leg | lymphatic | spinal meninges | - |
| colon | colorectal | main bronchus | lung | splenic flexure of colon | colorectal |
| connective, subcutaneous and other soft tissues | connective | mandible | bone | stomach | stomach |
| connective, subcutaneous and other soft tissues of abdomen | connective | mediastinum | - | submandibular gland | - |
| connective, subcutaneous and other soft tissues of head, face, and neck | connective | medulla of adrenal gland | adrenal | supraglottis | - |
| connective, subcutaneous and other soft tissues of lower limb and hip | connective | middle lobe, lung | lung | tail of pancreas | pancreas |
| connective, subcutaneous and other soft tissues of pelvis | connective | middle third of esophagus | esophagus | temporal lobe | brain |
| connective, subcutaneous and other soft tissues of thorax | connective | mouth | head neck | testis | testis |
| connective, subcutaneous and other soft tissues of trunk | connective | myometrium | uterus | thoracic esophagus | esophagus |
| connective, subcutaneous and other soft tissues of upper limb and shoulder | connective | nervous system | - | thorax | - |
| corpus uteri | uterus | non-clear cell renal cell carcinoma | kidney | thymus | thymus |
| cortex of adrenal gland | adrenal | occipital lobe | brain | thyroid gland | thyroid |
| descending colon | colorectal | orbit | - | tongue | head neck |
| dome of bladder | bladder | oropharynx | head neck | tonsil | head neck |
| endometrium | uterus | other ill-defined sites | - | transverse colon | colorectal |
| esophagus | esophagus | ovarian serous cystadenocarcinoma | ovary | trigone of bladder | bladder |
| external upper lip | head neck | ovary | ovary | unknown primary site | - |
| extrahepatic bile duct | bile ducts | overlapping lesion of breast | breast | upper gum | head neck |
| fallopian tube | fallopian tube | overlapping lesion of colon | - | upper lobe, lung | lung |
| floor of mouth | head neck | overlapping lesion of connective, subcutaneous and other soft tissues | - | upper third of esophagus | esophagus |
| frontal lobe | brain | overlapping lesion of eye and adnexa | - | upper-inner quadrant of breast | breast |
| fundus of stomach | stomach | overlapping lesion of ill-defined sites | - | upper-outer quadrant of breast | breast |
| fundus uteri | uterus | overlapping lesion of lip, oral cavity and pharynx | head neck | ureteric orifice | bladder |
| gallbladder | bile ducts | overlapping lesion of lung | - | urethra | urethra |
| gastric antrum | stomach | overlapping lesion of pancreas | - | uterine corpus endometrial carcinoma | uterus |
| gastrointestinal tract | - |  |  | uterus | uterus |
| glioblastoma | brain |  |  | ventral surface of tongue | head neck |
| gum | head neck |  |  |  |  |
| hard palate | head neck |  |  |  |  |

